# Autopolyploidy exacerbates dominance masking under negative frequency-dependent selection : evidence from sporophytic self-incompatibility in *Arabidopsis arenosa* and *A. lyrata*

**DOI:** 10.1101/2025.06.11.659117

**Authors:** Xavier Vekemans, Uliana Kolesnikova, Jakub Vlcek, Mathieu Genete, Adrien Dufour, Rowan Welti, Céline Poux, Vincent Castric, Filip Kolar, Polina Novikova

## Abstract

Polyploidy is a widespread phenomenon in flowering plants, with up to 30% of extant species being recent polyploids. Whether and how polyploidy modulates the action of natural selection remains debated, and the particular case of balancing selection has been poorly explored. This study investigates the impact of autopolyploidy on sporophytic self-incompatibility in plants, a striking example of a genetic system evolving under a special form of balancing selection (strong negative frequency-dependent selection). Numerical simulations reveal that under strict codominance, the number of S-alleles maintained in tetraploid populations is expected to double as compared to diploid populations. However, under a model with strict hierarchical dominance among alleles, the number of S-alleles increases only slightly, but gene diversity and observed heterozygosity are substantially lower in tetraploids as compared to diploids, due to enhanced dominance masking effects. Empirical data on *Arabidopsis arenosa* and *A. lyrata* confirm the latter predictions, showing similar levels of allelic diversity but dramatically lower observed and expected heterozygosity at the self-incompatibility locus in tetraploids compared to diploids. The study highlights the significant impact of autopolyploidy on patterns of diversity at the self-incompatibility locus, emphasizing the increased dominance effect in tetraploids. The results also allow us to reject a scenario of strong founder effects associated with evolution of the polyploid lineages.

## Introduction

The evolutionary transition from diploidy to polyploidy by whole-genome duplication (WGD) is widespread among flowering plants. About 35% of extant species might be recent polyploids (Wood et al. 2009), resulting from WGD either within one lineage (autopolyploids) or between different lineages (allopolyploids). The impact of polyploidy on the action of natural selection has spurred considerable interest, dating back to the early days of the field of population genetics (reviewed by Bever & Felber 1992), to establish clear theoretical expectations and test them in empirical studies.

How deleterious mutations are expected to accumulate in tetraploids as compared to diploids (i.e. the mutation load) can be challenging to predict because of conflicting effects. For instance, autotetraploids have a higher effective population size and thus potentially more efficient purifying selection, but at the same time fully or partially recessive deleterious alleles are more likely to be masked by a functional copy, and their elimination by purging under partial selfing was shown to proceed differently in polyploids as compared to diploids (Lande & Schemske 1995; Ronfort 1999; Otto & Whitton 2000). In addition, the population mutation rate is higher in autotetraploids, due to increased gene copies per individual. Overall, at mutation-selection equilibrium under full outcrossing, a higher mutation load of partially recessive deleterious alleles is expected in polyploid than in diploid populations (Ronfort 1999; Otto & Whitton 2000; Vlcek et al. 2025). However, this general prediction is only valid over the long term, and in the transient stage after an autopolyploid is formed the instantaneous masking effect is instead expected to temporarily reduce the mutation load until new deleterious alleles start accumulating again (Otto & Whitton 2000). These predictions are largely confirmed by a series of studies comparing autotetraploid with diploid reference populations of *Arabidopsis arenosa.* Specifically, tetraploids tended to show a higher ratio of non-synonymous to synonymous polymorphisms in protein-coding genes (Monnahan et al. 2019; Vlcek et al. 2025), more common insertions of transposable elements within exons (Baduel et al. 2019) and a higher prevalence of large structural variants (Vlcek et al. 2025) as compared to diploids. Hence, in these populations, the increased mutational input and increased effect of masking tend to dominate over the increased effective population size in polyploids. The fate of beneficial mutations may also differ between diploid and tetraploids, and theoretical investigations also identified potentially complex interactions between the masking effect (Hill 1971) and the pace of adaptive evolution (Otto & Whitton 2000; Selmecki et al. 2015; Griswold 2021). Overall, beneficial alleles that are at least partially dominant (i.e have a dominance coefficient *h*>0) are expected to fix more readily in autopolyploids as compared to diploids, resulting in higher rates of fitness increase (Otto & Whitton 2000). Again, these predictions were confirmed by empirical studies using population genomics approaches in *A. arenosa*, with significantly higher proportions of non-synonymous polymorphisms fixed by positive selection (Monnahan et al. 2019) and higher numbers of putatively adaptive new Copia-elements transpositions reaching high frequency in autotetraploid as compared to diploid populations (Baduel et al. 2019).

Balancing selection is a form of natural selection maintaining, rather than depleting, genetic polymorphisms. In contrast to directional (positive and negative) selection, the effect of autopolyploidy on genetic systems subject to balancing selection has received much less attention (but see Hill 1971). Yet, the increased effective population size and the more prevalent masking of recessive alleles in polyploids may affect the distribution of allele frequencies and allelic diversity at loci subject to balancing selection in non-trivial manners. In particular, in systems subject to long-term multiallelic balancing selection, the strength of selection varies with the number of balanced allelic lines (Yokoyama & Nei 1979), so the possible loss of alleles associated with founder effects upon establishment of the polyploid (Layman & Busch 2018) would be expected to affect these systems in ways that have not been characterized yet.

Multiallelic self-incompatibility (SI) systems are iconic examples of long-term negative frequency-dependent selection (Lawrence 2000; Castric & Vekemans 2004; Durand et al. 2020). Two main types of SI systems occur in flowering plants. In sporophytic SI (SSI), the pollen compatibility phenotype is determined by the diploid paternal genotype at the SI locus (S-locus). In gametophytic SI (GSI), the pollen compatibility phenotype is determined by the haploid pollen genotype itself. SSI systems, as well as some GSI systems (Papaveraceae: Thomas & Frankling-Tong 2004; Prunus: Vieira et al. 2010) function as self-recognition systems, in which the pollen and pistil proteins produced at the S-locus from the same haplotype recognize each other and trigger self-pollen rejection (Fuji et al. 2016; Vekemans & Castric 2021). In these self-recognition systems, the self-pollen rejection function was found to be maintained in extent autopolyploid populations or in synthetic neopolyploids (e.g. in Brassicaceae: Mable et al. 2004, Duan et al. 2024; in Prunus: Hauck et al. 2006). In contrast, the most widely distributed GSI system, known as the S-RNase system (Ramanauskas et al. 2024), functions as a non-self recognition system in which the toxic pistil protein (a ribonuclease) is taken up by the pollen tubes and degraded by a dozen of pollen antitoxins encoded at the S-locus, which can detoxify collaboratively all S-alleles of the population except the one present in the same haplotype (Kubo et al. 2010; Fuji et al. 2016). Polyploidy causes an immediate loss of the self-pollen rejection function in the S-RNase system (Fuji et al. 2016), leading to self-compatibility. This is due to the fact that diploid pollen tubes that are heterozygous at the S-locus will contain all the battery of detoxifying genes necessary to inactivate both S-RNases expressed by their own pistil (Kubo et al. 2010), explaining a commonly observed phenomenon named “competitive interaction” (Tsukamoto et al. 2005). This functional property leads to a strong association between polyploidy and loss of SI in plant families with a S-RNase system (Robertson et al. 2011), which is not found in plants with SSI (Duan et al. 2024).

SSI systems are particularly interesting in the context of polyploidy because they often exhibit complex patterns of dominance relationships among alleles at the S-locus (Schierup et al. 1997), which could generate qualitative differences between diploids and tetraploids on the outcome of negative frequency-dependent selection. However, detailed population genetic predictions about the impact of polyploidy on SSI systems are currently lacking, especially considering dominance effects at the S-locus. In addition, plant families with SSI, such as the Brassicaceae and Asteraceae, comprise numerous independently emerged polyploid lineages (Huang et al. 2016; Kolar et al. 2017; Mandakova et al. 2017), raising the question of the coevolution of SI and polyploidy. In Brassicaceae, the general pattern seems to be that autopolyploids derived from diploid self-incompatible progenitors maintain a strictly outcrossing mating system with functional SI, while the SI system is systematically lost in allopolyploids, often followed by evolution of high selfing (reviewed in Novikova et al. 2023). Interestingly, several self-incompatible Brassicaceae species, e.g. *Arabidopsis arenosa* and *A. lyrata*, are composed of a combination of diploid and autotetraploid populations (Mable et al. 2004; Schmickl et al. 2012), which provides unique opportunities to investigate the impact of polyploidy on patterns of allele diversity and allelic frequency distributions at the S-locus. In *A. arenosa* and *A. lyrata*, autotetraploid populations have independently established in nature from conspecific diploid lineage(s), as documented using population genomic approaches (Marburger et al. 2019; Monnahan et al. 2019; Bohutinska et al. 2024; Scott et al. 2024). Both species share the same SSI system that has been well characterized in *A. lyrata* (e.g. Kusaba et al. 2001; Schierup et al. 2001; Prigoda et al. 2005; Kolesnikova et al. 2023). Patterns of allelic variation at the S-locus have been investigated in limited samples of tetraploid populations from both species, and were consistent in terms of patterns of polymorphism with a fully functional SI system (Mable et al. 2004; Mable et al. 2018; Scott et al. 2024).

In this study, we combined theoretical and empirical approaches to investigate the effect of autopolyploidy on patterns of variation at the locus controlling SSI, a locus subject to strong negative frequency-dependent selection. We first used numerical simulations to determine the expected effect of polyploidy on S-locus diversity under a range of SSI models (Schierup et al. 1997). We then analyzed individual short-read resequencing data to evaluate S-locus diversity in diploid (N=207) and tetraploid (N=365) populations of *Arabidopsis arenosa* and diploid (N=417) and tetraploid (N=138) populations of *A. lyrata* We found similar levels of allelic diversity at the S-locus in tetraploid as in diploid cytotypes, suggesting the absence of major founder events during establishment of the autotetraploid populations. However, in line with our simulation results for sporophytic SI models with dominance, and in stark contrast to the expectation for a neutral locus, we found that tetraploids exhibited dramatically lower observed and expected heterozygosity at the S-locus as compared to diploids. Hence, the shift from diploidy to tetraploidy has a major effect on patterns of diversity at the S-locus. This highlights that, under the particular form of negative frequency-dependent selection under which this S-locus evolves, the impact of the increased dominance effect incurred by tetraploids largely outperforms that of their increased effective population size.

## Results

### Numerical simulations of sporophytic SI in diploid and tetraploid populations

We performed numerical simulations in finite populations to compare the effect of polyploidy on S-locus diversity between symmetric (with strict codominance among alleles in pollen and pistil, referred to as SSI-COD) and asymmetric (with strict hierarchical dominance in pollen and pistil, called SSI-DOM) models of sporophytic SI. We used the sporophytic SI simulation framework of Schierup et al. (1997), modified to implement tetraploid populations, and used the same simulation parameters for comparison. Under the model with full co-dominance (SSI-COD), we observed that the number of S-alleles maintained in tetraploid populations is expected to approximately double as compared to that in diploid populations (from 8.6 to 17.5 with *N*=100, and from 15.7 to 31.5 with *N*=200, Table 1). Gene diversity increased with tetraploidy, but only slightly (from 0.87 to 0.94 with *N*=100, and from 0.92 to 0.96 with *N*=200, Table 1). Note that S-locus homozygotes cannot be formed under this model due to the strict codominance of S-alleles (Schierup et al. 1997), such that the observed heterozygosity was 100% in both diploid and tetraploid populations. Results under the model with dominance interactions between S-alleles (SSI-DOM) were strikingly different. Under this model, where natural selection acts asymmetrically on dominant vs recessive S-alleles, the number of S-alleles within tetraploid populations was only slightly increased as compared to diploids (from 4.49 to 5.07 with *N*=100, and from 9.28 to 12.20 with *N*=200, Table 1). However, both gene diversity (*H_e_*, probability that a randomly chosen pair of gene copies sampled from the overall cytotype gene pool corresponds to different S-alleles) and observed heterozygosity (*H*_o_, same but sampled within one diploid or tetraploid individual) were substantially lower (15-30% lower) in tetraploid as compared to diploid populations (*e.g.* with *N*=100, *H*_e_ decreased from 0.65 to 0.49 and *H*_o_ decreased from 0.73 to 0.51 Table 1). A striking prediction is that the distribution of S-allele frequencies are expected to differ strongly between diploid and tetraploid populations (Fig. 1). Specifically, in our simulations the most recessive S-allele reached frequencies more than ten-fold higher than the most dominant S-allele in tetraploid populations, whereas in diploids the asymmetry was less marked, with at most a four-fold higher frequency of the most recessive as compared to the most dominant S-allele (Table 1). Hence, our results clearly reveal that polyploidy is expected to exacerbate the effect of the asymmetrical negative frequency-dependent selection on the different S-alleles, resulting in sharply different distributions of S-alleles in natural populations. The shift from diploidy to polypoidy had an especially strong impact on the frequency of the most recessive S-allele (with N=100, Fig. 1A) or the two most recessive S-alleles (with N=200, Fig. 1B). This pattern was confirmed under deterministic simulations with a fixed number of S-alleles, although in this case only the most recessive S-allele reached higher frequencies in tetraploid as compared to diploid populations, and the magnitude of the difference was slightly larger than in finite populations (Fig. 1 C and D). Strikingly, when 12 alleles are segregating, the most recessive allele is expected to reach a population frequency of 0.57 in tetraploids (*versus* only 0.31 in diploids), meaning that a single S-allele comprises more than half of the gene copies at this locus (Fig. 1D), illustrating the sharply asymmetrical effect that negative frequency-dependent selection is expected to have on the different S-alleles according to whether they are high or low along the dominance hierarchy. This can be interpreted as the result of a strong increase of the dominance effect in polyploids, which in SSI systems leads to asymmetrical allele frequencies at equilibrium (Sampson 1974; Schierup et al. 1997).

**Fig. 1.**
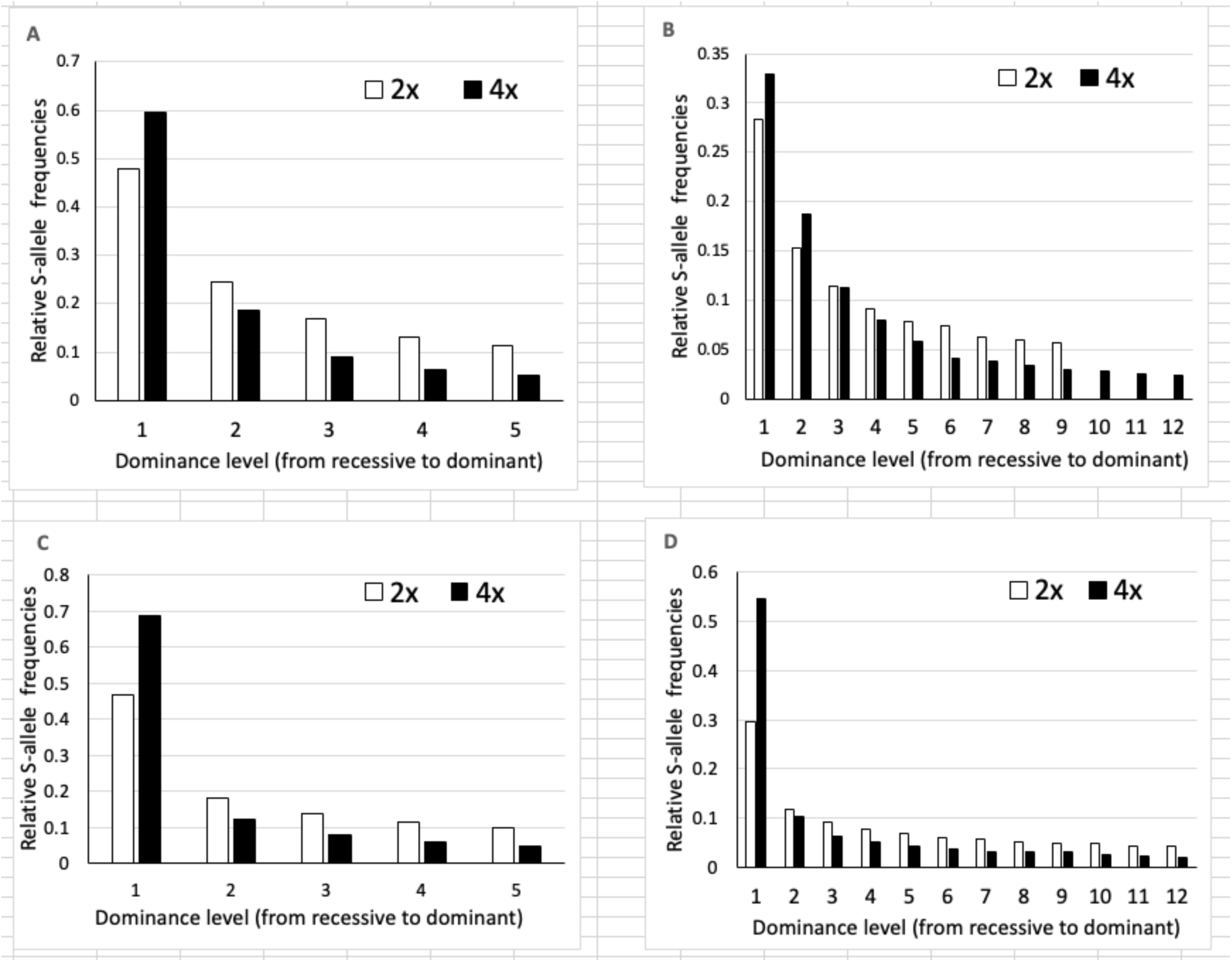
Expected relative frequencies of S-alleles as a function of allele dominance obtained by numerical simulations of the SSI-DOM model in diploid (white bars) and tetraploid (black bars) populations. Simulation of finite populations with *N*=100 and **μ**=1.10^-4^ (panel A) and *N*=200 and **μ**=5.10^-4^ (panel B). Deterministic simulations (*N*=7000, **μ**=0) with a number of S-alleles fixed to 5 (panel C) and 12 (panel D).

**Table 1.** Results of simulations of sporophytic SI in finite diploid (2x) and tetraploid (4x) populations with linear hierarchical dominance in both pollen and pistil. Mean values are given over 1000 replicate simulations ± standard deviation. *N* is the population size and *μ* is the mutation rate to new S-alleles. The statistics reported are the number of segregating S-alleles, Nei’s gene diversity (*H*_e_), observed heterozygosity computed as the gametic heterozygosity of an individual (*H*_o_, Moody et al. 1993), and for the model with dominance, the mean frequencies of the most recessive or most dominant alleles and their ratio.

| Ploidy | Number of S-alleles | Gene diversity ($H_e$) | Observed heterozygosity ($H_o$) | Freq. most recessive allele | Freq. most dominant allele | ratio freq. most recess. /most dominant |
| --- | --- | --- | --- | --- | --- | --- |
| $N=100$ $\mu=0.0001$ | | | | | | |
| *SSI with strict codominance (SSI-COD)* |  |  |  |  |  |  |
| 2x | 8.61 ± 0.98 | 0.870 ± 0.012 | 1.0 ± 0.0 | | | |
| 4x | 17.47 ± 1.33 | 0.936 ± 0.004 | 1.0 ± 0.0 | | | |
| *SSI with pollen and pistil dominance (SSI-DOM)* |  |  |  |  |  |  |
| 2x | 4.49 ± 0.81 | 0.650 ± 0.065 | 0.725 ± 0.069 | 0.479 ± 0.130 | 0.114 ± 0.043 | 4.2 |
| 4x | 5.07 ± 0.94 | 0.488 ± 0.088 | 0.509 ± 0.088 | 0.600 ± 0.232 | 0.053 ± 0.022 | 11.2 |
| $N=200$ $\mu=0.0005$ | | | | | | |
| *SSI with strict codominance (SSI-COD)* |  |  |  |  |  |  |
| 2x | 15.74 ± 9.99 | 0.920 ± 0.007 | 1.0 ± 0.0 | | | |
| 4x | 31.5 ± 2.60 | 0.960 ± 0.003 | 1.0 ± 0.0 | | | |
| *SSI with pollen and pistil dominance (SSI-DOM)* |  |  |  |  |  |  |
| 2x | 9.28 ± 1.67 | 0.7880 ± 0.049 | 0.832 ± 0.051 | 0.283 ± 0.158 | 0.056 ± 0.035 | 5.0 |
| 4x | 12.20 ± 2.14 | 0.667 ± 0.097 | 0.678 ± 0.097 | 0.329 ± 0.265 | 0.024 ± 0.016 | 14.0 |

### S-locus genotypes in natural diploid and tetraploid A. arenosa and A. lyrata populations

Next we applied the NGSgenotyp pipeline (Genete et al. 2020) on short-read resequencing data to obtain the genotype at the S-locus female specificity gene (*SRK*) of diploid and tetraploid *A. arenosa* and *A. lyrata* individuals. We used a reference database for *SRK* that includes known sequences from *A. halleri*, *A. lyrata* and *Capsella grandiflora* (Duan et al. 2024), and a limited dataset of partial *SRK* sequences of *A. arenosa* (Ruiz-Duarte 2012; Mable et al. 2018). Using the *de novo* assembly module included in NGSgenotyp, we were able to retrieve full exon 1 sequences of most *A. arenosa SRK* alleles detected in this study (Table S3) as well as of new *A. lyrata* alleles (Table S4),. With the genotyping module of NGSgenotyp, we were then able to fully resolve the 4x genotype of tetraploid individuals, using variations of the average read depth to evaluate precisely the number of copies of a given S-allele within the genotype of each individual (*e.g.* distinguishing S_1_S_1_S_1_S_2_, S_1_S_1_S_2_S_2_ and S_1_S_2_S_2_S_2_ genotypes, where the S_1_ and S_2_ alleles are present but in different copy numbers). With this strategy, we obtained the full genotypes of *N*=207 diploid *A. arenosa* and *N*=417 diploid *A. lyrata* individuals, and the full genotypes of *N*=356 tetraploid *A. arenosa* and *N*=138 tetraploid *A. lyrata* individuals. For nine *A. arenosa* (2.5%) and five *A. lyrata* (3.6%) tetraploid individuals, only one gene copy out of four was missing 9. Over the whole dataset, we identified a total of 83 S-alleles in *A. arenosa* (Table S3), with 56 alleles newly obtained in this study. Three additional sequences that are very similar to *SRK* sequences but are not linked to the S-locus (referred to as Aly13-2, Aly13-7 and Ah08) in *A. halleri* and *A. lyrata* (Charlesworth et al. 2003; Prigoda et al. 2005; Castric & Vekemans 2007) were also present. These sequences were systematically detected in individuals whose full genotype had already been determined with other *SRK* sequences, so we concluded that these sequences are also paralogs in *A. arenosa*, and we excluded them from further analysis. In *A. lyrata,* we identified a total of 66 S-alleles (Table S4), with ten alleles newly obtained in this study. Prigoda et al. (2005) classified S-alleles of *A. lyrata* in four phylogenetic groups, *i.e.* A1, B, A2 and A3 (renamed classes I to IV, respectively, following Goubet et al. 2012), which were shown to be associated with increasing levels of allelic dominance (from the most recessive class I to the most dominant class IV). Among the 83 S-alleles identified in *A. arenosa*, one belongs to class I, 20 to class II, 20 to class III, and 42 to class IV (Table S3), whereas for the 66 S-alleles of *A. lyrata*, one belongs to class I, 9 to class II, 15 to class III, and 41 to class IV (Table S4).

### Patterns of S-locus polymorphism in diploid versus tetraploid populations

Out of the total 83 S-alleles observed in *A. arenosa,* 81 were detected in diploid and 80 in tetraploid individuals. Similarly, out of the total 66 S-alleles observed in *A. lyrata*, 61 were detected in diploid and 61 in tetraploid individuals (Table 2). Hence, taken at face value the diploid and tetraploid cytotype gene pools share most of their S-alleles within each species. However, in *A. arenosa* the number of individuals investigated in tetraploids (N=365) is higher than in diploids (N=207), while the reverse is true in *A. lyrata* (417 diploids and 138 tetraploids). Applying a rarefaction algorithm to control for differences in sample sizes, we estimated the average allelic richness among 20 gene copies to be 15.96 in diploids and only 9.41 in tetraploids in *A. arenosa* (13.88 and 10.36, respectively in diploids and tetraploids of *A. lyrata*). Hence, allelic richness is actually substantially reduced in tetraploids (Table 2). This is confirmed by estimates of gene diversity (*H_e_*) and of observed heterozygosity (*H_o_*), which showed much higher values in diploids (*H_e_* = 0.964 and *H_o_* = 0.957 in *A. arenosa*; *H_e_* = 0.933 and *H_o_* = 0.865 in *A. lyrata*) than in tetraploids (*H_e_* = 0.699 and *H_o_*= 0.681 in *A. arenosa*; *H_e_* = 0.772 and *H_o_* = 0.715 in *A. lyrata*). On average, the reduction in *H_e_* and *H_o_* is thus about 30%, which is in the upper bound of the differences observed in our finite population simulations of the SSI-DOM model (Table 1).

**Table 2.** Estimation of the number of alleles, allelic richness after rarefaction, Nei’s gene diversity (*H*_e_) and observed heterozygosity computed as the gametic heterozygosity of an individual (*H*_o_, Moody et al. 1993) at the S-locus in *A. arenosa* and *A. lyrata* in overall samples from the diploid (2x) and tetraploid (4x) cytotypes, as well as in a pooled sample per species.

| | Cytotype | Number of individuals | Number of missing gene copies (%) | Number of S-alleles | Allelic richness (among 20 gene copies) | Gene diversity ($H_e$) | Observed heterozygosity ($H_o$) |
| --- | --- | --- | --- | --- | --- | --- | --- |
| *A. arenosa* | Pooled | 572 | 9 (0.5%) | 83 | 10.92 | 0.829 | 0.781 |
|  | 2x | 207 | 0 (0%) | 81 | 15.96 | 0.964 | 0.957 |
|  | 4x | 365 | 9 (0.6%) | 80 | 9.42 | 0.699 | 0.681 |
| *A. lyrata* | Pooled | 555 | 6 (1.1%) | 66 | 12.66 | 0.905 | 0.828 |
|  | 2x | 417 | 1 (0.2%) | 62 | 13.88 | 0.933 | 0.865 |
|  | 4x | 138 | 5 (3.6%) | 61 | 10.36 | 0.772 | 0.715 |

These results were qualitatively unchanged when estimates of S-locus diversity were computed on regional samples, corresponding with previously identified intraspecific genetic lineages within each cytotype gene pool (supplemental data Tables S5, S6). With the exception of the number of alleles (which is highly dependent on the sample size), all other statistics (allelic richness, gene diversity and observed heterozygosity) were very similar in regional samples as compared to overall samples within cytotype gene pools, and clearly indicate higher S-locus diversity in diploids than in tetraploids. Hence, the differences observed between cytotypes are not due to differences in population genetic structure.

Examination of S-allele frequency distributions revealed a striking difference between cytotypes. The most recessive S-allele (allele H1001, belonging to class I) was found at moderate frequencies of 0.162 and 0.227 in diploid *A. arenosa* and *A. lyrata*, but at the much higher frequencies of 0.546 and 0.471 in tetraploid *A. arenosa* and *A. lyrata*, respectively (Fig.2). Hence, in tetraploids, about 50% of the gene copies belong to a single S-allele, despite the overall high allelic diversity at the S-locus within the entire lineage. Indeed, in *A. arenosa*, of the 365 tetraploid individuals analyzed, 36 carried four copies of the H1001 allele, 107 individuals carried three copies, 124 individuals carried two copies, 75 individuals carried a single copy, and only 23 individuals did not carry any copy of H1001. Among the 137 tetraploid individuals of *A. lyrata*, 13 carried four copies of H1001, 29 carried three copies, 38 carried two copies, 39 carried a single copy, and 18 individuals did not carry any copy of H1001. As a result, the average frequencies of the other alleles (those from classes II to IV) are lower in tetraploids than in diploids in both species, with the exception of class II alleles in *A. lyrata* which show slightly higher average frequency in tetraploids than in diploids (Fig. 2). These observations are largely in line with numerical simulations under the SSI-DOM model (Fig. 1).

**Fig. 2.**
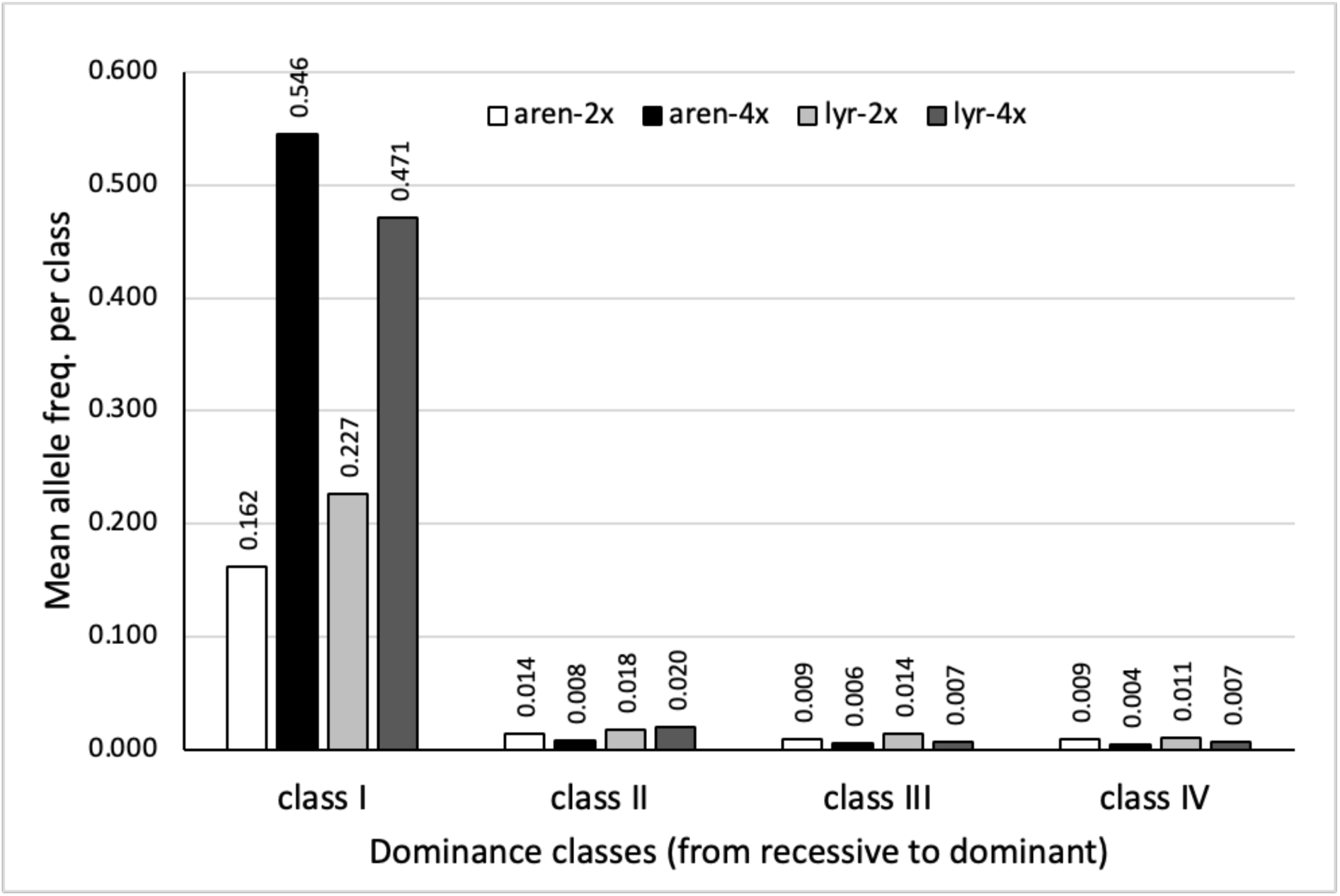
Average observed frequencies of S-alleles belonging to dominance classes I to IV in the diploid (white bars for *A. arenosa*, light grey bars for *A. lyrata*) and tetraploid (black bars for *A. arenosa*, dark grey bars for *A. lyrata*) gene pools.

### Nucleotide polymorphism among gene copies within S-alleles in diploid versus tetraploid populations

In order to test for founder effects associated with the origin of tetraploid populations, we looked for potential signatures on the within S-allele nucleotide diversity (**π**) at the *SRK* gene, and compared the average values of **π** between the two cytotype gene pools over alleles belonging to the same class of dominance (Table 3). In both *A. arenosa* and *A. lyrata*, average **π** values within S-alleles were not significantly different between diploids and tetraploids, and the net divergence between gene copies of a given S-allele from the two cytotypes was very low (Table 3). Hence, we found no reduction in polymorphism and genetic divergence within S-alleles in association with the origin and subsequent diversification of polyploids.

**Table 3.**
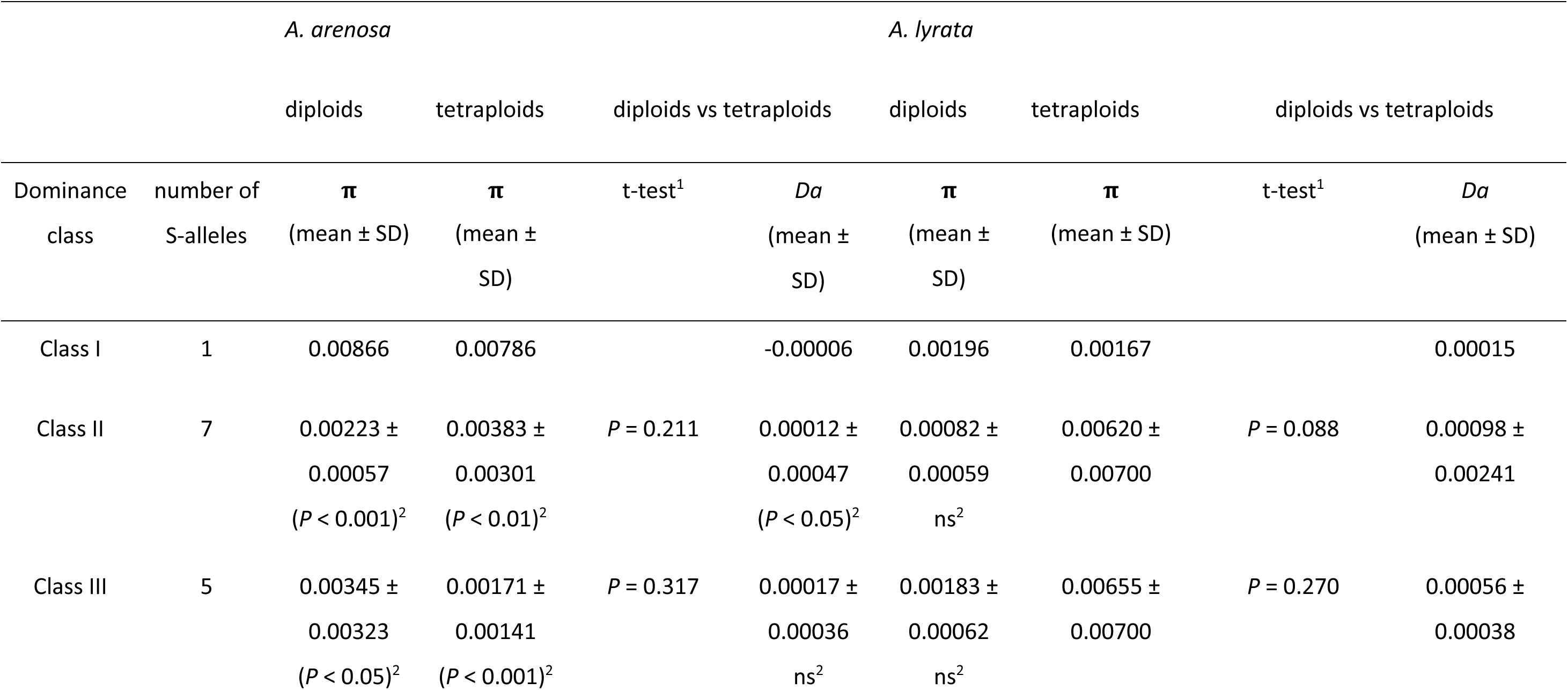
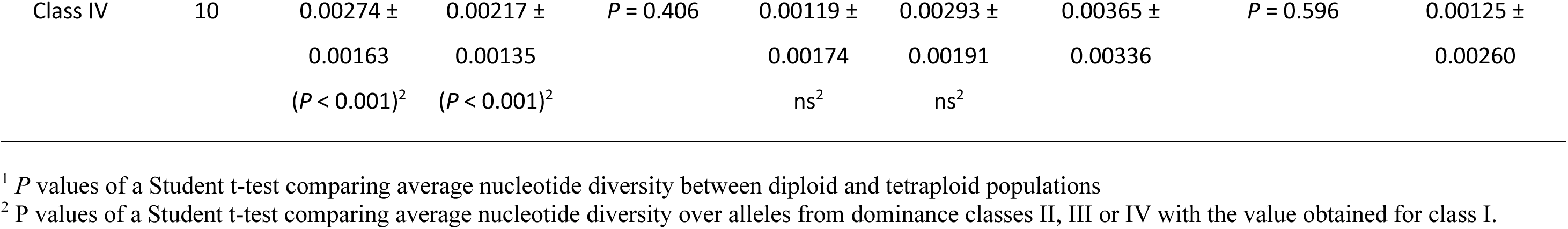
Comparison of within S-allele nucleotide diversity between diploid and tetraploid populations of *A. arenosa* and *A. lyrata*. Values of nucleotide diversity (π) and net nucleotide divergence (*Da*) between cytotypes are reported as averages over different alleles for each dominance class (except the most recessive class I, which has only one S-allele).

A striking effect of dominance was observed on levels of polymorphism within S-alleles in *A. arenosa*, where nucleotide diversity of the most recessive allele (class I) was significantly higher (by a factor varying between 2.5 to 3.9) than the average values for the other (more dominant) classes (Table 3). This is expected for SSI because of the higher effective population size of the most recessive S-allele (Castric et al. 2010). Despite the much higher frequency of the class I allele in tetraploids as compared to diploids (Fig. 2), we do not see a corresponding increase in within S-allele nucleotide diversity, probably indicating that mutation-drift equilibrium has not yet been reached due to the recent origin of polyploids (Monnahan et al. 2019). This higher nucleotide diversity for the most recessive S-allele was not apparent in *A. lyrata,* despite its higher frequency as compared to other dominance classes in diploid and especially so in tetraploid populations (Fig. 2).

## Discussion

### Polyploidy strongly modulates allele frequencies at the locus controlling sporophytic self-incompatibility

In the absence of selection, autopolyploidy is expected to result in higher levels of genetic diversity within populations at mutation-drift equilibrium, because with similar numbers of individuals, the total number of gene copies immediately doubles in polyploid as compared to diploid populations (Meirmans et al. 2018; Vlcek et al. 2025). This basic prediction should also apply to multiallelic SI systems in plants subject to negative frequency-dependent selection. Indeed, for gametophytic SI, the only SI system for which predictions in terms of allelic diversity in finite population have been derived analytically (Wright 1966; Czuppon & Billiard 2022), the main driver predicting allelic richness is the number of gene copies in the population. Under sporophytic SI, our simulation results show that this prediction holds in a model with strict co-dominance among S-alleles in both pollen and pistil (SSI-COD model in Schierup et al. 1997), with a two-fold increase in the number of S-alleles predicted to be maintained at equilibrium in tetraploid as compared to diploid populations. We note that the magnitude of the increase is higher than expected based on the doubling of the effective population size alone, as for instance the increase in number of alleles expected for a GSI model when doubling the population size is only about 1.5-fold instead of two-fold (e.g. from 15.7 to 23.4 alleles with *N* increasing from 200 to 400 with a mutation rate of 0.0005, according to the derivation of Yokoyama & Nei 1979). This is the consequence of a strong increase in the intensity of negative frequency-dependent selection under the SSI-COD model in tetraploids, because of its specific constraints in terms of mate compatibility imposing that all four alleles of the pollen parent should be different from each of the four alleles of the maternal parent. This model also imposes that the minimum number of S-alleles for a functionally self-incompatible tetraploid population to be viable is *n*=8 (versus *n*=4 in diploid populations; Schierup et al. 1997), therefore putting even stronger constraints on minimum viable population sizes.

In contrast to the SSI-COD model, the model of SSI with dominance in both pollen and pistil (SSI-DOM model in Schierup et al. 1997) predicted no major impact of polyploidy on the number of alleles maintained in populations, but a strong reduction in gene diversity and in observed heterozygosity at the S-locus. Inspection of S-allele frequencies indicated that the key to this unexpected result resides in the masking effect of dominance, which is more effective in polyploids (Otto 2007; Baduel et al. 2019). Indeed, dominance in sporophytic SI produces an asymmetry in allele frequencies, known as the "recessive effect" (Bateman 1952; Sampson 1974; Schierup et al. 1997; Billiard et al. 2007), whereby recessive S-alleles reach higher population frequencies than more dominant S-alleles as they are partially hidden from the action of negative frequency-dependent selection. Our simulations showed that this “recessive effect” is magnified in polyploids, as the ratio between the frequency of the most recessive and that of the most dominant S-allele was almost three times higher in polyploids than in diploids in finite populations (Table 1), and the frequency of the most recessive allele rose above 0.5 in deterministic simulations (Fig. 1 C and D). This increased asymmetry causes in turn a reduction in heterozygosity at the S-locus, which outperforms the increase in gene diversity due to increased effective population size (as seen in the absence of dominance, i.e. under the SSI-COD model). This constitutes a remarkable new example of the population genetic consequences of an increased dominance effect in polyploids.

These theoretical predictions were largely confirmed by our empirical results on the S-locus of *A. arenosa* and *A. lyrata*, as we indeed observed a strong reduction in allelic richness, gene diversity and observed heterozygosity in autotetraploid as compared to diploid populations of both species (Table 2). This reduction in heterozygosity cannot be explained by a reduction of allelic diversity, as the total number of S-alleles was similar in the tetraploid and the diploid gene pools. Also, it cannot be explained by double reduction in autotetraploids, the phenomenon by which two sister chromatids migrate to the same gamete during meiosis (Bever & Felber 1992), because although it would also cause a reduction in heterozygosity, it would not explain the observed increase in frequency of the most recessive allele in tetraploid as compared to diploid populations (Fig. 2). High proportions of tetraploid individuals carrying copies of the most recessive S-allele (here, 94% of *A. arenosa* tetraploids) were already reported by Mable et al. (2018) in *A. lyrata* and *A. arenosa*, although their method could not determine the exact number of gene copies for each S-allele within individuals. An important assumption of our implementation of the SSI-DOM model in tetraploids is that a single copy of the most dominant allele can impede expression of all three other gene copies in a tetraploid individual, *i.e.* that the pollen S-locus phenotype is determined by the allele with the highest dominance rank in a genotype, such that even a single dominant S-allele has the capacity to silence phenotypic expression of up to three more recessive S-alleles in a tetraploid genotype. This possibility was first presented by Mable et al. (2004), who examined segregation of S-locus phenotypes in crosses of tetraploid *A. lyrata*, and concluded that dosage effects did not alter the dominance relationships. Dominance between S-alleles is now known in Brassicaceae to be determined by S-locus-linked small RNAs produced by dominant S-alleles that specifically target and suppress expression of the *SCR* gene (controlling the pollen incompatibility phenotype) of recessive S-alleles (Durand et al. 2014; Tarutani et al. 2010). In a previous work, we showed that the transcriptional silencing mechanism is functional in synthetic autopolyploid *Capsella grandiflora* individuals, although we were unable to determine precisely the exact number of copies of the dominant and recessive S-alleles within tetraploid genotypes (Duan et al. 2024).

### No evidence for a major founder effect at the S-locus during evolution of tetraploid populations in *A. arenosa and A. lyrata*

Inferring the mode of origin of novel polyploid lineages is essential to determine the sources and fate of genetic variation (Kolar et al. 2017). The initial formation of polyploids via either somatic doubling or via crosses involving unreduced gametes (Ramsey & Schemske 1998, 2002) are believed to be rare events, such that drastic founder effects are expected to accompany the origin of polyploid lineages (Layman & Busch 2018). However, additional processes such as homoploid hybridisation among previously established tetraploid lineages and/or interploidy gene flow from diverse diploid ancestors may contribute to replenish the initially depauperated gene pools of the nascent polyploid (Soltis & Soltis 1999, Baduel et al. 2018, Schmickl and Yant 2021, Bartolic et al. 2024). The existence of founder effects is particularly important for multiallelic SI systems, as a decrease in S-allele diversity can reduce their resilience to the spread of self-compatibility mutations (Charlesworth & Charlesworth 1979; Brom et al. 2019). However, if multiple origins or interploidy gene flow quickly restore S-allele diversity in novel polyploids, such a shift towards selfing might not be favored.

In *A. arenosa* and *A. lyrata*, tetraploid populations are characterized by a functional SI system (Mable et al. 2004; Mable et al. 2018; Scott et al. 2024), demonstrating that a scenario of loss of SI (e.g. due an immediate functional effect or to a reduction in S-allele diversity) did not occur. Here, we showed that tetraploid populations host similar numbers of S-alleles as diploid populations, and that most S-alleles are shared between gene pools of the two cytotypes. This means that about 80 and 60 S-alleles in *A. arenosa* and *A. lyrata*, respectively, were transferred from the parental diploid gene pool to the tetraploid gene pool. This could have occurred through two different non-exclusive evolutionary mechanisms: either a high degree of repeated formation of autotetraploid individuals at the onset of the tetraploid gene pool, or a high degree of introgression of S-alleles from the diploid to the tetraploid gene pool, through inter-cytotype gene flow. In *A. arenosa*, no sign of genome-wide founder effect associated with polyploidization was observed, and a population of many individuals were inferred by coalescent simulations to have given rise to the autotetraploid lineage (Arnold et al. 2015). Furthermore, this variation was enriched by additional significant pulses of interploidy introgression from distinct diploid lineages of the same species in several secondary contact zones (Monnahan et al 2019). In *A. lyrata*, multiple populations gave rise to independent autotetraploid lineages experiencing gene flow, but the occurrence of founder effects has not been tested (Scott et al. 2024). In addition to comparisons of allelic diversity at the S-locus, we also compared levels of within S-allele nucleotide diversity between cytotypes. We hypothesized that independent founder effects within each S-allele transferred from the diploid parent to the tetraploid gene pool would cause a reduction in nucleotide diversity, and produce substantial nucleotide divergence between the cytotypes. Our results showed similar levels of diversity and low divergence among the two cytotypes for all alleles tested in both species (Table 3), confirming the absence of founder effects and/or an important role of genetic introgression between the two cytotypes (Monnahan et al. 2019).

### Different evolutionary impact of polyploidy on mating system evolution under autopolyploidy and allopolyploidy events in Brassicaceae

Associations between polyploidy events and mating system shifts from outcrossing to selfing have been largely debated in the plant biology literature, but a general consensus is still lacking (Barringer, 2007; Husband et al. 2008; Mable 2004; Robertson et al. 2011). Duan et al. (2024) argued that this confusion arises because of functional differences between the genetic systems enforcing outcrossing in parental diploid lineages (e.g. the type of SI systems) and their interaction with the mode of polyploidy (autopolyploidy vs allopolyploidy). In particular, evidence from studies of recent polyploids in Brassicaceae points to a general pattern of maintenance of a strictly outcrossing mating system with functional SI in autopolyploids derived from diploid self-incompatible parents, while allopolyploids show systematic loss of the SI system, often followed by evolution of high selfing (reviewed in Novikova et al. 2023). This pattern could be caused by several factors differentiating auto- and allopolyploids, such as differential reduction in S-allele diversity and/or inbreeding depression during founding events, factors which are known to facilitate the spread of self-compatible mutations within populations (Charlesworth & Charlesworth 1979; Porcher & Lande 2005; Gervais et al. 2014). Our results clearly indicate noreduction of S-allele diversity in autotetraploid as compared to diploid populations of *A. arenosa* and *A. lyrata*. In contrast, evolution of allopolyploids seems to be associated with founder effects, causing strong reduction in S-allele diversity (Novikova et al. 2007; Shimizu-Inatsugi et al. 2009; Duan et al. 2024). Regarding levels of inbreeding depression, theory (Lande & Schemske 1985; Ronfort 1999; Layman & Busch 2018) and some empirical studies (Husband & Schemske 1997; Husband et al. 2008; Siopa et al. 2020) seem to indicate that it would be reduced in autopolyploids as compared to the diploid parents, especially in nascent polyploids (Otto & Whitton 2000; Clo & Kolar 2022). This would cause a substantial increase in the probability of loss of SI, but this does not seem to fit with empirical evidence, as shown here. For allopolyploids, although we are not aware of results from dedicated population genetic models, one could argue that the expected strong founding effects associated with their formation should also cause a substantial initial reduction in inbreeding depression (Layman & Busch 2018), as predicted by theory for diploids (Kirkpatrick & Jarne 2000). Hence, the combination of potential reductions in S-allele diversity and inbreeding depression in young allopolyploids could be responsible for their observed pattern of loss of SI. Moreover, allopolyploidy in Brassicaceae is often found to evolve from hybridization between a selfing and a self-incompatible parental species or population (e.g. *Arabidopsis suecica*, *Arabidopsis kamchatika*, *Capsella bursa-pastoris*, all reviewed in Novikova et al. 2023; *Cardamine flexuosa*, Mandakova et al. 2014). In this case, non-functional S-alleles from the selfing parent will be introduced in the allopolyploid and could generate instant self-compatibility (Novikova et al. 2023; Duan et al. 2024). New empirical studies of the SI system in recent polyploid species with a well characterized mechanism of polyploidy (i.e. auto-vs allo-polyploidy) are greatly needed, especially in other plant families with sporophytic SI (e.g. Asteraceae, Convolvulaceae), to evaluate the generality of the association of SI with autopolyploidy on the one hand, and SC with allopolyploidy on the other hand.

## Materials & Methods

### Numerical simulations

We used the simulation program of SSI from Schierup et al. (1997) to compare the equilibrium allelic diversity and distribution of allele frequencies between diploid and autotetraploid populations. We investigated two of the models proposed by Schierup et al. (1997), representing contrasted situations regarding patterns of dominance among S-alleles: 1) a model with strict codominance in the determination of the pollen and pistil SI phenotypes (SSI-COD) and 2) a model with strictly hierarchical dominance relationships among alleles in both pollen and pistil (SSI-DOM). Deterministic equilibrium frequencies (Schierup et al. 1997) of the SSI-COD model showed identical expected values for all S-alleles (symmetrical model), whereas for the SSI-DOM model the expected frequencies were inversely related to dominance (asymmetrical model). According to Prigoda et al. (2005) and Llaurens et al. (2008) the latter is closer to the patterns determined empirically in outcrossing *Arabidopsis* species. We modified the simulation program, and in particular the algorithm to accept or reject a randomly chosen pollen, to implement autotetraploid individuals. We assumed polysomic inheritance and absence of double reduction in the tetraploids. For the SSI-DOM model, we assumed that dominance is qualitative rather than quantitative, meaning that a single most dominant allele copy can impede expression of all three other gene copies in a tetraploid individual (Duan et al. 2024), i.e. the S-locus phenotype of a given individual is determined by the allele with the highest dominance rank in its genotype. Each simulation was started with 2*N* (4*N* in tetraploids, with *N* the number of individuals) functionally different alleles in the population and allowed to evolve for 30,000 generations, a time chosen to ensure that mutation–selection–drift equilibrium was attained. We then computed the following statistics (averaged over 1,000 simulation replicates): number of S-alleles present; gene diversity (*H_e_*, probability that a randomly chosen pair of gene copies sampled from the overall cytotype gene pool corresponds to different S-alleles); observed heterozygosity (*H_o_*, computed in tetraploids as the gametic heterozygosity of an individual, i.e. the frequency of heterozygotes among randomly sampled diploid gametes formed from its four gene copies, according to Moody et al. 1993). For the SSI-DOM model, we also compared the frequency of the most recessive allele with that of the most dominant allele present in the population. For “finite population” simulations, we chose two parameter sets used by Schierup et al. (1997): *N* = 100 with a mutation rate *μ* = 0.0001; and *N* = 200 with a mutation rate *μ* = 0.0005. We used an infinite mutation model, and for the SSI-DOM model each new S-allele was placed randomly within the hierarchical dominance ladder. For comparisons with Schierup et al. (1997), we simulated negative frequency-dependent selection acting only on the male fitness, i.e. we did not implement fecundity selection under pollen limitation (Vekemans et al. 1998). For so-called “deterministic” simulations, we used a large population size (N = 7,000) and introduced a fixed number of S-alleles (5 or 12) in the population, in the absence of mutation.

### S-locus genotyping in diploid and tetraploid populations of A. arenosa and A. lyrata and population genetic analyses

For *A. arenosa*, we gathered a dataset of short-read whole-genome Illumina sequences for 207 diploid and 365 tetraploid individuals from Sequence Read Archive (SRA, listed in Supplementary Table S1). For *A. lyrata*, we gathered a similar dataset for 417 diploid and 138 tetraploid individuals (Table S2). For each individual, we analyzed the short-read data with the NGSgenotyp pipeline (Genete et al., 2020) to genotype the S-locus using a reference database of *SRK* sequences available from the related species *A. lyrata* and *A. halleri*, and from *Capsella grandiflora* (see Duan et al. 2024; available at https://www.doi.org/10.6084/m9.figshare.22567558.v2), to which we added a limited dataset of partial *SRK* sequences of *A. arenosa* available from Ruiz-Darte et al. (2012) and Mable et al. (2018). Briefly, this pipeline filters short-reads with a library of k-mers extracted from the reference database, then uses Bowtie2 to align filtered reads sequentially against each reference sequence and produces summary statistics with Samtools (v1.4; Danecek et al., 2021), allowing it to identify S-alleles present in each individual and their copy number. An important challenge in autopolyploids is to be able to discriminate among alternative partial heterozygote genotypes (Dufresne et al. 2014), and this was achieved using NGSgenotyp thanks to the deeply sequenced individual genomes. The NGSgenotyp pipeline also contains a *de novo* assembly module, which produces full sequences of the S-domain of *SRK* for alleles present as partial sequences in the database as well as for newly identified S-alleles. The population genetic statistics described in the previous section, as well as allelic richness computed with rarefaction methods, were analyzed based on the S-locus genotypes of diploid and tetraploid genotypes with the software SPAGEDI that allows datasets with mixed ploidy and incomplete genotypes (Hardy & Vekemans 2002; available at https://github.com/reedacartwright/spagedi). These analyses were performed at different levels: (1) at the species-level; (2) within each cytotype gene pool; (3) within regional samples for each cytotype gene pool (see Tables S1 and S2 for individual assignments to regional samples in *A. arenosa* and *A. lyrata*, respectively). In order to compare observed S-allele frequencies between the two cytotype gene pools and between classes of dominance, we grouped alleles based on phylogenetic relationships (Prigoda et al. 2005; Durand et al. 2014) according to the classification of Goubet et al. (2012), from class I (the most recessive) to class IV (the most dominant alleles), and computed average frequencies per class. In this paper, we used the S-allele identification scheme introduced by Duan et al. (2024), in the form HXYYY, with X corresponding to the dominance class of the allele (ranging from 1 to 4), and YYY corresponding to its functional specificity (ranging from 001 up to a limit of 999) determined based on phylogenetic relationships among *SRK* sequences from different species.

### Analyses of nucleotide diversity at the SRK gene within S-alleles

In order to test for founder effects associated with the origin of tetraploid populations on haplotype diversity at the S-locus, we gathered *SRK* sequence datasets of multiple gene copies from diploid and tetraploid individuals, for a subsample of S-alleles found in each species. Sequences from different *SRK* gene copies of the same S-allele were obtained with the *de novo* assembly module of NGSgenotyp based on individual short read sequencing data. After alignment of gene copy sequences for each S-allele, we computed nucleotide diversity, separately for diploids and tetraploids, and average net nucleotide divergence between the two cytotypes, using the maximum composite likelihood method implemented in Mega V.12 (Kumar et al. 2024). We then computed average nucleotide diversities over alleles within the same dominance class and tested for differences between the two cytotypes using a Student’t-test with the t.test function of the R Statistical Software (v4.5.0; R Core Team 2021).

## Data accessibility

Supplementary material is available online. Sequence data for *SRK* alleles in *A. arenosa* and *A. lyrata* are available here: https://www.doi.org/10.6084/m9.figshare.29294138.

