## Supplemental data for "Autopolyploidy exacerbates dominance masking under negative frequency-dependent selection : evidence from sporophytic self-incompatibility in *Arabidopsis arenosa* and *A. lyrata*"

### Supplemental information

**Table S1.** S-locus genotypes of 207 diploid and 365 tetraploid individuals from *Arabidopsis arenosa* , as obtained by the NGSgenotyp pipeline on whole-genome resequencing data retrieved from Genbank (SRA accession ID indicated). Allele sequences are identified according to a Brassicaceae functional sequence group nomenclature, as determined based on sequence similarity with alleles from related Arabidopsis species (see Table S3 for correspondence with *A. arenosa* -specific S-allele IDs).

| Sample | ploidy | Population | Regional sample | allele #1 | allele #2 | allele #3 | allele #4 | SRA_accession* | Depth of coverage |
|---|---|---|---|---|---|---|---|---|---|
| BAB_01da | 2 | BAB | WCarpathian | H1001 | H3022 | | | SRR10549990 | 48.9 |
| BAB_03da | 2 | BAB | WCarpathian | H4006 | H4036 | | | SRR23469399 | 40.7 |
| BAB_04da | 2 | BAB | WCarpathian | H2001 | H2007 | | | SRR10549988 | 43.3 |
| BAB_05da | 2 | BAB | WCarpathian | H2008 | H4029 | | | SRR10549987 | 35.6 |
| BAB_06da | 2 | BAB | WCarpathian | H1001 | H3028 | | | SRR10549985 | 30.0 |
| BAB_08da | 2 | BAB | WCarpathian | H4013 | H4018 | | | SRR10549984 | 19.8 |
| BAB_09da | 2 | BAB | WCarpathian | H2001 | H4035 | | | SRR10549983 | 45.5 |
| BAB_10da | 2 | BAB | WCarpathian | H3003 | H3009 | | | SRR23469397 | 44.8 |
| BAB_11da | 2 | BAB | WCarpathian | H1001 | H2024 | | | SRR23469375 | 34.6 |
| BAB_12da | 2 | BAB | WCarpathian | H1001 | H4031 | | | SRR10549982 | 42.3 |
| BAB_13da | 2 | BAB | WCarpathian | H1001 | H3022 | | | SRR23469353 | 47.6 |
| BAL_01ta | 4 | BAL | SCarpathian | H1001 | H2028 | H4010 | H4027 | SRR10549963 | 43.6 |
| BAL_02ta | 4 | BAL | SCarpathian | H1001 | H1001 | H1001 | H1001 | SRR10549962 | 36.8 |
| BAL_03ta | 4 | BAL | SCarpathian | H1001 | H1001 | H2004 | H4034 | SRR10549961 | 33.7 |
| BAL_04ta | 4 | BAL | SCarpathian | H1001 | H1001 | H1001 | H4027 | SRR10549960 | 60.6 |
| BAL_05ta | 4 | BAL | SCarpathian | H1001 | H1001 | H1001 | H4003 | SRR10549959 | 49.0 |
| BAL_06ta | 4 | BAL | SCarpathian | H1001 | H1001 | H3004 | H3015 | SRR10549958 | 36.4 |
| BAL_07ta | 4 | BAL | SCarpathian | H1001 | H1001 | H3025 | H4034 | SRR10549957 | 38.4 |
| BAL_08ta | 4 | BAL | SCarpathian | H1001 | H1001 | H2004 | H3004 | SRR10549956 | 43.1 |
| BDO_01da | 2 | BDO | Pannonian | H2010 | H4005 | | | SRR23282202 | 25.4 |
| BDO_02da | 2 | BDO | Pannonian | H4019 | H4043 | | | SRR23282201 | 21.7 |
| BDO_03da | 2 | BDO | Pannonian | H2001 | H4024 | | | SRR23282145 | 27.9 |
| BDO_04da | 2 | BDO | Pannonian | H4040 | H4042 | | | SRR23282208 | 23.7 |
| BDO_05da | 2 | BDO | Pannonian | H1001 | H4020 | | | SRR23282109 | 34.4 |
| BDO_06da | 2 | BDO | Pannonian | H2004 | H2004 | | | SRR23282098 | 25.4 |
| BDO_07da | 2 | BDO | Pannonian | H1001 | H3006 | | | SRR23282055 | 22.8 |
| BDO_08da | 2 | BDO | Pannonian | H3001 | H4022 | | | SRR23282044 | 26.8 |
| BEL_01da | 2 | BEL | Dinaric | H2010 | H4048 | | | SRR7637493 | 10.9 |
| BEL_02da | 2 | BEL | Dinaric | H3025 | H4026 | | | SRR7637492 | 11.3 |
| BEL_03da | 2 | BEL | Dinaric | H2001 | H4048 | | | SRR7637491 | 11.9 |
| BEL_04da | 2 | BEL | Dinaric | H4001 | H4034 | | | SRR7637317 | 12.9 |
| BEL_05da | 2 | BEL | Dinaric | H3024 | H4020 | | | SRR7637489 | 13.1 |
| BEL_06da | 2 | BEL | Dinaric | H3024 | H4009 | | | SRR7637320 | 15.0 |
| BEL_07da | 2 | BEL | Dinaric | H4014 | H4017 | | | SRR7637487 | 15.4 |
| BEL_08da | 2 | BEL | Dinaric | H4009 | H4013 | | | SRR7637486 | 16.0 |
| BGS_01ta | 4 | BGS | Ruderal-Alpine admix | H1001 | H2024 | H3025 | H4011 | SRR7637496 | 12.9 |
| BGS_02ta | 4 | BGS | Ruderal-Alpine admix | H1001 | H2026 | H2006 | H4006 | SRR7637495 | 14.4 |
| BGS_03ta | 4 | BGS | Ruderal-Alpine admix | H1001 | H1001 | H1001 | H0000 | SRR7637465 | 12.5 |
| BGS_04ta | 4 | BGS | Ruderal-Alpine admix | H1001 | H1001 | H1001 | H3014 | SRR7637468 | 14.0 |
| BGS_05ta | 4 | BGS | Ruderal-Alpine admix | H1001 | H4010 | H4015 | H4021 | SRR7637467 | 16.5 |
| BGS_06ta_fi | 4 | NA | Baltic | H1001 | H1001 | H1001 | H2006 | SRR7637466 | NA |
| BGS_07ta_fi | 4 | NA | Baltic | H1001 | H1001 | H2026 | H3025 | SRR7637462 | NA |
| BGS_08ta_fi | 4 | NA | Baltic | H1001 | H1001 | H1001 | H2007 | SRR7637461 | NA |
| BIH_01da | 2 | BIH | Dinaric | H3002 | H3008 | | | SRR7637464 | 11.0 |
| BIH_02da | 2 | BIH | Dinaric | H2009 | H3014 | | | SRR7637463 | 10.2 |
| BIH_03da | 2 | BIH | Dinaric | H4009 | H2010 | | | SRR7637460 | 12.6 |
| BIH_04da | 2 | BIH | Dinaric | H4023 | H2021 | | | SRR7637459 | 11.9 |
| BIH_05da | 2 | BIH | Dinaric | H4008 | H4035 | | | SRR7637354 | 12.5 |
| BIH_06da | 2 | BIH | Dinaric | H3002 | H4012 | | | SRR7637355 | 13.6 |
| BIH_07da | 2 | BIH | Dinaric | H2008 | H4009 | | | SRR7637356 | 9.9 |
| BIH_08da | 2 | BIH | Dinaric | H3022 | H4002 | | | SRR7637357 | 11.9 |
| BOR_01ta | 4 | BOR | Hercynian | H1001 | H2026 | H2008 | H4017 | SRR12778344 | 18.4 |
| BOR_02ta | 4 | BOR | Hercynian | H1001 | H3004 | H4015 | H4023 | SRR12778343 | 16.3 |
| BOR_03ta | 4 | BOR | Hercynian | H1001 | H1001 | H4008 | H2024 | SRR12778332 | 14.9 |
| BOR_04ta | 4 | BOR | Hercynian | H1001 | H4001 | H4019 | H4020 | SRR12778321 | 15.3 |
| BOR_05ta | 4 | BOR | Hercynian | H1001 | H1001 | H1001 | H4023 | SRR12778310 | 15.9 |
| BOR_06ta | 4 | BOR | Hercynian | H1001 | H1001 | H1001 | H2027 | SRR12778299 | 16.6 |
| BOR_07ta | 4 | BOR | Hercynian | H1001 | H1001 | H2008 | H3010 | SRR12778288 | 15.9 |
| BOR_08ta | 4 | BOR | Hercynian | H1001 | H1001 | H1001 | H3010 | SRR12778277 | 17.3 |
| BRD_01ta | 4 | BRD | Hercynian | H1001 | H2004 | H2005 | H4034 | SRR7637358 | NA |
| BRD_02ta | 4 | BRD | Hercynian | H1001 | H1001 | H1001 | H2006 | SRR7637359 | NA |
| BRD_03ta | 4 | BRD | Hercynian | H1001 | H1001 | H3005 | H4013 | SRR7637360 | NA |
| BRD_04ta | 4 | BRD | Hercynian | H1001 | H1001 | H1001 | H1001 | SRR7637361 | NA |
| BRD_05ta | 4 | BRD | Hercynian | H1001 | H2019 | H3015 | H4034 | SRR7637421 | NA |
| BUD_01da | 2 | BUD | SCarpathian_2x | H2007 | H4007 | | | SRR23281946 | 46.0 |

| | | | | | | | | | |
|---|---|---|---|---|---|---|---|---|---|
| BUD_02da | 2 | BUD | SCarpathian_2x | H4041 | H2026 | | | SRR23281935 | 29.6 |
| BUD_03da | 2 | BUD | SCarpathian_2x | H2004 | H4019 | | | SRR23282200 | 37.8 |
| BUD_04da | 2 | BUD | SCarpathian_2x | H2004 | H4004 | | | SRR23282189 | 24.6 |
| BUD_05da | 2 | BUD | SCarpathian_2x | H4012 | H2018 | | | SRR23282178 | 51.1 |
| BUD_06da | 2 | BUD | SCarpathian_2x | H4026 | H2026 | | | SRR23282028 | 25.3 |
| BUD_07da | 2 | BUD | SCarpathian_2x | H2004 | H3004 | | | SRR23282017 | 28.3 |
| BUD_08da | 2 | BUD | SCarpathian_2x | H1001 | H4017 | | | SRR23282006 | 78.0 |
| BUT_01ta | 4 | BUT | SCarpathian | H1001 | H3011 | H3028 | H4016 | SRR23281995 | 37.3 |
| BUT_02ta | 4 | BUT | SCarpathian | H1001 | H1001 | H3003 | H4010 | SRR23281984 | 24.7 |
| BUT_03ta | 4 | BUT | SCarpathian | H1001 | H1001 | H3012 | H3025 | SRR23282167 | 26.1 |
| BUT_04ta | 4 | BUT | SCarpathian | H1001 | H2003 | H2009 | H4010 | SRR23282156 | 25.0 |
| BUT_05ta | 4 | BUT | SCarpathian | H1001 | H1001 | H2004 | H2009 | SRR23282144 | 43.9 |
| BUT_06ta | 4 | BUT | SCarpathian | H1001 | H2009 | H3025 | H4003 | SRR23282133 | 29.9 |
| BUT_07ta | 4 | BUT | SCarpathian | H1001 | H1001 | H2008 | H3025 | SRR23282122 | 26.0 |
| BUT_08ta | 4 | BUT | SCarpathian | H4023 | H4023 | H2021 | H2021 | SRR23282085 | 23.1 |
| CAR_01ta | 4 | CAR | ECarpathian | H2005 | H3022 | H2022 | H2022 | SRR10549938 | 49.5 |
| CAR_02ta | 4 | CAR | ECarpathian | H2005 | H3022 | H4038 | H4041 | SRR10549937 | 58.2 |
| CAR_03ta | 4 | CAR | ECarpathian | H1001 | H1001 | H4011 | H4036 | SRR10549936 | 24.6 |
| CAR_04ta | 4 | CAR | ECarpathian | H1001 | H1001 | H3011 | H4020 | SRR10549935 | 35.2 |
| CAR_05ta | 4 | CAR | ECarpathian | H1001 | H1001 | H2006 | H4035 | SRR10549934 | 36.9 |
| CAR_06ta | 4 | CAR | ECarpathian | H1001 | H4001 | H4004 | H4036 | SRR10549933 | 27.9 |
| CAR_07ta | 4 | CAR | ECarpathian | H1001 | H2001 | H3004 | H4011 | SRR10549932 | 30.7 |
| CAR_08ta | 4 | CAR | ECarpathian | H2001 | H2025 | H3006 | H4018 | SRR10549930 | 44.9 |
| CHO_01ta | 4 | CHO | Swabian | H1001 | H1001 | H1001 | H4016 | SRR7637363 | NA |
| CHO_02ta | 4 | CHO | Swabian | H1001 | H1001 | H2003 | H4034 | SRR7637316 | NA |
| CHO_04ta | 4 | CHO | Swabian | H1001 | H1001 | H4014 | H2022 | SRR7637315 | NA |
| CHO_05ta | 4 | CHO | Swabian | H1001 | H1001 | H1001 | H4015 | SRR7637314 | NA |
| CHO_06ta | 4 | CHO | Swabian | H1001 | H1001 | H1001 | H2022 | SRR7637313 | NA |
| CHO_07ta | 4 | CHO | Swabian | H1001 | H1001 | H1001 | H0000 | SRR7637312 | NA |
| CHO_08ta | 4 | CHO | Swabian | H1001 | H1001 | H3024 | H2022 | SRR7637311 | NA |
| CVH_01da | 2 | CVH | WCarpathian | H1001 | H4026 | | | SRR3111435 | NA |
| DON_01da | 2 | DON | WCarpathian | H1001 | H4035 | | | SRR3111437 | NA |
| DRA_01ta | 4 | DRA | SCarpathian | H1001 | H1001 | H2007 | H2007 | SRR7637350 | 12.3 |
| DRA_02ta | 4 | DRA | SCarpathian | H1001 | H1001 | H1001 | H2009 | SRR7637347 | 11.0 |
| DRA_03ta | 4 | DRA | SCarpathian | H1001 | H1001 | H2007 | H4007 | SRR7637343 | 9.7 |
| DRA_04ta | 4 | DRA | SCarpathian | H1001 | H1001 | H1001 | H4010 | SRR7637344 | 10.0 |
| DRA_05ta | 4 | DRA | SCarpathian | H1001 | H1001 | H2026 | H3003 | SRR7637351 | 8.2 |
| DRA_06ta | 4 | DRA | SCarpathian | H2007 | H2007 | H2026 | H4035 | SRR7637352 | 13.8 |
| DRA_07ta | 4 | DRA | SCarpathian | H1001 | H2005 | H4005 | H4017 | SRR7637349 | 9.6 |
| DRA_08ta | 4 | DRA | SCarpathian | H2007 | H3005 | H3005 | H4028 | SRR7637391 | 12.4 |
| DUM_01ta | 4 | DUM | WCarpathian | H1001 | H1001 | H4026 | H4034 | SRR2040831 | NA |
| EIS_01ta | 4 | EIS | Alpine | H1001 | H3025 | H4004 | H4010 | SRR2040821 | NA |
| FOJ_01da | 2 | FOJ | Dinaric | H1001 | H4007 | | | SRR7637301 | 14.3 |
| FOJ_02da | 2 | FOJ | Dinaric | H1001 | H3025 | | | SRR7637299 | 13.1 |
| FOJ_03da | 2 | FOJ | Dinaric | H1001 | H2005 | | | SRR7637302 | 13.9 |
| FOJ_04da | 2 | FOJ | Dinaric | H1001 | H2018 | | | SRR7637303 | 15.8 |
| FOJ_06da | 2 | FOJ | Dinaric | H2004 | H4017 | | | SRR7637304 | 16.8 |
| FOJ_07da | 2 | FOJ | Dinaric | H1001 | H4014 | | | SRR7637306 | 17.5 |
| FOJ_08da | 2 | FOJ | Dinaric | H2007 | H3006 | | | SRR7637300 | 13.9 |
| FUG_01ta | 4 | FUG | Hercynian | H1001 | H1001 | H2026 | H4003 | SRR12778276 | 20.0 |
| FUG_02ta | 4 | FUG | Hercynian | H1001 | H1001 | H3011 | H4041 | SRR12778275 | 15.9 |
| FUG_03ta | 4 | FUG | Hercynian | H1001 | H1001 | H2006 | H4042 | SRR12778342 | 15.7 |
| FUG_04ta | 4 | FUG | Hercynian | H1001 | H1001 | H3001 | H2026 | SRR12778341 | 17.1 |
| FUG_05ta | 4 | FUG | Hercynian | H1001 | H1001 | H2006 | H2026 | SRR12778340 | 18.2 |
| FUG_06ta | 4 | FUG | Hercynian | H1001 | H1001 | H2006 | H3012 | SRR12778339 | 18.9 |
| FUG_07ta | 4 | FUG | Hercynian | H1001 | H2026 | H3006 | H4002 | SRR12778338 | 18.1 |
| FUG_08ta | 4 | FUG | Hercynian | H1001 | H1001 | H2026 | H4015 | SRR12778337 | 15.3 |
| GOR_01da | 2 | GOR | ECarpathian_2x | H1001 | H3015 | | | SRR7637559 | 13.1 |
| GOR_02da | 2 | GOR | ECarpathian_2x | H1001 | H3015 | | | SRR7637558 | 16.0 |
| GOR_03da | 2 | GOR | ECarpathian_2x | H1001 | H1001 | | | SRR7637525 | 18.1 |
| GOR_04da | 2 | GOR | ECarpathian_2x | H4019 | H4034 | | | SRR7637555 | 11.6 |
| GOR_05da | 2 | GOR | ECarpathian_2x | H1001 | H4035 | | | SRR7637526 | 14.0 |
| GOR_06da | 2 | GOR | ECarpathian_2x | H4007 | H4041 | | | SRR7637307 | 13.7 |
| GOR_07da | 2 | GOR | ECarpathian_2x | H1001 | H4028 | | | SRR7637308 | 15.8 |
| GOR_08da | 2 | GOR | ECarpathian_2x | H1001 | H1001 | | | SRR7637514 | 14.1 |
| GUL_01ta | 4 | GUL | Alpine | H1001 | H1001 | H2004 | H2006 | SRR7637556 | 19.2 |
| GUL_02ta | 4 | GUL | Alpine | H1001 | H1001 | H1001 | H2019 | SRR7637557 | 21.5 |
| GUL_04ta | 4 | GUL | Alpine | H1001 | H1001 | H3001 | H3003 | SRR7637563 | 10.9 |
| GUL_05ta | 4 | GUL | Alpine | H1001 | H1001 | H1001 | H3025 | SRR7637520 | 31.1 |
| GUL_06ta | 4 | GUL | Alpine | H1001 | H1001 | H1001 | H2003 | SRR7637519 | 19.3 |
| GUL_07ta | 4 | GUL | Alpine | H1001 | H1001 | H1001 | H1001 | SRR7637518 | 21.5 |
| GUL_08ta | 4 | GUL | Alpine | H1001 | H1001 | H1001 | H1001 | SRR7637517 | 17.7 |
| GUL_09ta | 4 | GUL | Alpine | H1001 | H1001 | H1001 | H2004 | SRR7637524 | 14.6 |

| | | | | | | | | | |
|---|---|---|---|---|---|---|---|---|---|
| GUL_10ta | 4 | GUL | Alpine | H1001 | H1001 | H2024 | H0000 | SRR7637523 | 21.2 |
| GUL_11ta | 4 | GUL | Alpine | H1001 | H1001 | H1001 | H1001 | SRR12778336 | 21.8 |
| GUL_12ta | 4 | GUL | Alpine | H1001 | H1001 | H1001 | H4003 | SRR12778335 | 19.3 |
| GUL_13ta | 4 | GUL | Alpine | H1001 | H2009 | H3008 | H4003 | SRR12778334 | 14.8 |
| GUL_14ta | 4 | GUL | Alpine | H1001 | H1001 | H2006 | H2003 | SRR12778333 | 19.4 |
| GUL_15ta | 4 | GUL | Alpine | H1001 | H1001 | H2001 | H2004 | SRR12778331 | 17.4 |
| GUL_16ta | 4 | GUL | Alpine | H1001 | H3007 | H3014 | H4037 | SRR12778330 | 19.1 |
| GUL_18ta | 4 | GUL | Alpine | H1001 | H1001 | H1001 | H3006 | SRR12778328 | 16.6 |
| GUL_19ta | 4 | GUL | Alpine | H1001 | H1001 | H1001 | H2022 | SRR12778327 | 30.2 |
| HAR_01ta | 4 | HAR | Hercynian | H1001 | H1001 | H1001 | H2028 | SRR7637522 | NA |
| HLI_01da | 2 | HLI | WCarpathian | H1001 | H4041 | | | SAMN03733467 | NA |
| HLI_02da | 2 | HLI | WCarpathian | H2005 | H4022 | | | SRR3111445 | NA |
| HMC_01da | 2 | HMC | WCarpathian | H2011 | H3006 | | | SRR2040771 | NA |
| HMC_02da | 2 | HMC | WCarpathian | H2006 | H2008 | | | SRR2040772 | NA |
| HMC_03da | 2 | HMC | WCarpathian | H2010 | H4003 | | | SRR2040773 | NA |
| HMC_04da | 2 | HMC | WCarpathian | H2026 | H3008 | | | SRR3111434 | NA |
| HNE_01da | 2 | HNE | Pannonian | H3004 | H4022 | | | SRR7637516 | 11.8 |
| HNE_02da | 2 | HNE | Pannonian | H2024 | H3015 | | | SRR7637515 | 9.4 |
| HNE_03da | 2 | HNE | Pannonian | H2004 | H4004 | | | SRR7637446 | 10.8 |
| HNE_04da | 2 | HNE | Pannonian | H4015 | H4015 | | | SRR7637469 | 9.7 |
| HNE_05da | 2 | HNE | Pannonian | H4010 | H4015 | | | SRR7637353 | 8.3 |
| HNE_06da | 2 | HNE | Pannonian | H2008 | H4027 | | | SRR7637447 | 9.1 |
| HNE_07da | 2 | HNE | Pannonian | H2021 | H4017 | | | SRR7637426 | 7.5 |
| HNI_02da_fi | 2 | HNI | arpathian-ECarpathian | H1001 | H3004 | | | SRR7637401 | NA |
| HOC_01ta | 4 | HOC | Alpine | H1001 | H1001 | H3012 | H4010 | SRR7637425 | 17.7 |
| HOC_03ta | 4 | HOC | Alpine | H1001 | H2007 | H4010 | H4018 | SRR7637398 | 10.9 |
| HOC_04ta | 4 | HOC | Alpine | H3006 | H3015 | H4029 | H4030 | SRR7637429 | 13.5 |
| HOC_05ta | 4 | HOC | Alpine | H1001 | H1001 | H2027 | H4042 | SRR7637428 | 12.7 |
| HOC_06ta | 4 | HOC | Alpine | H1001 | H1001 | H2008 | H2018 | SRR7637420 | 12.7 |
| HOC_07ta | 4 | HOC | Alpine | H1001 | H1001 | H2009 | H3006 | SRR7637364 | 14.2 |
| HOC_08ta | 4 | HOC | Alpine | H1001 | H1001 | H3011 | H4035 | SRR7637397 | 11.4 |
| HRA_01ta | 4 | HRA | WCarpathian | H1001 | H1001 | H1001 | H1001 | SRR10549929 | 47.2 |
| HRA_02ta | 4 | HRA | WCarpathian | H1001 | H3007 | H3021 | H4013 | SRR10549928 | 52.1 |
| HRA_04ta | 4 | HRA | WCarpathian | H1001 | H1001 | H2024 | H3007 | SRR10549927 | 58.5 |
| HRA_05ta | 4 | HRA | WCarpathian | H1001 | H1001 | H2010 | H3002 | SRR10549926 | 34.8 |
| HRA_06ta | 4 | HRA | WCarpathian | H1001 | H1001 | H3002 | H4034 | SRR10549925 | 44.9 |
| HRA_07ta | 4 | HRA | WCarpathian | H2001 | H2022 | H3003 | H4034 | SRR12778325 | 30.3 |
| HRA_08ta | 4 | HRA | WCarpathian | H1001 | H1001 | H1001 | H3028 | SRR10549924 | 26.0 |
| HRA_09ta | 4 | HRA | WCarpathian | H1001 | H1001 | H2003 | H3025 | SRR12778324 | 29.5 |
| HRA_10ta | 4 | HRA | WCarpathian | H1001 | H1001 | H2003 | H4010 | SRR12778323 | 36.7 |
| HRA_12ta | 4 | HRA | WCarpathian | H1001 | H1001 | H1001 | H2027 | SRR10549923 | 44.4 |
| HRA_13ta | 4 | HRA | WCarpathian | H1001 | H1001 | H2010 | H2019 | SRR10549922 | 44.4 |
| HRC_01ta | 4 | HRC | Alpine | H2022 | H2022 | H1001 | H4036 | SRR2040825 | NA |
| HRN_01ta | 4 | HRN | Ruderal | H1001 | H1001 | H1001 | H4002 | SRR2040813 | 36.2 |
| HRN_02ta | 4 | HRN | Ruderal | H1001 | H1001 | H1001 | H4010 | SRR2040814 | 31.3 |
| HRN_03ta | 4 | HRN | Ruderal | H1001 | H2006 | H4001 | H4037 | SRR2040815 | 38.3 |
| HRN_04ta | 4 | HRN | Ruderal | H1001 | H1001 | H1001 | H1001 | SRR2040816 | 31.7 |
| HRN_05ta | 4 | HRN | Ruderal | H1001 | H1001 | H1001 | H4037 | SRR2040817 | 28.5 |
| HRN_06ta | 4 | HRN | Ruderal | H1001 | H1001 | H1001 | H4030 | SRR2040818 | 32.0 |
| INE_01ta | 4 | INE | ECarpathian | H1001 | H1001 | H1001 | H2006 | SRR10549946 | 56.9 |
| INE_02ta | 4 | INE | ECarpathian | H1001 | H1001 | H1001 | H1001 | SRR10549945 | 51.1 |
| INE_03ta | 4 | INE | ECarpathian | H1001 | H1001 | H2007 | H4009 | SRR10549944 | 43.5 |
| INE_04ta | 4 | INE | ECarpathian | H1001 | H1001 | H1001 | H2006 | SRR10549943 | 52.1 |
| INE_05ta | 4 | INE | ECarpathian | H1001 | H1001 | H1001 | H4009 | SRR10549941 | 34.8 |
| INE_06ta | 4 | INE | ECarpathian | H1001 | H1001 | H1001 | H2018 | SRR10549940 | 63.8 |
| INE_08ta | 4 | INE | ECarpathian | H2006 | H2028 | H3009 | H4038 | SRR10549939 | 45.7 |
| ING_01ta | 4 | ING | Alpine | H1001 | H1001 | H1001 | H4020 | SRR10549972 | 23.2 |
| ING_02ta | 4 | ING | Alpine | H1001 | H1001 | H3022 | H4041 | SRR10549971 | 15.6 |
| ING_03ta | 4 | ING | Alpine | H1001 | H1001 | H1001 | H3022 | SRR10549970 | 16.3 |
| ING_04ta | 4 | ING | Alpine | H1001 | H1001 | H1001 | H1001 | SRR10549969 | 17.1 |
| ING_05ta | 4 | ING | Alpine | H1001 | H2001 | H2008 | H3022 | SRR10549968 | 18.7 |
| ING_06ta | 4 | ING | Alpine | H1001 | H2001 | H2027 | H3022 | SRR10549967 | 19.7 |
| ING_07ta | 4 | ING | Alpine | H1001 | H1001 | H1001 | H4003 | SRR10549966 | 18.6 |
| ING_08ta | 4 | ING | Alpine | H1001 | H1001 | H2007 | H4017 | SRR10549965 | 17.5 |
| JVR_01da | 2 | JVR | WCarpathian | H2026 | H4039 | | | SRR2040808 | NA |
| JZC_01da | 4 | JZC | WCarpathian | H4004 | H4041 | H2024 | H1001 | SRR2040774 | NA |
| KAM_01ta | 4 | KAM | WCarpathian | H1001 | H1001 | H2003 | H4031 | SRR23282074 | 31.0 |
| KAM_02ta | 4 | KAM | WCarpathian | H1001 | H1001 | H2006 | H2019 | SRR23282063 | 31.6 |
| KAM_03ta | 4 | KAM | WCarpathian | H1001 | H2006 | H2026 | H4009 | SRR23281971 | 36.1 |
| KAM_04ta | 4 | KAM | WCarpathian | H1001 | H1001 | H1001 | H1001 | SRR23281960 | 42.1 |
| KAM_05ta | 4 | KAM | WCarpathian | H1001 | H1001 | H4010 | H4025 | SRR23281949 | 42.3 |
| KAM_06ta | 4 | KAM | WCarpathian | H1001 | H1001 | H2001 | H2006 | SRR23282219 | 32.9 |
| KAM_07ta | 4 | KAM | WCarpathian | H1001 | H2026 | H4034 | H4043 | SRR23282207 | 38.0 |

| | | | | | | | | | |
|---|---|---|---|---|---|---|---|---|---|
| KAM_08ta | 4 | KAM | WCarpathian | H1001 | H3024 | H4037 | H2019 | SRR23282118 | 35.1 |
| KAS_01ta | 4 | KAS | Alpine | H1001 | H1001 | H1001 | H2024 | SRR7637366 | 13.2 |
| KAS_02ta | 4 | KAS | Alpine | H1001 | H1001 | H1001 | H3010 | SRR7637458 | 14.3 |
| KAS_03ta | 4 | KAS | Alpine | H1001 | H1001 | H1001 | H2007 | SRR7637537 | 16.5 |
| KAS_04ta | 4 | KAS | Alpine | H1001 | H1001 | H1001 | H3025 | SRR7637550 | 9.8 |
| KAS_05ta | 4 | KAS | Alpine | H1001 | H1001 | H1001 | H1001 | SRR7637551 | 13.3 |
| KAS_06ta_fil | 4 | KAS | Alpine | H1001 | H1001 | H3010 | H4041 | SRR7637552 | NA |
| KAS_07ta_fil | 4 | KAS | Alpine | H1001 | H1001 | H1001 | H3010 | SRR7637553 | NA |
| KAS_08ta_fil | 4 | KAS | Alpine | H1001 | H1001 | H1001 | H4015 | SRR7637546 | NA |
| KEK_01da | 2 | KEK | Pannonian | H2027 | H4026 | | | SRR2040809 | NA |
| KLE_01ta | 4 | KLE | Ruderal | H1001 | H1001 | H1001 | H2003 | SRR8241415 | 16.1 |
| KLE_02ta | 4 | KLE | Ruderal | H1001 | H2008 | H3003 | H4002 | SRR8241413 | 12.4 |
| KLE_03ta | 4 | KLE | Ruderal | H1001 | H1001 | H2006 | H4015 | SRR8241418 | 18.6 |
| KLE_04ta | 4 | KLE | Ruderal | H1001 | H3003 | H3008 | H4004 | N/A | 13.0 |
| KLE_05ta | 4 | KLE | Ruderal | H1001 | H1001 | H1001 | H4002 | SRR8241411 | 16.1 |
| KLE_06ta | 4 | KLE | Ruderal | H1001 | H1001 | H3014 | H4001 | SRR8241412 | 13.3 |
| KLE_07ta | 4 | KLE | Ruderal | H1001 | H1001 | H3008 | H3014 | SRR8241414 | 12.4 |
| KLE_08ta | 4 | KLE | Ruderal | H1001 | H1001 | H2028 | H4001 | SRR8241416 | 11.4 |
| KLE_09ta | 4 | KLE | Ruderal | H1001 | H1001 | H1001 | H2006 | SRR8241417 | 18.5 |
| KOS_01ta | 4 | KOS | Alpine | H1001 | H1001 | H1001 | H2004 | SRR7637547 | 12.0 |
| KOS_02ta | 4 | KOS | Alpine | H1001 | H1001 | H1001 | H2010 | SRR7637548 | 8.2 |
| KOS_03ta | 4 | KOS | Alpine | H1001 | H1001 | H2004 | H3002 | SRR7637549 | 18.3 |
| KOS_04ta | 4 | KOS | Alpine | H1001 | H1001 | H1001 | H2001 | SRR7637542 | 12.6 |
| KOS_05ta | 4 | KOS | Alpine | H1001 | H2004 | H3004 | H4010 | SRR7637543 | 10.9 |
| KOS_06ta | 4 | KOS | Alpine | H1001 | H1001 | H2006 | H4002 | SRR7637294 | 14.3 |
| KOS_07ta | 4 | KOS | Alpine | H1001 | H1001 | H1001 | H1001 | SRR7637293 | 13.2 |
| KOW_01ta | 4 | KOW | Ruderal | H1001 | H2007 | H2028 | H2028 | SRR7637292 | 10.5 |
| KOW_02ta | 4 | KOW | Ruderal | H1001 | H1001 | H4040 | H0000 | SRR7637291 | 9.9 |
| KOW_03ta | 4 | KOW | Ruderal | H1001 | H2007 | H3001 | H3001 | SRR7637298 | 13.0 |
| KOW_04ta | 4 | KOW | Ruderal | H1001 | H1001 | H2028 | H4040 | SRR7637297 | 16.0 |
| KOW_05ta | 4 | KOW | Ruderal | H2028 | H3001 | H3001 | H4009 | SRR7637296 | 17.7 |
| KOW_06ta | 4 | KOW | Ruderal | H1001 | H1001 | H2007 | H3009 | SRR7637295 | 15.4 |
| KOW_07ta | 4 | KOW | Ruderal | H1001 | H1001 | H2007 | H3001 | SRR7637286 | 14.7 |
| KOW_08ta | 4 | KOW | Ruderal | H1001 | H1001 | H1001 | H3001 | SRR7637285 | 9.2 |
| KPS_01da | 2 | KPS | WCarpathian | H2007 | H4040 | | | SRR2040799 | NA |
| KPS_02da | 2 | KPS | WCarpathian | H2004 | H4029 | | | SRR3111447 | NA |
| KPS_03da | 2 | KPS | WCarpathian | H3003 | H3006 | | | SRR2040801 | NA |
| KPT_01ta | 4 | KPT | Hercynian | H2004 | H2004 | H2006 | H3009 | SRR2040823 | NA |
| KRM_01da | 2 | KRM | Baltic | H4037 | H4038 | | | SRR23281929 | 24.4 |
| KRM_02da | 2 | KRM | Baltic | H1001 | H2006 | | | SRR23281928 | 20.7 |
| KRM_03da | 2 | KRM | Baltic | H1001 | H2001 | | | SRR23281927 | 14.6 |
| KRM_04da | 2 | KRM | Baltic | H2026 | H4004 | | | SRR23281926 | 20.7 |
| KRM_05da | 2 | KRM | Baltic | H2003 | H2007 | | | SRR23281925 | 21.1 |
| KRM_06da | 2 | KRM | Baltic | H4004 | H4030 | | | SRR23282199 | 16.9 |
| KRM_07da | 2 | KRM | Baltic | H2006 | H3001 | | | SRR23282198 | 18.0 |
| KRM_08da | 2 | KRM | Baltic | H1001 | H3003 | | | SRR23282197 | 17.7 |
| KZL_01da | 2 | KZL | Pannonian | H2022 | H4030 | | | SRR7637337 | NA |
| KZL_03da | 2 | KZL | Pannonian | H1001 | H3008 | | | SRR7637338 | NA |
| KZL_05da | 2 | KZL | Pannonian | H3002 | H4023 | | | SRR7637335 | NA |
| LAC_01ta | 4 | LAC | SCarpathian | H1001 | H1001 | H2001 | H4004 | SRR7637334 | 12.2 |
| LAC_02ta | 4 | LAC | SCarpathian | H1001 | H1001 | H1001 | H3001 | SRR7637331 | 14.8 |
| LAC_03ta | 4 | LAC | SCarpathian | H1001 | H1001 | H1001 | H4037 | SRR7637332 | 14.0 |
| LAC_04ta | 4 | LAC | SCarpathian | H1001 | H1001 | H3009 | H4019 | SRR7637341 | 18.0 |
| LAC_05ta | 4 | LAC | SCarpathian | H1001 | H2001 | H3004 | H4031 | SRR7637342 | 15.2 |
| LAC_06ta | 4 | LAC | SCarpathian | H1001 | H1001 | H4003 | H4015 | SRR7637382 | 17.1 |
| LAC_07ta | 4 | LAC | SCarpathian | H1001 | H1001 | H1001 | H3003 | SRR7637381 | 18.6 |
| LAC_08ta | 4 | LAC | SCarpathian | H1001 | H1001 | H1001 | H2007 | SRR7637384 | 14.9 |
| LTW_01ta | 4 | LTW | Ruderal | H1001 | H1001 | H1001 | H3008 | SRR2040811 | NA |
| LUB_01ta | 4 | LUB | Hercynian | H1001 | H1001 | H1001 | H4011 | SRR2040820 | NA |
| MAR_01ta | 4 | MAR | Alpine | H1001 | H2008 | H4015 | H4019 | SRR2040824 | NA |
| MIA_02ta | 4 | MIA | Ruderal-WCarp mix | H1001 | H1001 | H2007 | H4005 | SRR8241453 | 15.3 |
| MIA_03ta | 4 | MIA | Ruderal-WCarp mix | H1001 | H1001 | H1001 | H1001 | SRR8241450 | 16.1 |
| MIA_04ta | 4 | MIA | Ruderal-WCarp mix | H1001 | H1001 | H2007 | H4030 | SRR8241452 | 12.5 |
| MIA_05ta | 4 | MIA | Ruderal-WCarp mix | H1001 | H1001 | H1001 | H2026 | SRR8241424 | 10.9 |
| MIA_06ta | 4 | MIA | Ruderal-WCarp mix | H1001 | H1001 | H1001 | H1001 | SRR8241434 | 11.6 |
| MIA_07ta | 4 | MIA | Ruderal-WCarp mix | H1001 | H1001 | H2005 | H2007 | SRR8241433 | 10.9 |
| MIA_09ta | 4 | MIA | Ruderal-WCarp mix | H1001 | H1001 | H2005 | H3004 | SRR8241423 | 13.0 |
| MIE_01da | 2 | MIE | Baltic | H1001 | H3002 | | | SRR7637383 | 17.2 |
| MIE_02da | 2 | MIE | Baltic | H2007 | H4012 | | | SRR7637378 | 14.2 |
| MIE_03da | 2 | MIE | Baltic | H4021 | H4023 | | | SRR7637377 | 15.3 |
| MIE_04da | 2 | MIE | Baltic | H1001 | H4023 | | | SRR7637380 | 10.5 |
| MIE_05da | 2 | MIE | Baltic | H2006 | H4016 | | | SRR7637379 | 12.5 |
| MIE_06da | 2 | MIE | Baltic | H2007 | H4015 | | | SRR7637386 | 15.1 |

| | | | | | | | | | |
|---|---|---|---|---|---|---|---|---|---|
| MIE_07da | 2 | MIE | Baltic | H3004 | H4026 | | | SRR7637385 | 18.4 |
| MIE_08da | 2 | MIE | Baltic | H2006 | H3004 | | | SRR7637410 | 15.5 |
| MIE_09da | 2 | MIE | Baltic | H2007 | H4003 | | | SRR23281941 | 16.5 |
| MIE_10da | 2 | MIE | Baltic | H2007 | H3002 | | | SRR23281940 | 9.6 |
| MIE_11da | 2 | MIE | Baltic | H3004 | H4003 | | | SRR23281939 | 15.4 |
| OPP_01ta | 4 | OPP | Alpine | H1001 | H1001 | H2007 | H3014 | SRR12778320 | 17.4 |
| OPP_02ta | 4 | OPP | Alpine | H1001 | H2007 | H3014 | H3025 | SRR12778319 | 17.3 |
| OPP_03ta | 4 | OPP | Alpine | H1001 | H4004 | H4019 | H4042 | SRR12778318 | 17.7 |
| OPP_04ta | 4 | OPP | Alpine | H2008 | H4004 | H4019 | H4027 | SRR12778317 | 18.1 |
| OPP_05ta | 4 | OPP | Alpine | H1001 | H1001 | H4027 | H4027 | SRR12778316 | 20.9 |
| OPP_06ta | 4 | OPP | Alpine | H1001 | H1001 | H1001 | H4015 | SRR12778315 | 19.1 |
| OPP_07ta | 4 | OPP | Alpine | H1001 | H2008 | H3012 | H0000 | SRR12778314 | 14.9 |
| OPP_08ta | 4 | OPP | Alpine | H1001 | H2009 | H3012 | H4004 | SRR12778313 | 17.9 |
| OSO_01da | 2 | OSO | WCarpathian | H2010 | H4001 | | | SRR2040800 | NA |
| PAD_01da | 2 | PAD | ECarpathian_2x | H1001 | H4001 | | | SRR23282117 | 34.8 |
| PAD_02da | 2 | PAD | ECarpathian_2x | H2007 | H3012 | | | SRR23282116 | 15.7 |
| PAD_03da | 2 | PAD | ECarpathian_2x | H1001 | H1001 | | | SRR23282115 | 30.8 |
| PAD_04da | 2 | PAD | ECarpathian_2x | H1001 | H1001 | | | SRR23282114 | 36.2 |
| PAD_05da | 2 | PAD | ECarpathian_2x | H2007 | H3025 | | | SRR23282113 | 28.6 |
| PAD_06da | 2 | PAD | ECarpathian_2x | H1001 | H4042 | | | SRR23282112 | 29.0 |
| PAD_07da | 2 | PAD | ECarpathian_2x | H2027 | H3004 | | | SRR23282111 | 27.1 |
| PAD_08da | 2 | PAD | ECarpathian_2x | H2021 | H3011 | | | SRR23282110 | 22.4 |
| PAD_09da | 2 | PAD | ECarpathian_2x | H2010 | H4040 | | | SRR23282108 | 31.7 |
| PAT_01ta | 4 | PAT | ECarpathian | H1001 | H2009 | H3002 | H4027 | SRR23282107 | 15.1 |
| PAT_02ta | 4 | PAT | ECarpathian | H2024 | H2009 | H3002 | H3011 | SRR23282106 | 28.8 |
| PAT_03ta | 4 | PAT | ECarpathian | H1001 | H1001 | H2009 | H3007 | SRR23282105 | 23.1 |
| PAT_04ta | 4 | PAT | ECarpathian | H1001 | H1001 | H3002 | H4039 | SRR23282104 | 18.2 |
| PAT_05ta | 4 | PAT | ECarpathian | H1001 | H1001 | H3002 | H4037 | SRR23282103 | 32.9 |
| PAT_06ta | 4 | PAT | ECarpathian | H1001 | H1001 | H1001 | H4015 | SRR23282102 | 26.7 |
| PAT_07ta | 4 | PAT | ECarpathian | H1001 | H4006 | H4025 | H4027 | SRR23282101 | 24.0 |
| PER_01ta | 4 | PER | Alpine | H3002 | H3009 | H3021 | H4005 | SRR12778312 | 19.5 |
| PER_02ta | 4 | PER | Alpine | H1001 | H1001 | H1001 | H3014 | SRR12778311 | 18.5 |
| PER_03ta | 4 | PER | Alpine | H1001 | H1001 | H1001 | H0000 | SRR12778309 | 16.5 |
| PER_04ta | 4 | PER | Alpine | H1001 | H2003 | H2007 | H3010 | SRR12778308 | 23.7 |
| PER_05ta | 4 | PER | Alpine | H1001 | H1001 | H1001 | H2007 | SRR12778307 | 44.1 |
| PER_06ta | 4 | PER | Alpine | H1001 | H1001 | H1001 | H1001 | SRR12778306 | 18.5 |
| PER_07ta | 4 | PER | Alpine | H1001 | H1001 | H1001 | H3010 | SRR12778305 | 19.2 |
| PER_08ta | 4 | PER | Alpine | H1001 | H1001 | H1001 | H4043 | SRR12778304 | 24.0 |
| PHD_01da | 2 | PHD | WCarpathian | H2007 | H4027 | | | SRR23282100 | 32.5 |
| PHD_02da | 2 | PHD | WCarpathian | H3008 | H4027 | | | SRR23282099 | 27.0 |
| PHD_03da | 2 | PHD | WCarpathian | H2003 | H4014 | | | SRR23282097 | 23.7 |
| PHD_04da | 2 | PHD | WCarpathian | H2004 | H3001 | H3011 | | SRR23282096 | 26.9 |
| PHD_05da | 2 | PHD | WCarpathian | H1001 | H2009 | | | SRR23282095 | 16.9 |
| PHD_06da | 2 | PHD | WCarpathian | H2007 | H4023 | | | SRR23282094 | 19.8 |
| PHD_07da | 2 | PHD | WCarpathian | H2024 | H3011 | | | SRR23282093 | 24.7 |
| PHD_08da | 2 | PHD | WCarpathian | H3007 | H3021 | | | SRR23282061 | 26.0 |
| PHT_01ta | 4 | PHT | WCarpathian | H1001 | H1001 | H1001 | H2005 | SRR23282059 | 31.6 |
| PHT_02ta | 4 | PHT | WCarpathian | H2006 | H2008 | H4002 | H4011 | SRR23282058 | 24.3 |
| PHT_03ta | 4 | PHT | WCarpathian | H1001 | H1001 | H2009 | H4005 | SRR23282057 | 19.3 |
| PHT_04ta | 4 | PHT | WCarpathian | H2006 | H3008 | H3014 | H4003 | SRR23282056 | 42.5 |
| PHT_05ta | 4 | PHT | WCarpathian | H1001 | H1001 | H2007 | H4038 | SRR23282054 | 24.7 |
| PHT_06ta | 4 | PHT | WCarpathian | H2024 | H2024 | H2003 | H4010 | SRR23282053 | 33.7 |
| PHT_07ta | 4 | PHT | WCarpathian | H1001 | H1001 | H1001 | H1001 | SRR23282052 | 28.5 |
| PHT_08ta | 4 | PHT | WCarpathian | H1001 | H1001 | H1001 | H4026 | SRR23282051 | 26.1 |
| PRE_01da | 2 | PRE | Baltic | H1001 | H4004 | | | SRR7637411 | 20.1 |
| PRE_02da | 2 | PRE | Baltic | H1001 | H4003 | | | SRR7637412 | 20.8 |
| PRE_03da | 2 | PRE | Baltic | H4004 | H4026 | | | SRR7637413 | 29.1 |
| PRE_04da | 2 | PRE | Baltic | H1001 | H4009 | | | SRR7637414 | 15.7 |
| PRE_05da | 2 | PRE | Baltic | H4001 | H4030 | | | SRR7637415 | 19.0 |
| PRE_06da | 2 | PRE | Baltic | H1001 | H4010 | | | SRR7637416 | 16.0 |
| PRE_07da | 2 | PRE | Baltic | H2007 | H4039 | | | SRR7637417 | 23.7 |
| PRE_08da | 2 | PRE | Baltic | H3025 | H4029 | | | SRR7637418 | 16.5 |
| PUP_01da | 2 | PUP | WCarpathian | H3008 | H4008 | | | SRR3111436 | NA |
| PVO_01ta | 4 | PVO | WCarpathian | H1001 | H1001 | H4026 | H4034 | SRR2040832 | NA |
| RFT_01ta | 4 | RFT | Swabian | H1001 | H2006 | H2008 | H4041 | SRR7637419 | 9.8 |
| RFT_02ta | 4 | RFT | Swabian | H1001 | H1001 | H1001 | H1001 | SRR7637445 | 15.9 |
| RFT_03ta | 4 | RFT | Swabian | H1001 | H1001 | H1001 | H1001 | SRR7637444 | 12.4 |
| RFT_04ta | 4 | RFT | Swabian | H1001 | H1001 | H2004 | H4034 | SRR7637443 | 12.3 |
| RFT_05ta | 4 | RFT | Swabian | H1001 | H1001 | H1001 | H2008 | SRR7637442 | 13.6 |
| RFT_06ta | 4 | RFT | Swabian | H1001 | H1001 | H3009 | H3009 | SRR7637440 | 12.2 |
| RFT_07ta | 4 | RFT | Swabian | H1001 | H1001 | H1001 | H4035 | SRR7637439 | 16.5 |
| RFT_09ta | 4 | RFT | Swabian | H1001 | H3008 | H3009 | H4015 | SRR7637437 | 8.5 |
| RFT_10ta | 4 | RFT | Swabian | H1001 | H1001 | H1001 | H1001 | SRR7637436 | 16.2 |

| | | | | | | | | | |
|---|---|---|---|---|---|---|---|---|---|
| RFT_11ta_fil | 4 | RFT | Swabian | H1001 | H2001 | H2024 | H3012 | SRR7637480 | NA |
| RZA_01da | 2 | RZA | ECarpathian_2x | H1001 | H3006 | | | SRR7637481 | 10.1 |
| RZA_02da | 2 | RZA | ECarpathian_2x | H2026 | H4030 | | | SRR7637478 | 10.5 |
| RZA_03da | 2 | RZA | ECarpathian_2x | H1001 | H4003 | | | SRR7637479 | 10.7 |
| RZA_04da | 2 | RZA | ECarpathian_2x | H3022 | H4010 | | | SRR7637484 | 9.2 |
| RZA_05da | 2 | RZA | ECarpathian_2x | H4001 | H4024 | | | SRR7637485 | 13.1 |
| RZA_06da | 2 | RZA | ECarpathian_2x | H2008 | H3010 | | | SRR7637482 | 10.3 |
| RZA_08da | 2 | RZA | ECarpathian_2x | H2028 | H3001 | | | SRR7637472 | 9.3 |
| RZA_09da | 2 | RZA | ECarpathian_2x | H1001 | H4039 | | | SRR7637473 | 9.1 |
| SCH_03ta | 4 | SCH | Alpine | H1001 | H1001 | H3012 | H4029 | SRR7637507 | NA |
| SCH_04ta | 4 | SCH | Alpine | H1001 | H4015 | H4029 | H4035 | SRR7637506 | NA |
| SCH_05ta | 4 | SCH | Alpine | H1001 | H1001 | H1001 | H4015 | SRR7637509 | NA |
| SCH_06ta | 4 | SCH | Alpine | H1001 | H2026 | H4015 | H4029 | SRR7637508 | NA |
| SCH_07ta | 4 | SCH | Alpine | H1001 | H4015 | H4015 | H4029 | SRR7637511 | NA |
| SNO_01da | 2 | SNO | WCarpathian | H4017 | H4027 | | | SRR7637512 | NA |
| SNO_02da | 2 | SNO | WCarpathian | H1001 | H2018 | | | SRR7637499 | NA |
| SNO_03da_f | 2 | SNO | WCarpathian | H2021 | H3005 | | | SRR7637498 | NA |
| SPI_01ta | 4 | SPI | WCarpathian | H1001 | H2007 | H3006 | H4042 | SRR7637533 | 10.6 |
| SPI_02ta | 4 | SPI | WCarpathian | H1001 | H1001 | H3009 | H3009 | SRR7637534 | 13.9 |
| SPI_03ta | 4 | SPI | WCarpathian | H1001 | H1001 | H3003 | H3009 | SRR7637527 | 13.1 |
| SPI_04ta | 4 | SPI | WCarpathian | H1001 | H1001 | H4004 | H3001 | SRR7637528 | 12.5 |
| SPI_05ta | 4 | SPI | WCarpathian | H1001 | H2029 | H4001 | H4030 | SRR7637529 | 14.3 |
| SPI_06ta | 4 | SPI | WCarpathian | H1001 | H1001 | H3003 | H3009 | SRR7637530 | 16.3 |
| SPI_07ta | 4 | SPI | WCarpathian | H1001 | H1001 | H1001 | H3010 | SRR7637535 | 13.2 |
| SPI_08ta | 4 | SPI | WCarpathian | H1001 | H1001 | H1001 | H3010 | SRR7637536 | 13.8 |
| SPI_09ta | 4 | SPI | WCarpathian | H1001 | H1001 | H3006 | H3015 | SRR7637276 | 16.7 |
| SPI_10ta | 4 | SPI | WCarpathian | H1001 | H1001 | H1001 | H4041 | SRR7637275 | 12.6 |
| SPI_11ta | 4 | SPI | WCarpathian | H1001 | H1001 | H3024 | H4034 | SRR7637274 | 12.3 |
| SPI_12ta | 4 | SPI | WCarpathian | H1001 | H1001 | H2006 | H2007 | SRR7637273 | 11.9 |
| SPI_13ta | 4 | SPI | WCarpathian | H1001 | H2007 | H4023 | H4027 | SRR7637280 | 14.5 |
| SRL_01ta | 4 | SRL | Ruderal | H1001 | H1001 | H1001 | H1001 | SRR2040812 | NA |
| SSP_01da | 2 | SSP | Pannonian | H1001 | H4031 | | | SRR2040807 | NA |
| STE_01ta | 4 | STE | Ruderal | H1001 | H1001 | H1001 | H3004 | SRR7637279 | 13.8 |
| STE_02ta | 4 | STE | Ruderal | H1001 | H1001 | H1001 | H3006 | SRR7637278 | 13.1 |
| STE_03ta | 4 | STE | Ruderal | H1001 | H1001 | H1001 | H4023 | SRR7637277 | 14.7 |
| STE_04ta | 4 | STE | Ruderal | H1001 | H1001 | H1001 | H1001 | SRR7637282 | 15.2 |
| STE_05ta | 4 | STE | Ruderal | H1001 | H1001 | H2006 | H0000 | SRR7637281 | 14.2 |
| STE_06ta | 4 | STE | Ruderal | H1001 | H1001 | H2008 | H4023 | SRR7637365 | 14.3 |
| STE_07ta | 4 | STE | Ruderal | H1001 | H2008 | H2024 | H3004 | SRR7637362 | 13.9 |
| STE_08ta | 4 | STE | Ruderal | H1001 | H1001 | H3002 | H3003 | SRR7637435 | 11.3 |
| STG_01ta | 4 | STG | Hercynian | H1001 | H1001 | H2003 | H4011 | SRR12778303 | 18.5 |
| STG_02ta | 4 | STG | Hercynian | H1001 | H1001 | H1001 | H1001 | SRR12778302 | 19.8 |
| STG_03ta | 4 | STG | Hercynian | H1001 | H1001 | H2005 | H2010 | SRR12778301 | 17.7 |
| STG_04ta | 4 | STG | Hercynian | H1001 | H1001 | H3009 | H4035 | SRR12778300 | 17.4 |
| STG_05ta | 4 | STG | Hercynian | H1001 | H1001 | H2004 | H4002 | SRR12778298 | 17.8 |
| STG_06ta | 4 | STG | Hercynian | H1001 | H1001 | H2001 | H3009 | SRR12778297 | 17.6 |
| STG_07ta | 4 | STG | Hercynian | H1001 | H1001 | H2009 | H2028 | SRR12778296 | 11.4 |
| STG_08ta | 4 | STG | Hercynian | H1001 | H2026 | H3003 | H4001 | SRR12778295 | 16.0 |
| SUB_01da | 2 | SUB | WCarpathian | H2022 | H3025 | | | SRR23469345 | 78.2 |
| SUB_02da | 2 | SUB | WCarpathian | H2009 | H4008 | | | SRR23469343 | 69.1 |
| SUB_03da | 2 | SUB | WCarpathian | H2028 | H4034 | | | SRR23469341 | 64.8 |
| SUB_04da | 2 | SUB | WCarpathian | H4016 | H4043 | | | SRR23469339 | 84.8 |
| SUB_05da | 2 | SUB | WCarpathian | H2010 | H2021 | | | SRR10549920 | 97.7 |
| SUB_06da | 2 | SUB | WCarpathian | H2026 | H4010 | | | SRR10549909 | 86.2 |
| SUB_07da | 2 | SUB | WCarpathian | H1001 | H4023 | | | SRR23469337 | 72.1 |
| SUB_08da | 2 | SUB | WCarpathian | H1001 | H2010 | | | SRR23469335 | 97.5 |
| SUB_10da | 2 | SUB | WCarpathian | H1001 | H3023 | | | SRR10550007 | 100.9 |
| SUB_11da | 2 | SUB | WCarpathian | H1001 | H2004 | | | SRR10549996 | 114.6 |
| SUB_12da | 2 | SUB | WCarpathian | H2001 | H2006 | | | SRR10549994 | 92.1 |
| SUB_13da | 2 | SUB | WCarpathian | H3003 | H4016 | | | SRR23469395 | 84.4 |
| SUB_14da | 2 | SUB | WCarpathian | H2004 | H3003 | | | SRR10549993 | 103.4 |
| SUB_15da | 2 | SUB | WCarpathian | H2005 | H3003 | | | SRR10549992 | 87.5 |
| SUB_16da | 2 | SUB | WCarpathian | H2028 | H4026 | | | SRR23469393 | 73.9 |
| SUB_17da | 2 | SUB | WCarpathian | H2009 | H2010 | | | SRR23469391 | 53.6 |
| SUB_18da | 2 | SUB | WCarpathian | H1001 | H4039 | | | SRR10549991 | 88.6 |
| SWA_01ta | 4 | SWA | Swabian | H1001 | H1001 | H1001 | H1001 | SRR7637434 | 17.9 |
| SWA_02ta | 4 | SWA | Swabian | H1001 | H1001 | H2006 | H2006 | SRR7637433 | 16.5 |
| SWA_03ta | 4 | SWA | Swabian | H1001 | H1001 | H2006 | H0000 | SRR7637432 | 9.6 |
| SWA_04ta | 4 | SWA | Swabian | H1001 | H1001 | H1001 | H2007 | SRR7637431 | 9.5 |
| SWA_05ta | 4 | SWA | Swabian | H1001 | H1001 | H1001 | H3004 | SRR7637430 | 14.1 |
| SWA_07ta | 4 | SWA | Swabian | H1001 | H1001 | H2007 | H4018 | SRR7637389 | 11.6 |
| SWA_08ta | 4 | SWA | Swabian | H1001 | H1001 | H1001 | H1001 | SRR7637402 | 7.6 |
| SWD_01da | 2 | SWD | Baltic | H2028 | H4005 | | | SRR23281938 | 8.8 |

| | | | | | | | | | |
|---|---|---|---|---|---|---|---|---|---|
| SWD_02da | 2 | SWD | Baltic | H4010 | H4022 | | | SRR23281937 | 15.3 |
| SWD_03da | 2 | SWD | Baltic | H3011 | H4010 | | | SRR23281936 | 8.2 |
| SWD_04da | 2 | SWD | Baltic | H2028 | H4010 | | | SRR23281934 | 14.2 |
| SWD_05da | 2 | SWD | Baltic | H3002 | H4022 | | | SRR23281933 | 18.7 |
| SZI_02da | 2 | SZI | Pannonian | H1001 | H4020 | | | SRR7637405 | NA |
| SZI_03da | 2 | SZI | Pannonian | H1001 | H4025 | | | SRR7637406 | NA |
| SZI_04da | 2 | SZI | Pannonian | H1001 | H4029 | | | SRR7637407 | NA |
| SZI_05da | 2 | SZI | Pannonian | H1001 | H3012 | | | SRR7637408 | NA |
| TBG_01ta_fi | 4 | TBG | Ruderal | H1001 | H1001 | H1001 | H1001 | SRR7637394 | NA |
| TBG_03ta | 4 | TBG | Ruderal | H1001 | H1001 | H1001 | H1001 | SRR7637395 | NA |
| TBG_04ta | 4 | TBG | Ruderal | H1001 | H1001 | H1001 | H3001 | SRR7637494 | NA |
| TBG_05ta | 4 | TBG | Ruderal | H1001 | H1001 | H1001 | H1001 | SRR7637490 | NA |
| TIS_01ta | 4 | TIS | SCarpathian | H1001 | H1001 | H2006 | H4041 | SRR10549949 | 23.4 |
| TIS_02ta | 4 | TIS | SCarpathian | H1001 | H2004 | H4040 | H4041 | SRR10549950 | 42.7 |
| TIS_03ta | 4 | TIS | SCarpathian | H1001 | H1001 | H2024 | H3006 | SRR10549948 | 43.9 |
| TIS_04ta | 4 | TIS | SCarpathian | H1001 | H1001 | H4003 | H4025 | SRR10549947 | 23.7 |
| TIS_05ta | 4 | TIS | SCarpathian | H1001 | H2009 | H4039 | H4041 | SRR10549954 | 46.6 |
| TIS_06ta | 4 | TIS | SCarpathian | H1001 | H1001 | H4003 | H4040 | SRR10549951 | 45.7 |
| TIS_07ta | 4 | TIS | SCarpathian | H1001 | H1001 | H3023 | H4009 | SRR10549952 | 44.6 |
| TIS_08ta | 4 | TIS | SCarpathian | H3012 | H4017 | H4039 | H4041 | SRR10549955 | 49.5 |
| TKO_01ta | 4 | TKO | WCarpathian | H1001 | H1001 | H1001 | H4034 | SRR7637500 | 19.1 |
| TKO_02ta | 4 | TKO | WCarpathian | H1001 | H1001 | H1001 | H2003 | SRR7637503 | 20.0 |
| TKO_03ta | 4 | TKO | WCarpathian | H1001 | H2003 | H4002 | H4026 | SRR7637502 | 17.0 |
| TKO_04ta | 4 | TKO | WCarpathian | H1001 | H1001 | H2003 | H4023 | SRR7637505 | 18.1 |
| TKO_05ta | 4 | TKO | WCarpathian | H1001 | H2022 | H2024 | H3006 | SRR7637504 | 16.3 |
| TKO_06ta | 4 | TKO | WCarpathian | H1001 | H1001 | H2024 | H2028 | SRR7637396 | 18.7 |
| TKO_07ta | 4 | TKO | WCarpathian | H1001 | H1001 | H2007 | H2024 | SRR7637388 | 14.7 |
| TKO_08ta | 4 | TKO | WCarpathian | H1001 | H1001 | H2007 | H2009 | SRR7637554 | 18.8 |
| TRA_01ta | 4 | NA | Baltic | H1001 | H1001 | H1001 | H4043 | SRR2040822 | NA |
| TRD_03da | 2 | NA | Baltic | H3021 | H4043 | | | SRR7637560 | NA |
| TRD_04da | 2 | NA | Baltic | H3012 | H4014 | | | SRR7637470 | NA |
| TRD_05da | 2 | NA | Baltic | H2005 | H4015 | | | SRR7637471 | NA |
| TRD_06da | 2 | NA | Baltic | H1001 | H3023 | | | SRR7637476 | NA |
| TRE_01ta | 4 | TRE | WCarpathian | H1001 | H1001 | H1001 | H2028 | SRR7637475 | 24.7 |
| TRE_02ta | 4 | TRE | WCarpathian | H1001 | H1001 | H1001 | H4001 | SRR7637283 | 16.3 |
| TRE_03ta | 4 | TRE | WCarpathian | H1001 | H1001 | H2001 | H4029 | SRR7637289 | 16.2 |
| TRE_04ta | 4 | TRE | WCarpathian | H1001 | H2027 | H4003 | H4015 | SRR7637400 | 15.5 |
| TRE_05ta | 4 | TRE | WCarpathian | H1001 | H1001 | H2022 | H4008 | SRR7637348 | 16.8 |
| TRE_06ta | 4 | TRE | WCarpathian | H1001 | H1001 | H3028 | H4001 | SRR7637284 | 15.1 |
| TRE_08ta | 4 | TRE | WCarpathian | H1001 | H1001 | H2028 | H3028 | SRR7637290 | 14.0 |
| TRT_05ta | 4 | TRT | WCarpathian | H1001 | H2004 | H3015 | H4017 | SRR7637288 | NA |
| TRT_06ta | 4 | TRT | WCarpathian | H1001 | H1001 | H2005 | H3025 | SRR7637287 | NA |
| TRT_07ta | 4 | TRT | WCarpathian | H1001 | H2010 | H3003 | H3021 | SRR7637427 | NA |
| TRT_08ta | 4 | TRT | WCarpathian | H1001 | H2007 | H4008 | H4041 | SRR7637424 | NA |
| TRT_09ta | 4 | TRT | WCarpathian | H1001 | H2010 | H3003 | H3028 | SRR7637387 | NA |
| TZI_01ta | 4 | TZI | SCarpathian | H1001 | H1001 | H1001 | H4003 | SRR7637545 | 11.6 |
| TZI_02ta | 4 | TZI | SCarpathian | H1001 | H2025 | H4005 | H4012 | SRR7637538 | 8.3 |
| TZI_03ta | 4 | TZI | SCarpathian | H1001 | H1001 | H2026 | H4002 | SRR7637539 | 9.2 |
| TZI_05ta | 4 | TZI | SCarpathian | H1001 | H1001 | H1001 | H1001 | SRR7637541 | 9.9 |
| TZI_06ta | 4 | TZI | SCarpathian | H1001 | H1001 | H1001 | H3025 | SRR7637497 | 8.7 |
| TZI_07ta | 4 | TZI | SCarpathian | H2024 | H3001 | H3001 | H4011 | SRR7637561 | 8.8 |
| TZI_08ta | 4 | TZI | SCarpathian | H1001 | H2005 | H3015 | H4012 | SRR7637374 | 7.5 |
| TZI_09ta | 4 | TZI | SCarpathian | H1001 | H1001 | H1001 | H1001 | SRR7637373 | 8.7 |
| TZI_10ta | 4 | TZI | SCarpathian | H1001 | H2006 | H3001 | H3002 | SRR2040834 | 10.4 |
| VEL_01da | 2 | VEL | WCarpathian | H2006 | H4039 | | | SRR7637376 | 13.6 |
| VEL_02da | 2 | VEL | WCarpathian | H1001 | H2003 | | | SRR7637375 | 15.3 |
| VEL_03da | 2 | VEL | WCarpathian | H2006 | H2007 | | | SRR7637370 | 13.4 |
| VEL_04da | 2 | VEL | WCarpathian | H3012 | H3024 | | | SRR7637369 | 13.2 |
| VEL_05da | 2 | VEL | WCarpathian | H2006 | H2019 | | | SRR7637372 | 14.6 |
| VEL_06da | 2 | VEL | WCarpathian | H2028 | H2028 | | | SRR7637371 | 15.1 |
| VEL_07da | 2 | VEL | WCarpathian | H1001 | H4039 | | | SRR7637368 | 15.0 |
| VEL_08da | 2 | VEL | WCarpathian | H3002 | H4003 | | | SRR7637367 | 19.2 |
| VEL_09da | 2 | VEL | WCarpathian | H3024 | H4043 | | | SRR3111444 | 21.2 |
| VID_02da | 2 | VID | SCarpathian_2x | H3010 | H4016 | | | SRR7637330 | 13.5 |
| VID_03da | 2 | VID | SCarpathian_2x | H2027 | H4021 | | | SRR7637327 | 11.2 |
| VID_04da | 2 | VID | SCarpathian_2x | H2026 | H3010 | | | SRR7637328 | 11.1 |
| VID_05da | 2 | VID | SCarpathian_2x | H3006 | H4024 | | | SRR7637325 | 13.2 |
| VID_06da | 2 | VID | SCarpathian_2x | H2011 | H2022 | | | SRR7637326 | 13.1 |
| VID_07da | 2 | VID | SCarpathian_2x | H2009 | H4034 | | | SRR7637323 | 14.0 |
| VID_08da | 2 | VID | SCarpathian_2x | H2009 | H3006 | | | SRR7637324 | 15.9 |
| VLA_01ta | 4 | VLA | Hercynian | H1001 | H2006 | H2007 | H4004 | SRR12778294 | 20.9 |
| VLA_02ta | 4 | VLA | Hercynian | H1001 | H1001 | H1001 | H3002 | SRR12778293 | 16.5 |
| VLA_03ta | 4 | VLA | Hercynian | H1001 | H2007 | H4004 | H4029 | SRR12778292 | 18.6 |

| | | | | | | | | | |
|---|---|---|---|---|---|---|---|---|---|
| VLA_04ta | 4 | VLA | Hercynian | H1001 | H1001 | H1001 | H4001 | SRR12778291 | 19.1 |
| VLA_05ta | 4 | VLA | Hercynian | H1001 | H1001 | H1001 | H2011 | SRR12778290 | 20.6 |
| VLA_06ta | 4 | VLA | Hercynian | H1001 | H1001 | H1001 | H2011 | SRR12778289 | 20.8 |
| VLA_07ta | 4 | VLA | Hercynian | H1001 | H1001 | H3012 | H3014 | SRR12778287 | 21.3 |
| VLA_08ta | 4 | VLA | Hercynian | H1001 | H1001 | H2007 | H4012 | SRR12778286 | 17.8 |
| VOR_01ta | 4 | VOR | Alpine | H1001 | H1001 | H1001 | H4009 | SRR12778285 | 21.9 |
| VOR_02ta | 4 | VOR | Alpine | H1001 | H1001 | H1001 | H1001 | SRR12778284 | 19.1 |
| VOR_03ta | 4 | VOR | Alpine | H1001 | H1001 | H1001 | H3001 | SRR12778283 | 20.3 |
| VOR_04ta | 4 | VOR | Alpine | H1001 | H2009 | H4001 | H4035 | SRR12778282 | 17.6 |
| VOR_05ta | 4 | VOR | Alpine | H1001 | H2001 | H4014 | H4035 | SRR12778281 | 14.4 |
| VOR_06ta | 4 | VOR | Alpine | H1001 | H1001 | H1001 | H2028 | SRR12778280 | 15.7 |
| VOR_07ta | 4 | VOR | Alpine | H1001 | H1001 | H1001 | H1001 | SRR12778279 | 17.8 |
| VOR_08ta | 4 | VOR | Alpine | H1001 | H1001 | H1001 | H4001 | SRR12778278 | 18.4 |
| VSD_01da | 2 | VSD | WCarpathian | H2008 | H2008 | | | SRR3111446 | NA |
| VTA_01ta | 4 | VTA | WCarpathian | H1001 | H1001 | H1001 | H1001 | SRR2040833 | NA |
| VYR_01ta | 4 | VYR | Ruderal | H1001 | H1001 | H1001 | H1001 | SRR23282196 | 18.1 |
| VYR_02ta | 4 | VYR | Ruderal | H3009 | H3021 | H4002 | H4009 | SRR23282195 | 20.5 |
| VYR_03ta | 4 | VYR | Ruderal | H1001 | H1001 | H1001 | H4030 | SRR23282194 | 22.8 |
| VYR_04ta | 4 | VYR | Ruderal | H1001 | H2022 | H2022 | H4030 | SRR23282193 | 28.1 |
| VYR_05ta | 4 | VYR | Ruderal | H1001 | H1001 | H1001 | H1001 | SRR23282192 | 22.5 |
| VYR_06ta | 4 | VYR | Ruderal | H1001 | H1001 | H1001 | H4030 | SRR23282191 | 20.5 |
| VYR_07ta | 4 | VYR | Ruderal | H1001 | H1001 | H1001 | H2022 | SRR23282190 | 16.4 |
| VYR_08ta | 4 | VYR | Ruderal | H1001 | H1001 | H1001 | H4030 | SRR23282188 | 19.5 |
| WEK_03ta | 4 | WEK | Hercynian | H2021 | H2021 | H2026 | H4010 | SRR7637321 | NA |
| WEK_06ta | 4 | WEK | Hercynian | H1001 | H1001 | H2009 | H3006 | SRR7637322 | NA |
| WIL_01ta | 4 | WIL | Alpine | H1001 | H1001 | H1001 | H4027 | SRR10549981 | 44.8 |
| WIL_02ta | 4 | WIL | Alpine | H1001 | H1001 | H1001 | H4011 | SRR10549980 | 43.2 |
| WIL_03ta | 4 | WIL | Alpine | H1001 | H2009 | H2009 | H2009 | SRR10549979 | 40.1 |
| WIL_04ta | 4 | WIL | Alpine | H1001 | H4015 | H4028 | H4041 | SRR10549978 | 51.8 |
| WIL_05ta | 4 | WIL | Alpine | H1001 | H1001 | H1001 | H2006 | SRR10549977 | 45.0 |
| WIL_06ta | 4 | WIL | Alpine | H1001 | H1001 | H2007 | H4027 | SRR10549976 | 49.1 |
| WIL_07ta | 4 | WIL | Alpine | H1001 | H2007 | H3025 | H4011 | SRR10549974 | 36.7 |
| WIL_08ta | 4 | WIL | Alpine | H1001 | H3025 | H4039 | H4040 | SRR10549973 | 38.3 |
| WMH_01ta | 4 | WMH | Swabian | H1001 | H2008 | H2010 | H3008 | SRR2040819 | NA |
| ZAP_01ta | 4 | ZAP | WCarpathian | H1001 | H1001 | H1001 | H2019 | SRR8241451 | 11.0 |
| ZAP_02ta | 4 | ZAP | WCarpathian | H1001 | H1001 | H1001 | H1001 | SRR7637451 | 7.5 |
| ZAP_03ta | 4 | ZAP | WCarpathian | H1001 | H1001 | H1001 | H2006 | SRR8241455 | 10.8 |
| ZAP_04ta | 4 | ZAP | WCarpathian | H1001 | H1001 | H1001 | H2006 | SRR7637450 | 10.8 |
| ZAP_06ta | 4 | ZAP | WCarpathian | H1001 | H1001 | H1001 | H3021 | SRR7637448 | 12.1 |
| ZAP_07ta | 4 | ZAP | WCarpathian | H1001 | H1001 | H2029 | H4011 | SRR8241422 | 10.8 |
| ZAP_08ta | 4 | ZAP | WCarpathian | H1001 | H1001 | H1001 | H2001 | SRR7637423 | 9.8 |
| ZEP_01da | 2 | ZEP | WCarpathian | H4005 | H4021 | | | SRR10550009 | 50.7 |
| ZEP_02da | 2 | ZEP | WCarpathian | H1001 | H1001 | | | SRR10550008 | 39.0 |
| ZEP_03da | 2 | ZEP | WCarpathian | H1001 | H4003 | | | SRR23469389 | 35.2 |
| ZEP_04da | 2 | ZEP | WCarpathian | H4018 | H4021 | | | SRR10549986 | 37.4 |
| ZEP_05da | 2 | ZEP | WCarpathian | H3014 | H4027 | | | SRR10549975 | 23.3 |
| ZEP_06da | 2 | ZEP | WCarpathian | H4011 | H2018 | | | SRR10549964 | 36.0 |
| ZEP_07da | 2 | ZEP | WCarpathian | H3022 | H4027 | | | SRR10549953 | 36.7 |
| ZEP_08da | 2 | ZEP | WCarpathian | H3003 | H4030 | | | SRR10549942 | 25.7 |
| ZEP_09da | 2 | ZEP | WCarpathian | H1001 | H2005 | | | SRR23469387 | 25.3 |
| ZEP_10da | 2 | ZEP | WCarpathian | H1001 | H4004 | | | SRR23469385 | 33.6 |
| ZEP_11da | 2 | ZEP | WCarpathian | H3004 | H4004 | | | SRR23469383 | 31.8 |
| ZEP_12da | 2 | ZEP | WCarpathian | H4004 | H4023 | | | SRR10549931 | 45.7 |
| ZEP_13da | 2 | ZEP | WCarpathian | H4004 | H4040 | | | SRR23469381 | 56.9 |
| ZID_1da | 2 | ZID | Dinaric | H1001 | H4035 | | | SRR23282050 | 21.3 |
| ZID_2da | 2 | ZID | Dinaric | H2003 | H2021 | | | SRR23282049 | 28.0 |
| ZID_3da | 2 | ZID | Dinaric | H1001 | H2003 | | | SRR23282048 | 26.1 |
| ZID_4da | 2 | ZID | Dinaric | H2018 | H4034 | | | SRR23282047 | 20.5 |
| ZID_5da | 2 | ZID | Dinaric | H2007 | H3009 | | | SRR23282046 | 27.0 |
| ZID_6da | 2 | ZID | Dinaric | H1001 | H2001 | | | SRR23282045 | 22.7 |
| ZID_8da | 2 | ZID | Dinaric | H3001 | H4007 | | | SRR23282042 | 28.5 |
| ZIT_1ta | 4 | ZIT | Alpine-Dinaric admix | H1001 | H1001 | H3001 | H3006 | SRR23282041 | 33.9 |
| ZIT_2ta | 4 | ZIT | Alpine-Dinaric admix | H1001 | H1001 | H1001 | H1001 | SRR23282040 | 26.0 |
| ZIT_3ta | 4 | ZIT | Alpine-Dinaric admix | H1001 | H2009 | H3001 | H4037 | SRR23282039 | 32.0 |
| ZIT_4ta | 4 | ZIT | Alpine-Dinaric admix | H1001 | H1001 | H2010 | H4023 | SRR23282038 | 27.5 |
| ZIT_5ta | 4 | ZIT | Alpine-Dinaric admix | H1001 | H1001 | H1001 | H4040 | SRR23282037 | 22.7 |
| ZIT_6ta | 4 | ZIT | Alpine-Dinaric admix | H1001 | H1001 | H2001 | H4038 | SRR23282036 | 28.0 |
| ZIT_7ta | 4 | ZIT | Alpine-Dinaric admix | H1001 | H1001 | H1001 | H4007 | SRR23282035 | 22.7 |
| ZIT_8ta | 4 | ZIT | Alpine-Dinaric admix | H1001 | H1001 | H4011 | H4042 | SRR23281948 | 15.0 |

*Sequences taken from the following references: Novikova et al. (2017); Monnahan et al. (2019); Konečná, et al. (2021); Bohutínská, et al. (2021)

**Table S2.** S-locus genotypes of 417 diploid and 138 tetraploid individuals from *Arabidopsis lyrata* , as obtained by the NGSgenotyp pipeline on whole-genome resequencing data obtained from Scott et al. (2024). Allele sequences are identified according to a Brassicaceae functional sequence group nomenclature, as determined based on sequence similarity with alleles from related Arabidopsis species (see Table S4 for correspondence with *A. lyrata* -specific S-allele IDs).

| Accession | Ploidy | Lineage | Regional Sample | allele #1 | allele #2 | allele #3 | allele #4 | ENA run | ENA biosample |
|---|---|---|---|---|---|---|---|---|---|
| al1_1_4n | 4 | Central_Siberia_4n | 4x_Siberia | H1001 | H1001 | H2009 | H4010 | ERR12736014 | ERS18382727 |
| al2_1_4n | 4 | Central_Siberia_4n | 4x_Siberia | H1001 | H1001 | H3011 | H4034 | ERR12736043 | ERS18382728 |
| al3_1_4n | 4 | Central_Siberia_4n | 4x_Siberia | H1001 | H3011 | H4023 | H4037 | ERR12736039 | ERS18382729 |
| al4_1_4n | 4 | Central_Siberia_4n | 4x_Siberia | H1001 | H1001 | H2009 | H4015 | ERR12736034 | ERS18382730 |
| BAM_12.1-1 | 2 | East_Siberia_2n | 2x_Siberia | H4014 | H4030 | | | ERR12736064 | ERS18382731 |
| ERR3397904 | 2 | Scandinavia_UK_2n | | H3002 | H4004 | | | ERR3397904 | SAMEA5738018 |
| ERR3397905 | 2 | Scandinavia_UK_2n | | H1001 | H3002 | | | ERR3397905 | SAMEA5738019 |
| ERR3397906 | 2 | Scandinavia_UK_2n | | H1001 | H4016 | | | ERR3397906 | SAMEA5738020 |
| ERR3397907 | 2 | Scandinavia_UK_2n | | H3002 | H4034 | | | ERR3397907 | SAMEA5738021 |
| ERR3397908 | 2 | Scandinavia_UK_2n | | H4004 | H4020 | | | ERR3397908 | SAMEA5738022 |
| ERR3397909 | 2 | Scandinavia_UK_2n | | H1001 | H3002 | | | ERR3397909 | SAMEA5738023 |
| ERR3397910 | 2 | Scandinavia_UK_2n | | H2007 | H4004 | | | ERR3397910 | SAMEA5738024 |
| ERR3397911 | 2 | Scandinavia_UK_2n | | H1001 | H1001 | | | ERR3397911 | SAMEA5738025 |
| ERR3397912 | 2 | Scandinavia_UK_2n | | H1001 | H4029 | | | ERR3397912 | SAMEA5738026 |
| ERR3397913 | 2 | Scandinavia_UK_2n | | H1001 | H4004 | | | ERR3397913 | SAMEA5738027 |
| ERR3397914 | 2 | Scandinavia_UK_2n | | H4001 | H4022 | | | ERR3397914 | SAMEA5738028 |
| ERR3514858 | 4 | Central_Europe_4n | 4x_Europe | H1001 | H1001 | H1001 | H2007 | ERR3514858 | SAMEA5947843 |
| ERR3514859 | 4 | Central_Europe_4n | 4x_Europe | H1001 | H1001 | H2007 | H4026 | ERR3514859 | SAMEA5947844 |
| ERR3514860 | 4 | Central_Europe_4n | 4x_Europe | H1001 | H1001 | H1001 | H4026 | ERR3514860 | SAMEA5947845 |
| ERR3514864 | 4 | Central_Europe_4n | 4x_Europe | H2008 | H2008 | H1001 | - | ERR3514864 | SAMEA5947849 |
| ERR3514865 | 4 | Central_Europe_4n | 4x_Europe | H1001 | H2008 | H4004 | H4021 | ERR3514865 | SAMEA5947850 |
| ERR3514866 | 4 | Central_Europe_4n | 4x_Europe | H1001 | H1001 | H1001 | H4029 | ERR3514866 | SAMEA5947851 |
| ERR3514869 | 4 | Central_Europe_4n | 4x_Europe | H1001 | H1001 | H1001 | H4012 | ERR3514869 | SAMEA5947854 |
| ERR3514870 | 4 | Central_Europe_4n | 4x_Europe | H1001 | H1001 | H1001 | H1001 | ERR3514870 | SAMEA5947855 |
| ERR3514871 | 4 | Central_Europe_4n | 4x_Europe | H1001 | H1001 | H1001 | H1001 | ERR3514871 | SAMEA5947856 |
| ERR3514872 | 4 | Central_Europe_4n | 4x_Europe | H1001 | H1001 | H2006 | H2006 | ERR3514872 | SAMEA5947857 |
| ERR3514873 | 4 | Central_Europe_4n | 4x_Europe | H1001 | H1001 | H2001 | H3001 | ERR3514873 | SAMEA5947858 |
| ERR3514874 | 4 | Central_Europe_4n | 4x_Europe | H1001 | H1001 | H2018 | H4017 | ERR3514874 | SAMEA5947859 |
| ERR3514875 | 4 | Central_Europe_4n | 4x_Europe | H1001 | H1001 | H1001 | H1001 | ERR3514875 | SAMEA5947860 |
| ERR3514876 | 4 | Central_Europe_4n | 4x_Europe | H1001 | H1001 | H2009 | H3024 | ERR3514876 | SAMEA5947861 |
| ERR3514877 | 4 | Central_Europe_4n | 4x_Europe | H1001 | H4004 | H4012 | H4041 | ERR3514877 | SAMEA5947862 |
| ERR3514878 | 4 | Central_Europe_4n | 4x_Europe | H1001 | H1001 | H1001 | H4031 | ERR3514878 | SAMEA5947863 |
| ERR3514879 | 4 | Central_Europe_4n | 4x_Europe | H1001 | H1001 | H3003 | H4009 | ERR3514879 | SAMEA5947864 |
| ERR3514880 | 4 | Central_Europe_4n | 4x_Europe | H1001 | H1001 | H1001 | H1001 | ERR3514880 | SAMEA5947865 |
| ERR3514883 | 2 | Central_Europe_2n | 2x_Europe | H1001 | H3008 | | | ERR3514883 | SAMEA5947868 |
| ERR3514884 | 2 | Central_Europe_2n | 2x_Europe | H2004 | H4006 | | | ERR3514884 | SAMEA5947869 |
| ERR3514885 | 2 | Central_Europe_2n | 2x_Europe | H1001 | H4029 | | | ERR3514885 | SAMEA5947870 |
| ERR3514886 | 2 | Central_Europe_2n | 2x_Europe | H1001 | H4010 | | | ERR3514886 | SAMEA5947871 |
| ERR3514887 | 4 | Central_Europe_4n | 4x_Europe | H1001 | H1001 | H4004 | H4034 | ERR3514887 | SAMEA5947872 |
| ERR3514888 | 4 | Central_Europe_4n | 4x_Europe | H1001 | H1001 | H1001 | H2001 | ERR3514888 | SAMEA5947873 |
| ERR3514889 | 4 | Central_Europe_4n | 4x_Europe | H1001 | H1001 | H1001 | H2007 | ERR3514889 | SAMEA5947874 |
| ERR3514892 | 4 | Central_Europe_4n | 4x_Europe | H1001 | H1001 | H1001 | H4010 | ERR3514892 | SAMEA5947877 |
| ERR3514893 | 4 | Central_Europe_4n | 4x_Europe | H1001 | H1001 | H4026 | H4037 | ERR3514893 | SAMEA5947878 |
| ERR3514895 | 4 | Central_Europe_4n | 4x_Europe | H1001 | H3004 | H3010 | H4041 | ERR3514895 | SAMEA5947880 |
| ERR3514896 | 4 | Central_Europe_4n | 4x_Europe | H2004 | H3010 | H4020 | H4021 | ERR3514896 | SAMEA5947881 |
| ERR3514897 | 2 | Central_Europe_2n | 2x_Europe | H3001 | H3001 | | | ERR3514897 | SAMEA5947882 |
| ERR3514898 | 2 | Central_Europe_2n | 2x_Europe | H4011 | H4035 | | | ERR3514898 | SAMEA5947883 |
| ERR3514899 | 2 | Central_Europe_2n | 2x_Europe | H3001 | H4016 | | | ERR3514899 | SAMEA5947884 |
| ERR4235088 | 2 | Central_Europe_2n | 2x_Europe | H1001 | H2007 | | | ERR4235088 | SAMEA6943357 |
| ERR4235089 | 2 | Central_Europe_2n | 2x_Europe | H1001 | H2007 | | | ERR4235089 | SAMEA6943358 |
| ERR4235090 | 2 | Central_Europe_2n | 2x_Europe | H1001 | H4001 | | | ERR4235090 | SAMEA6943359 |
| ERR4235091 | 2 | Central_Europe_2n | 2x_Europe | H4002 | H4008 | | | ERR4235091 | SAMEA6943360 |
| ERR4235092 | 2 | Central_Europe_2n | 2x_Europe | H2007 | H2007 | | | ERR4235092 | SAMEA6943361 |
| ERR4235093 | 2 | Central_Europe_2n | 2x_Europe | H4001 | H4039 | | | ERR4235093 | SAMEA6943362 |
| ERR4235094 | 2 | Central_Europe_2n | 2x_Europe | H4019 | H4040 | | | ERR4235094 | SAMEA6943363 |
| ERR4235095 | 2 | Central_Europe_2n | 2x_Europe | H2007 | H4041 | | | ERR4235095 | SAMEA6943364 |
| ERR4235096 | 2 | Central_Europe_2n | 2x_Europe | H1001 | H4019 | | | ERR4235096 | SAMEA6943365 |
| ERR4235097 | 2 | Central_Europe_2n | 2x_Europe | H4001 | H4001 | | | ERR4235097 | SAMEA6943366 |
| IRK-ID2040 | 2 | Central_Siberia_2n | 2x_Siberia | H1001 | H2007 | | | ERR12736052 | ERS18382732 |
| IRK-ID27652 | 2 | Central_Siberia_2n | 2x_Siberia | H1001 | H1001 | | | ERR12736057 | ERS18382733 |
| IRK-ID61141 | 2 | Central_Siberia_2n | 2x_Siberia | H1001 | H4030 | | | ERR12736056 | ERS18382737 |
| IRK-STAN | 2 | Central_Siberia_2n | 2x_Siberia | H1001 | | | | ERR12736051 | ERS18382738 |
| IRKU049793 | 4 | Central_Siberia_4n | 4x_Siberia | H1001 | H1001 | H1001 | H4027 | ERR12736049 | ERS18382740 |
| IRKU049794 | 4 | Central_Siberia_2n | | H1001 | H1001 | H1001 | H1001 | | |

| | | | | | | | | | |
|---|---|---|---|---|---|---|---|---|---|
| IRKU049798 | 2 | Central_Siberia_2n | 2x_Siberia | H1001 | H4034 | | | ERR12736062 | ERS18382742 |
| IRKU049804 | 2 | Central_Siberia_2n | 2x_Siberia | H2004 | H4027 | | | ERR12736059 | ERS18382745 |
| IRKU084814 | 2 | Central_Siberia_2n | 2x_Siberia | H1001 | H4038 | | | ERR12736058 | ERS18382746 |
| MN_47 | 2 | North_America_2n | | H3008 | H3008 | | | ERR12742582 | ERS18382747 |
| MW0079456 | 2 | selfer | | H4004 | H4004 | | | ERR9802385 | ERS10113953 |
| MW0079467_1 | 2 | East_Siberia_2n | 2x_Siberia | H3004 | H3011 | | | ERR12742392 | ERS18382748 |
| MW0079473_1 | 2 | East_Siberia_2n | 2x_Siberia | H1001 | H4039 | | | ERR12742391 | ERS18382750 |
| MW0079478 | 2 | Amur_Basin_2n | | H4009 | H4013 | | | ERR12735972 | ERS18382751 |
| MW0079478_3 | 2 | Amur_Basin_2n | | H4009 | H4013 | | | ERR12742390 | ERS18382752 |
| MW0079479 | 2 | Amur_Basin_2n | | H4016 | H4037 | | | ERR12735969 | ERS18382753 |
| MW0079480 | 2 | Amur_Basin_2n | | H4028 | H4029 | | | ERR12735968 | ERS18382754 |
| MW0079480_3 | 2 | Amur_Basin_2n | | H4028 | H4029 | | | ERR12742389 | ERS18382755 |
| MW0079481 | 2 | Amur_Basin_2n | | H1001 | H1001 | | | ERR12735991 | ERS18382756 |
| MW0079481_2 | 2 | Amur_Basin_2n | | H1001 | H1001 | | | ERR12742388 | ERS18382757 |
| MW0079482 | 2 | Amur_Basin_2n | | H2007 | H4037 | | | ERR12735971 | ERS18382758 |
| MW0079482_2 | 2 | Amur_Basin_2n | | H2007 | H4037 | | | ERR12742387 | ERS18382759 |
| MW0079483 | 2 | Amur_Basin_2n | | H1001 | H4036 | | | ERR12735970 | ERS18382760 |
| MW0079484_2 | 2 | Amur_Basin_2n | | H1001 | H2007 | | | ERR12742386 | ERS18382761 |
| MW0079485 | 2 | Amur_Basin_2n | | H1001 | H4024 | | | ERR12736003 | ERS18382762 |
| MW0079485_2 | 2 | Amur_Basin_2n | | H1001 | H4024 | | | ERR12742385 | ERS18382763 |
| MW0079486_3 | 2 | East_Siberia_2n | 2x_Siberia | H2009 | H2009 | | | ERR12742383 | ERS18382765 |
| MW0079491 | 2 | East_Siberia_2n | 2x_Siberia | H3006 | H4030 | | | ERR12735992 | ERS18382766 |
| MW0079491_2 | 2 | East_Siberia_2n | 2x_Siberia | H1001 | H4005 | | | ERR12742382 | ERS18382767 |
| MW0079493_1 | 2 | East_Siberia_2n | 2x_Siberia | H2007 | H2009 | | | ERR12742381 | ERS18382768 |
| MW0079494_2 | 2 | East_Siberia_2n | 2x_Siberia | H4010 | H4009 | | | ERR12742380 | ERS18382769 |
| MW0079496 | 2 | East_Siberia_2n | 2x_Siberia | H4006 | H4037 | | | ERR12735965 | ERS18382770 |
| MW0079497 | 2 | KIS_2n | | H2004 | H4026 | | | ERR12735996 | ERS18382771 |
| MW0079497_4 | 2 | KIS_2n | | H2004 | H4026 | | | ERR12742379 | ERS18382772 |
| MW0079500 | 2 | KIS_2n | | H4005 | H4007 | | | ERR12735975 | ERS18382773 |
| MW0079500_2 | 2 | KIS_2n | | H2004 | H4010 | | | ERR12742378 | ERS18382774 |
| MW0079507 | 4 | Central_Siberia_4n | 4x_Siberia | H2006 | H3002 | H4002 | H4023 | ERR12736002 | ERS18382775 |
| MW0079509 | 2 | Northern_Ural_2n | | H2003 | H2003 | | | ERR12735987 | ERS18382776 |
| MW0079510_1 | 2 | Northern_Ural_2n | | H1001 | H4010 | | | ERR12742377 | ERS18382777 |
| MW0079537 | 2 | East_Siberia_2n | 2x_Siberia | H4015 | H4039 | | | ERR12736015 | ERS18382778 |
| MW0079537_3 | 2 | East_Siberia_2n | 2x_Siberia | H2001 | H2009 | | | ERR12742376 | ERS18382779 |
| MW0079538 | 2 | East_Siberia_2n | 2x_Siberia | H4019 | H4037 | | | ERR12736016 | ERS18382780 |
| MW0079543_2 | 2 | East_Siberia_2n | 2x_Siberia | H3006 | H4026 | | | ERR12742375 | ERS18382781 |
| MW0079544 | 4 | Central_Siberia_4n | 4x_Siberia | H2003 | H2009 | H4004 | H4029 | ERR12736017 | ERS18382782 |
| MW0079545 | 2 | East_Siberia_2n | 2x_Siberia | H4010 | H4014 | | | ERR12736018 | ERS18382783 |
| MW0079546 | 4 | Central_Siberia_4n | 4x_Siberia | H2003 | H2006 | H3002 | H4002 | ERR12736019 | ERS18382784 |
| MW0079546_2 | 4 | Central_Siberia_4n | 4x_Siberia | H2003 | H2003 | H2006 | H4002 | ERR12742374 | ERS18382785 |
| MW0079552 | 4 | Central_Siberia_4n | 4x_Siberia | H1001 | H2004 | H3014 | H4024 | ERR12735998 | ERS18382786 |
| MW0079552_2 | 4 | Central_Siberia_4n | 4x_Siberia | H1001 | H3012 | H4034 | | ERR12742373 | ERS18382787 |
| MW0079552_3 | 4 | Central_Siberia_4n | 4x_Siberia | H1001 | H2006 | H4010 | H4019 | ERR12742372 | ERS18382788 |
| MW0079552_7 | 4 | Central_Siberia_4n | 4x_Siberia | H1001 | H1001 | H3022 | H4023 | ERR12742371 | ERS18382789 |
| MW0079552_8 | 4 | Central_Siberia_4n | 4x_Siberia | H1001 | H3022 | H4023 | H4039 | ERR12742370 | ERS18382790 |
| MW0079553 | 2 | KIS_2n | | H4012 | H4015 | | | ERR12735976 | ERS18382791 |
| MW0079553_5 | 2 | KIS_2n | | H1001 | H4042 | | | ERR12742369 | ERS18382792 |
| MW0079553_7 | 2 | KIS_2n | | H1001 | H4042 | | | ERR12742368 | ERS18382793 |
| MW0079558_2 | 4 | Central_Siberia_4n | 4x_Siberia | H3014 | H4001 | H4005 | H4029 | ERR12742367 | ERS18382794 |
| MW0079559 | 2 | West_Siberia_2n | | H1001 | H3008 | | | ERR12735986 | ERS18382795 |
| MW0079560 | 2 | Northern_Ural_2n | | H3011 | H4005 | | | ERR12742329 | ERS18382796 |
| MW0079561_8 | 2 | Northern_Ural_2n | | H3002 | H4018 | | | ERR12742366 | ERS18382797 |
| MW0079561_9 | 2 | Northern_Ural_2n | | H1001 | H4041 | | | ERR12742365 | ERS18382798 |
| MW0079564 | 2 | Northern_Ural_2n | | H1001 | H1001 | | | ERR12735988 | ERS18382799 |
| MW0079564_2 | 2 | Northern_Ural_2n | | H1001 | H1001 | | | ERR12742364 | ERS18382800 |
| MW0079568-1_1 | 4 | Central_Siberia_4n | 4x_Siberia | H1001 | H1001 | H3009 | H4029 | ERR12742363 | ERS18382801 |
| MW0079568-3_1 | 4 | Central_Siberia_4n | 4x_Siberia | H1001 | H1001 | H1001 | H4004 | ERR12742361 | ERS18382803 |
| MW0079568-4_1 | 4 | Central_Siberia_4n | 4x_Siberia | H1001 | H1001 | H2003 | H4028 | ERR12742360 | ERS18382804 |
| MW0079569-2_1 | 4 | Central_Siberia_4n | 4x_Siberia | H1001 | H1001 | H4021 | H4023 | ERR12742359 | ERS18382805 |
| MW0079572-1 | 4 | Central_Siberia_4n | 4x_Siberia | H1001 | H3014 | H4010 | H4029 | ERR12735978 | ERS18382806 |
| MW0079572-2 | 4 | Central_Siberia_4n | 4x_Siberia | H1001 | H2009 | H4035 | H4041 | ERR12735999 | ERS18382807 |
| MW0079572-3_1 | 4 | Central_Siberia_4n | 4x_Siberia | H1001 | H1001 | H2004 | H4026 | ERR12742358 | ERS18382808 |
| MW0079573 | 4 | Central_Siberia_4n | 4x_Siberia | H1001 | H4001 | H4026 | H4028 | ERR12735977 | ERS18382809 |
| MW0079573_1 | 4 | Central_Siberia_4n | 4x_Siberia | H2008 | H4002 | H4006 | H3012 | ERR12742357 | ERS18382810 |
| MW0079573_3 | 4 | Central_Siberia_4n | 4x_Siberia | H1001 | H2008 | H4019 | H4035 | ERR12742356 | ERS18382811 |
| MW0079573_4 | 4 | Central_Siberia_4n | 4x_Siberia | H1001 | H2008 | H4019 | H4035 | ERR12742355 | ERS18382812 |
| MW0079573_5 | 4 | Central_Siberia_4n | 4x_Siberia | H1001 | H2008 | H4019 | H4035 | ERR12742354 | ERS18382813 |
| MW0079575 | 4 | Central_Siberia_4n | 4x_Siberia | H1001 | H2004 | H4020 | H4037 | ERR12736020 | ERS18382814 |
| MW0079577_2 | 4 | Central_Siberia_4n | 4x_Siberia | H1001 | H1001 | H1001 | H4035 | ERR12742353 | ERS18382815 |

| | | | | | | | | | |
|---|---|---|---|---|---|---|---|---|---|
| MW0079578-1 | 4 | Central_Siberia_4n | 4x_Siberia | H2007 | H3002 | H3002 | H4015 | ERR12735997 | ERS18382816 |
| MW0079579 | 2 | Central_Siberia_2n | 2x_Siberia | H2001 | H4039 | | | ERR9802377 | ERS10113962 |
| MW0079581 | 2 | Central_Siberia_2n | 2x_Siberia | H3006 | H4003 | | | ERR9802376 | ERS10113963 |
| MW0079582 | 2 | Northern_Ural_2n | 2x_Siberia | H1001 | H4010 | | | ERR12735985 | ERS18382817 |
| MW0079583_1 | 2 | Northern_Ural_2n | 2x_Siberia | H2003 | H2003 | | | ERR12742352 | ERS18382818 |
| MW0079583_2 | 2 | Northern_Ural_2n | 2x_Siberia | H2003 | H2003 | | | ERR12742351 | ERS18382819 |
| MW0079583_3 | 2 | Northern_Ural_2n | 2x_Siberia | H2003 | H2003 | | | ERR12742350 | ERS18382820 |
| MW0079584 | 4 | Northern_Ural_4n | 4x_Siberia | H1001 | H2001 | H2004 | H4012 | ERR12735984 | ERS18382821 |
| MW0079585 | 2 | Northern_Ural_2n | 2x_Siberia | H1001 | H4001 | | | ERR12735967 | ERS18382822 |
| MW0079587_4 | 2 | West_Siberia_2n | | H4012 | H4035 | | | ERR12742349 | ERS18382823 |
| MW0079587_6 | 2 | West_Siberia_2n | | H4012 | H4035 | | | ERR12742348 | ERS18382824 |
| MW0079589 | 4 | Northern_Ural_4n | 4x_Siberia | H1001 | H1001 | H4020 | H4038 | ERR12735990 | ERS18382825 |
| MW0079589_8 | 4 | Northern_Ural_4n | 4x_Siberia | H1001 | H3024 | H4010 | H4013 | ERR12742346 | ERS18382827 |
| MW0079589_9 | 4 | Northern_Ural_4n | 4x_Siberia | H1001 | H2001 | H2008 | H4007 | ERR12742345 | ERS18382828 |
| MW0079591 | 2 | Northern_Ural_2n | 2x_Siberia | H2007 | H4012 | | | ERR12735994 | ERS18382829 |
| MW0079604 | 2 | Central_Siberia_2n | 2x_Siberia | H1001 | H1001 | | | ERR12735974 | ERS18382830 |
| MW0079604_5 | 2 | Central_Siberia_2n | 2x_Siberia | H1001 | H1001 | | | ERR12742344 | ERS18382831 |
| MW0079606 | 4 | Central_Siberia_4n | 4x_Siberia | H1001 | H2001 | H3008 | H4010 | ERR12735995 | ERS18382832 |
| MW0079608_3 | 2 | West_Siberia_2n | | H1001 | H4005 | | | ERR12742342 | ERS18382834 |
| MW0079609 | 4 | Central_Siberia_4n | 4x_Siberia | H1001 | H1001 | H1001 | H3010 | ERR12735993 | ERS18382835 |
| MW0079609_3 | 4 | Central_Siberia_4n | 4x_Siberia | H1001 | H1001 | H4024 | H4024 | ERR12742341 | ERS18382836 |
| MW0079609_4 | 4 | Central_Siberia_4n | 4x_Siberia | H1001 | H1001 | H2008 | H3006 | ERR12742340 | ERS18382837 |
| MW0079610 | 4 | Central_Siberia_4n | 4x_Siberia | H1001 | H3011 | H4039 | H4034 | ERR12736000 | ERS18382838 |
| MW0079895 | 2 | Northern_Ural_2n | 2x_Siberia | H1001 | H4005 | | | ERR12735989 | ERS18382839 |
| MW0157476_2 | 2 | Central_Siberia_2n | 2x_Siberia | H1001 | H4035 | | | ERR12742339 | ERS18382840 |
| MW0157478 | 4 | Central_Siberia_2n | 4x_Siberia | H1001 | H2001 | H4038 | H4042 | ERR12736004 | ERS18382841 |
| MW0157478_2 | 4 | Central_Siberia_2n | 4x_Siberia | H1001 | H1001 | H1001 | H3011 | ERR12742338 | ERS18382842 |
| MW0158706 | 2 | Central_Siberia_2n | 2x_Siberia | H4017 | H4018 | | | ERR9802390 | ERS10113958 |
| MW0158707 | 2 | Central_Siberia_2n | 2x_Siberia | H4012 | H4013 | | | ERR9802389 | ERS10113957 |
| MW0158708 | 4 | East_Siberia_4n | 4x_Siberia | H3003 | H3003 | H2009 | H2009 | ERR12735980 | ERS18382843 |
| MW0158708_3 | 4 | East_Siberia_4n | | H3003 | H3003 | H2009 | H2009 | | |
| MW0158708_4 | 4 | East_Siberia_4n | | H3003 | H3003 | H2009 | H2009 | | |
| MW0158708_6 | 4 | East_Siberia_4n | | H3003 | H3003 | H2009 | H2009 | | |
| MW0374083 | 4 | Northern_Ural_4n | 4x_Siberia | H1001 | H1001 | H2004 | H4035 | ERR12735979 | ERS18382844 |
| MW0374083_4 | 4 | Northern_Ural_4n | 4x_Siberia | H1001 | H1001 | H2004 | H4013 | ERR12742337 | ERS18382845 |
| MW0374083_6 | 4 | Northern_Ural_4n | 4x_Siberia | H1001 | H1001 | H2004 | H4035 | ERR12742336 | ERS18382846 |
| MW0374084 | 4 | Northern_Ural_4n | 4x_Siberia | H1001 | H1001 | H1001 | H1001 | ERR12735983 | ERS18382847 |
| MW0374084_2 | 4 | Northern_Ural_4n | 4x_Siberia | H2028 | H3002 | H4015 | H4039 | ERR12742335 | ERS18382848 |
| MW0374085_1 | 4 | Northern_Ural_4n | 4x_Siberia | H1001 | H2007 | H3015 | H4039 | ERR12742334 | ERS18382849 |
| MW0374085_11 | 4 | Northern_Ural_4n | 4x_Siberia | H4010 | H4010 | H3022 | H4020 | ERR12742333 | ERS18382850 |
| MW0374087 | 4 | Northern_Ural_4n | 4x_Siberia | H2004 | H3011 | H4015 | H4021 | ERR12736001 | ERS18382851 |
| MW0374087_5 | 4 | Northern_Ural_4n | 4x_Siberia | H1001 | H2007 | H3010 | H4015 | ERR12742332 | ERS18382852 |
| MW0374087_9 | 4 | Northern_Ural_4n | | H2007 | H2007 | H4015 | - | ERR12742331 | ERS18382853 |
| MW0374094_6 | 4 | | | H2004 | H2004 | - | - | ERR12742330 | ERS18382854 |
| MW0374095 | 4 | Northern_Ural_4n | 4x_Siberia | H1001 | H1001 | H1001 | H1001 | ERR12735982 | ERS18382855 |
| MW0374101 | 4 | Northern_Ural_4n | 4x_Siberia | H1001 | H4014 | H4022 | H4034 | ERR12735981 | ERS18382856 |
| MW0374102-2 | 4 | Northern_Ural_4n | 4x_Siberia | H1001 | H1001 | H1001 | H3011 | ERR12735973 | ERS18382857 |
| NT10_1_1 | 2 | East_Siberia_2n | 2x_Siberia | H4030 | H4040 | | | ERR12736022 | ERS18382858 |
| NT10_2_1 | 2 | East_Siberia_2n | 2x_Siberia | H2003 | H4019 | | | ERR12736047 | ERS18382859 |
| NT10_3_1 | 2 | East_Siberia_2n | 2x_Siberia | H2003 | H3004 | | | ERR12736013 | ERS18382860 |
| NT12_1_1 | 2 | East_Siberia_2n | 2x_Siberia | H4001 | H4004 | | | ERR9802379 | ERS10113960 |
| NT12_2_1 | 2 | East_Siberia_2n | 2x_Siberia | H4004 | H4040 | | | ERR9802378 | ERS10113961 |
| NT13_9_1 | 2 | East_Siberia_2n | 2x_Siberia | H1001 | H4038 | | | ERR12736012 | ERS18382861 |
| NT14_1_1 | 2 | East_Siberia_2n | 2x_Siberia | H1001 | H4010 | | | ERR12736011 | ERS18382862 |
| NT14_10_1 | 2 | East_Siberia_2n | 2x_Siberia | H3011 | H3011 | | | ERR12736044 | ERS18382863 |
| NT14_11_1 | 2 | East_Siberia_2n | 2x_Siberia | H4029 | H4043 | | | ERR12736040 | ERS18382864 |
| NT14_3_1 | 2 | East_Siberia_2n | 2x_Siberia | H1001 | H4013 | | | ERR12736010 | ERS18382865 |
| NT14_4_1 | 2 | East_Siberia_2n | 2x_Siberia | H3011 | H3011 | | | ERR12736009 | ERS18382866 |
| NT14_5_1 | 2 | East_Siberia_2n | 2x_Siberia | H2006 | H4030 | | | ERR12736028 | ERS18382867 |
| NT14_6_1 | 2 | East_Siberia_2n | 2x_Siberia | H4026 | H4027 | | | ERR12736025 | ERS18382868 |
| NT14_7_1 | 2 | East_Siberia_2n | 2x_Siberia | H3012 | H4014 | | | ERR12736021 | ERS18382869 |
| NT14_8_1 | 2 | East_Siberia_2n | 2x_Siberia | H1001 | H1001 | | | ERR12736046 | ERS18382870 |
| NT15_1_1 | 2 | East_Siberia_2n | 2x_Siberia | H3011 | H4010 | | | ERR12736035 | ERS18382871 |
| NT15_2_1 | 2 | East_Siberia_2n | 2x_Siberia | H2003 | H4028 | | | ERR12736008 | ERS18382872 |
| NT15_3_1 | 2 | East_Siberia_2n | 2x_Siberia | H3014 | H4043 | | | ERR12736027 | ERS18382873 |
| NT15_4_1 | 2 | East_Siberia_2n | 2x_Siberia | H4019 | H4020 | | | ERR12736024 | ERS18382874 |
| NT15_5_1 | 2 | East_Siberia_2n | 2x_Siberia | H4002 | H4021 | | | ERR12736007 | ERS18382875 |
| NT16.1-1 | 2 | East_Siberia_2n | 2x_Siberia | H2004 | H4012 | | | ERR12736119 | ERS18382876 |
| NT16.1-2 | 2 | East_Siberia_2n | 2x_Siberia | H2004 | H4041 | | | ERR12736118 | ERS18382877 |
| NT16.1-4 | 2 | East_Siberia_2n | 2x_Siberia | H1001 | H1001 | | | ERR12736117 | ERS18382878 |

| | | | | | | | | | |
|---|---|---|---|---|---|---|---|---|---|
| NT16.1-5 | 2 | East_Siberia_2n | 2x_Siberia | H1001 | H4041 | | | ERR12736116 | ERS18382879 |
| NT16.2-1 | 2 | East_Siberia_2n | 2x_Siberia | H1001 | H4038 | | | ERR12736115 | ERS18382880 |
| NT16.2-2 | 2 | East_Siberia_2n | 2x_Siberia | H1001 | H4038 | | | ERR12736114 | ERS18382881 |
| NT16.3-1 | 2 | East_Siberia_2n | 2x_Siberia | H4004 | H4034 | | | ERR12736113 | ERS18382882 |
| NT16.3-5 | 2 | East_Siberia_2n | 2x_Siberia | H1001 | H4002 | | | ERR12736112 | ERS18382883 |
| NT16.3-6 | 2 | East_Siberia_2n | 2x_Siberia | H1001 | H4034 | | | ERR12736111 | ERS18382884 |
| NT16.3-7 | 2 | East_Siberia_2n | 2x_Siberia | H4034 | H4041 | | | ERR12736110 | ERS18382885 |
| NT16.4-1 | 2 | East_Siberia_2n | 2x_Siberia | H2004 | H4009 | | | ERR12736109 | ERS18382886 |
| NT16.4-2 | 2 | East_Siberia_2n | 2x_Siberia | H1001 | H4041 | | | ERR12736108 | ERS18382887 |
| NT16.4-3 | 2 | East_Siberia_2n | 2x_Siberia | H4020 | H4041 | | | ERR12736107 | ERS18382888 |
| NT16.4-4 | 2 | East_Siberia_2n | 2x_Siberia | H1001 | H4034 | | | ERR12736106 | ERS18382889 |
| NT16.5-1 | 2 | East_Siberia_2n | 2x_Siberia | H2003 | H4002 | | | ERR12736105 | ERS18382890 |
| NT16.5-2 | 2 | East_Siberia_2n | 2x_Siberia | H1001 | H4025 | | | ERR12736104 | ERS18382891 |
| NT16.5-3 | 2 | East_Siberia_2n | 2x_Siberia | H1001 | H3011 | | | ERR12736103 | ERS18382892 |
| NT16.5-4 | 2 | East_Siberia_2n | 2x_Siberia | H2003 | H2004 | | | ERR12736102 | ERS18382893 |
| NT17.1-1 | 2 | East_Siberia_2n | 2x_Siberia | H3010 | H3012 | | | ERR12736101 | ERS18382894 |
| NT17.1-2 | 2 | East_Siberia_2n | 2x_Siberia | H2004 | H3012 | | | ERR12736100 | ERS18382895 |
| NT17.1-4 | 2 | East_Siberia_2n | 2x_Siberia | H3012 | H4019 | | | ERR12736099 | ERS18382896 |
| NT17.1-5 | 2 | East_Siberia_2n | 2x_Siberia | H4019 | H4027 | | | ERR12736098 | ERS18382897 |
| NT17.4-1 | 2 | East_Siberia_2n | 2x_Siberia | H3002 | H4021 | | | ERR12736097 | ERS18382898 |
| NT17.4-2 | 2 | East_Siberia_2n | 2x_Siberia | H3002 | H3008 | | | ERR12736096 | ERS18382899 |
| NT17.4-3 | 2 | East_Siberia_2n | 2x_Siberia | H4021 | H4041 | | | ERR12736095 | ERS18382900 |
| NT17.4-4 | 2 | East_Siberia_2n | 2x_Siberia | H1001 | H4038 | | | ERR12736094 | ERS18382901 |
| NT17.5-1 | 2 | East_Siberia_2n | 2x_Siberia | H1001 | H4017 | | | ERR12736093 | ERS18382902 |
| NT18-1 | 2 | East_Siberia_2n | 2x_Siberia | H2006 | H3004 | | | ERR12736092 | ERS18382903 |
| NT18-2 | 2 | East_Siberia_2n | 2x_Siberia | H2003 | H2003 | | | ERR12736091 | ERS18382904 |
| NT18-3 | 2 | East_Siberia_2n | 2x_Siberia | H2006 | H2006 | | | ERR12736090 | ERS18382905 |
| NT18-4 | 2 | East_Siberia_2n | 2x_Siberia | H2006 | H3010 | | | ERR12736089 | ERS18382906 |
| NT18-6 | 2 | East_Siberia_2n | 2x_Siberia | H2006 | H4004 | | | ERR12736088 | ERS18382907 |
| NT19.1-1 | 2 | East_Siberia_2n | 2x_Siberia | H4027 | H4041 | | | ERR12736087 | ERS18382908 |
| NT19.1-2 | 2 | East_Siberia_2n | 2x_Siberia | H2004 | H3002 | | | ERR12736086 | ERS18382909 |
| NT19.1-3 | 2 | East_Siberia_2n | 2x_Siberia | H4008 | H4027 | | | ERR12736085 | ERS18382910 |
| NT19.1-4 | 2 | East_Siberia_2n | 2x_Siberia | H3001 | H3002 | | | ERR12736084 | ERS18382911 |
| NT19.1-5 | 2 | East_Siberia_2n | 2x_Siberia | H2001 | H3002 | | | ERR12736083 | ERS18382912 |
| NT19.1-6 | 2 | East_Siberia_2n | 2x_Siberia | H4014 | H4027 | | | ERR12736082 | ERS18382913 |
| NT19.2-1 | 2 | East_Siberia_2n | 2x_Siberia | H2003 | H4035 | | | ERR12736081 | ERS18382914 |
| NT19.2-2 | 2 | East_Siberia_2n | 2x_Siberia | H4019 | H4041 | | | ERR12736080 | ERS18382915 |
| NT19.3-1 | 2 | East_Siberia_2n | 2x_Siberia | H4025 | H4036 | | | ERR12736079 | ERS18382916 |
| NT19.3-2 | 2 | East_Siberia_2n | 2x_Siberia | H4025 | H4036 | | | ERR12736078 | ERS18382917 |
| NT19.3-3 | 2 | East_Siberia_2n | 2x_Siberia | H4025 | H4027 | | | ERR12736077 | ERS18382918 |
| NT2_1_1 | 2 | East_Siberia_2n | 2x_Siberia | H4016 | H4018 | | | ERR12736038 | ERS18382919 |
| NT2_2_1 | 2 | East_Siberia_2n | 2x_Siberia | H3009 | H3009 | | | ERR12736033 | ERS18382920 |
| NT2_3_1 | 2 | East_Siberia_2n | 2x_Siberia | H1001 | H4042 | | | ERR12736030 | ERS18382921 |
| NT2_4_1 | 2 | East_Siberia_2n | 2x_Siberia | H3009 | H4041 | | | ERR12736006 | ERS18382922 |
| NT2_5_1 | 2 | East_Siberia_2n | 2x_Siberia | H2009 | H4042 | | | ERR12736023 | ERS18382923 |
| NT20.1-1 | 2 | East_Siberia_2n | 2x_Siberia | H3002 | H4043 | | | ERR12736076 | ERS18382924 |
| NT20.3-1 | 2 | East_Siberia_2n | 2x_Siberia | H4034 | H4043 | | | ERR12736075 | ERS18382925 |
| NT20.3-2 | 2 | East_Siberia_2n | 2x_Siberia | H4012 | H4034 | | | ERR12736074 | ERS18382926 |
| NT20.3-3 | 2 | East_Siberia_2n | 2x_Siberia | H4034 | H4043 | | | ERR12736073 | ERS18382927 |
| NT20.4-1 | 2 | East_Siberia_2n | 2x_Siberia | H4036 | H4040 | | | ERR12736072 | ERS18382928 |
| NT20.4-2 | 2 | East_Siberia_2n | 2x_Siberia | H4036 | H4040 | | | ERR12736071 | ERS18382929 |
| NT20.4-3 | 2 | East_Siberia_2n | 2x_Siberia | H4043 | H4043 | | | ERR12736070 | ERS18382930 |
| NT20.6-2 | 2 | East_Siberia_2n | 2x_Siberia | H1001 | H2004 | | | ERR12736069 | ERS18382931 |
| NT20.6-3 | 2 | East_Siberia_2n | 2x_Siberia | H4034 | H4043 | | | ERR12736068 | ERS18382932 |
| NT4_2_1 | 2 | East_Siberia_2n | 2x_Siberia | H1001 | H4018 | | | ERR12736048 | ERS18382933 |
| NT4_3_1 | 2 | East_Siberia_2n | 2x_Siberia | H3004 | H4039 | | | ERR12736045 | ERS18382934 |
| NT4.1 | 2 | East_Siberia_2n | 2x_Siberia | H3009 | H4028 | | | ERR12736067 | ERS18382935 |
| NT5_1_1 | 2 | East_Siberia_2n | 2x_Siberia | H1001 | H4042 | | | ERR12736042 | ERS18382936 |
| NT5_2_1 | 2 | East_Siberia_2n | 2x_Siberia | H3006 | H3022 | | | ERR12736037 | ERS18382937 |
| NT5_3_1 | 2 | East_Siberia_2n | 2x_Siberia | H3006 | H3022 | | | ERR12736032 | ERS18382938 |
| NT8_1_1 | 2 | East_Siberia_2n | 2x_Siberia | H4023 | H4041 | | | ERR9802386 | ERS10113954 |
| NT8_2_1 | 2 | East_Siberia_2n | 2x_Siberia | H1001 | H4019 | | | ERR9802387 | ERS10113955 |
| NT8_3_1 | 2 | East_Siberia_2n | 2x_Siberia | H3003 | H4001 | | | ERR9802388 | ERS10113956 |
| NT8_4_1 | 2 | East_Siberia_2n | 2x_Siberia | H3011 | H4004 | | | ERR9802380 | ERS10113959 |
| NT8_5_1 | 2 | East_Siberia_2n | 2x_Siberia | H4008 | H4034 | | | ERR12736005 | ERS18382939 |
| NT8.3-4 | 2 | East_Siberia_2n | 2x_Siberia | H4034 | H4037 | | | ERR12736066 | ERS18382940 |
| NT8.4-5 | 2 | East_Siberia_2n | 2x_Siberia | H3011 | H4038 | | | ERR12736065 | ERS18382941 |
| NT9_1_1 | 2 | East_Siberia_2n | 2x_Siberia | H1001 | H3003 | | | ERR12736041 | ERS18382942 |
| NT9_2_1 | 2 | East_Siberia_2n | 2x_Siberia | H1001 | H3003 | | | ERR12736036 | ERS18382943 |
| NT9_3_1 | 2 | East_Siberia_2n | 2x_Siberia | H4020 | H4035 | | | ERR12736031 | ERS18382944 |

| | | | | | | | | | |
|---|---|---|---|---|---|---|---|---|---|
| NT9_4_1 | 2 | East_Siberia_2n | 2x_Siberia | H4008 | H4016 | | | ERR12736029 | ERS18382945 |
| NT9_5_1 | 2 | East_Siberia_2n | 2x_Siberia | H1001 | H3008 | | | ERR12736026 | ERS18382946 |
| PU_1mix-1 | 4 | Northern_Ural_4n | 4x_Siberia | H1001 | H1001 | H3022 | H4012 | ERR12742581 | ERS18382947 |
| PU_1mix-2 | 4 | Northern_Ural_4n | 4x_Siberia | H1001 | H1001 | H1001 | H2001 | ERR12742580 | ERS18382948 |
| PU_2mix-1 | 4 | Northern_Ural_4n | 4x_Siberia | H1001 | H1001 | H1001 | H2008 | ERR12742579 | ERS18382949 |
| PU_2mix-2 | 4 | Northern_Ural_4n | 4x_Siberia | H1001 | H1001 | H2004 | H2008 | ERR12742578 | ERS18382950 |
| PU_3mix-1 | 4 | Northern_Ural_4n | 4x_Siberia | H1001 | H1001 | H1001 | H4027 | ERR12742577 | ERS18382951 |
| PU_3mix-2 | 4 | Northern_Ural_4n | 4x_Siberia | H1001 | H2004 | H3001 | - | ERR12742576 | ERS18382952 |
| PU_4mix-1 | 4 | Northern_Ural_4n | 4x_Siberia | H1001 | H2003 | H3008 | H4003 | ERR12742575 | ERS18382953 |
| PU_4mix-2 | 4 | Northern_Ural_4n | 4x_Siberia | H1001 | H1001 | H3021 | H4008 | ERR12742574 | ERS18382954 |
| PU_5mix-1 | 4 | Northern_Ural_4n | 4x_Siberia | H1001 | H2001 | H3010 | H4018 | ERR12742573 | ERS18382955 |
| PU_5mix-2 | 4 | Northern_Ural_4n | 4x_Siberia | H1001 | H1001 | H1001 | H4038 | ERR12742572 | ERS18382956 |
| PU_6.1-1 | 4 | Northern_Ural_4n | 4x_Siberia | H1001 | H1001 | H1001 | H4006 | ERR12742571 | ERS18382957 |
| PU_6.1-2 | 4 | Northern_Ural_4n | 4x_Siberia | H1001 | H1001 | H1001 | H2001 | ERR12742570 | ERS18382958 |
| PU_6.2-1 | 4 | Northern_Ural_4n | 4x_Siberia | H1001 | H1001 | H1001 | H2001 | ERR12742569 | ERS18382959 |
| PU_6.2-2 | 4 | Northern_Ural_4n | 4x_Siberia | H1001 | H1001 | H1001 | H4017 | ERR12742568 | ERS18382960 |
| PU_6.3-1 | 4 | Northern_Ural_4n | 4x_Siberia | H1001 | H1001 | H1001 | H4030 | ERR12742567 | ERS18382961 |
| PU_6.3-2 | 4 | Northern_Ural_4n | 4x_Siberia | H1001 | H1001 | H3004 | H4028 | ERR12742566 | ERS18382962 |
| PU_6.4-1 | 4 | Northern_Ural_4n | 4x_Siberia | H1001 | H2006 | H2006 | H4027 | ERR12742565 | ERS18382963 |
| PU_6.4-2 | 4 | Northern_Ural_4n | 4x_Siberia | H1001 | H2003 | H4013 | H4022 | ERR12742564 | ERS18382964 |
| PU_6.5-1 | 4 | Northern_Ural_4n | 4x_Siberia | H1001 | H1001 | H2001 | H3004 | ERR12742563 | ERS18382965 |
| PU_6.5-2_4n | 4 | Northern_Ural_4n | 4x_Siberia | H1001 | H4031 | H4037 | H4038 | ERR12742562 | ERS18382966 |
| PU_6.6-1 | 4 | Northern_Ural_4n | 4x_Siberia | H1001 | H1001 | H2028 | H4003 | ERR12742561 | ERS18382967 |
| PU_6.6-2 | 4 | Northern_Ural_4n | 4x_Siberia | H1001 | H1001 | H4003 | H4034 | ERR12742560 | ERS18382968 |
| PU_6.7-1 | 4 | Northern_Ural_4n | 4x_Siberia | H1001 | H1001 | H1001 | H4010 | ERR12742559 | ERS18382969 |
| PU_6.7-2 | 4 | Northern_Ural_4n | 4x_Siberia | H1001 | H1001 | H2007 | H4018 | ERR12742558 | ERS18382970 |
| PU_6.8-1 | 4 | Northern_Ural_4n | 4x_Siberia | H1001 | H1001 | H2007 | H3011 | ERR12742557 | ERS18382971 |
| PU_6.8-2 | 4 | Northern_Ural_4n | 4x_Siberia | H1001 | H1001 | H2028 | H4022 | ERR12742556 | ERS18382972 |
| PU_6mix-1 | 4 | Northern_Ural_4n | 4x_Siberia | H1001 | H2001 | H2006 | H2008 | ERR12742555 | ERS18382973 |
| PU_6mix-2 | 4 | Northern_Ural_4n | 4x_Siberia | H1001 | H1001 | H2001 | H2009 | ERR12742554 | ERS18382974 |
| PU_7.1-1 | 4 | Northern_Ural_4n | 4x_Siberia | H1001 | H4038 | H4039 | H4041 | ERR12742553 | ERS18382975 |
| PU_7.1-2 | 4 | Northern_Ural_4n | 4x_Siberia | H1001 | H1001 | H1001 | H4024 | ERR12742552 | ERS18382976 |
| PU_7.2-1 | 4 | Northern_Ural_4n | 4x_Siberia | H1001 | H1001 | H1001 | H4024 | ERR12742551 | ERS18382977 |
| PU_7.2-2 | 4 | Northern_Ural_4n | 4x_Siberia | H1001 | H1001 | H1001 | H1001 | ERR12742550 | ERS18382978 |
| PU_7mix-1 | 4 | Northern_Ural_4n | 4x_Siberia | H1001 | H1001 | H2003 | H4041 | ERR12742549 | ERS18382979 |
| PU_7mix-2 | 4 | Northern_Ural_4n | 4x_Siberia | H1001 | H2001 | H2003 | H4041 | ERR12742548 | ERS18382980 |
| SRR2040769 | 2 | North_America_2n | | H4018 | H4018 | | | SRR2040769 | SAMN03703442 |
| SRR2040770 | 2 | North_America_2n | | H4018 | H4018 | | | SRR2040770 | SAMN03703443 |
| SRR2040788 | 2 | North_America_2n | | H1001 | H1001 | | | SRR2040788 | SAMN03703461 |
| SRR2040789 | 2 | North_America_2n | | H1001 | H4042 | | | SRR2040789 | SAMN03703462 |
| SRR2040790 | 2 | Central_Europe_2n | 2x_Europe | H1001 | H4040 | | | SRR2040790 | SAMN03703463 |
| SRR2040791 | 2 | Central_Europe_2n | 2x_Europe | H1001 | H4004 | | | SRR2040791 | SAMN03703464 |
| SRR2040793 | 2 | Scandinavia_UK_2n | | H1001 | H4040 | | | SRR2040793 | SAMN03703466 |
| SRR2040794 | 2 | Scandinavia_UK_2n | | H2004 | H2007 | | | SRR2040794 | SAMN03703467 |
| SRR2040795 | 2 | Karelia_2n | | H1001 | H1001 | | | SRR2040795 | SAMN03703468 |
| SRR2040797 | 2 | Central_Europe_2n | 2x_Europe | H1001 | H1001 | | | SRR2040797 | SAMN03703470 |
| SRR2040798 | 2 | Central_Europe_2n | 2x_Europe | H2004 | H4031 | | | SRR2040798 | SAMN03703471 |
| SRR2040804 | 2 | East_Siberia_2n | 2x_Siberia | H4015 | H4039 | | | SRR2040804 | SAMN03703477 |
| SRR2040805 | 4 | Northern_Ural_4n | 4x_Siberia | H1001 | H2004 | H3001 | H4038 | SRR2040805 | SAMN03703478 |
| SRR2040826 | 4 | Central_Europe_4n | 4x_Europe | H1001 | H1001 | H1001 | H1001 | SRR2040826 | SAMN03703499 |
| SRR2040827 | 4 | Central_Europe_4n | 4x_Europe | H1001 | H1001 | H1001 | H1001 | SRR2040827 | SAMN03703500 |
| SRR2040828 | 4 | Central_Europe_4n | 4x_Europe | H4030 | H4030 | H3011 | H1001 | SRR2040828 | SAMN03703501 |
| SRR2040830 | 4 | Central_Europe_4n | 4x_Europe | H1001 | H4004 | H4034 | H4034 | SRR2040830 | SAMN03703503 |
| SRR3111439 | 2 | Central_Europe_2n | 2x_Europe | H1001 | H4013 | | | SRR3111439 | SAMN04432148 |
| SRR3111440 | 2 | Central_Europe_2n | 2x_Europe | H4010 | H4030 | | | SRR3111440 | SAMN04432149 |
| SRR3111441 | 2 | Central_Europe_2n | 2x_Europe | H3001 | H4038 | | | SRR3111441 | SAMN04432150 |
| SRR3111442 | 2 | Central_Europe_2n | 2x_Europe | H1001 | H3002 | | | SRR3111442 | SAMN04432151 |
| SRR3111443 | 2 | Central_Europe_2n | 2x_Europe | H4006 | H4027 | | | SRR3111443 | SAMN04432152 |
| SRR3111448 | 4 | Central_Europe_4n | 4x_Europe | H1001 | H1001 | H2003 | H2006 | SRR3111448 | SAMN04432157 |
| SRR5124974 | 2 | Karelia_2n | | H3003 | H4015 | | | SRR5124974 | SAMN06159139 |
| SRR5124975 | 2 | Central_Europe_2n | 2x_Europe | H1001 | H1001 | | | SRR5124975 | SAMN06141187 |
| SRR5124976 | 2 | Scandinavia_UK_2n | | H2008 | H4007 | | | SRR5124976 | SAMN06141186 |
| SRR5124977 | 2 | Scandinavia_UK_2n | | H1001 | H4010 | | | SRR5124977 | SAMN06141178 |
| SRR5124978 | 2 | Scandinavia_UK_2n | | H1001 | H3001 | | | SRR5124978 | SAMN06141181 |
| SRR5124979 | 2 | North_America_2n | | H1001 | H3008 | | | SRR5124979 | SAMN06141197 |
| SRR5124980 | 2 | Karelia_2n | | H3003 | H4023 | | | SRR5124980 | SAMN06159138 |
| SRR5124981 | 2 | North_America_2n | | H3008 | H3008 | | | SRR5124981 | SAMN06141194 |
| SRR5124982 | 2 | Scandinavia_UK_2n | | H3001 | H4020 | | | SRR5124982 | SAMN06141182 |
| SRR5124983 | 2 | Scandinavia_UK_2n | | H1001 | H3002 | | | SRR5124983 | SAMN06141175 |
| SRR5124984 | 2 | Central_Europe_2n | 2x_Europe | H2006 | H4024 | | | SRR5124984 | SAMN06141188 |

| | | | | | | | | | |
|---|---|---|---|---|---|---|---|---|---|
| SRR5124985 | 2 | Scandinavia_UK_2n | | H1001 | H3004 | | | SRR5124985 | SAMN06141176 |
| SRR5124986 | 2 | Central_Europe_2n | 2x_Europe | H1001 | H4018 | | | SRR5124986 | SAMN06141190 |
| SRR5124987 | 2 | Scandinavia_UK_2n | | H3001 | H4016 | | | SRR5124987 | SAMN06141180 |
| SRR5124988 | 2 | Scandinavia_UK_2n | | H3001 | H4007 | | | SRR5124988 | SAMN06141185 |
| SRR5124989 | 2 | Central_Europe_2n | 2x_Europe | H4005 | H4041 | | | SRR5124989 | SAMN06141191 |
| SRR5124990 | 2 | Central_Europe_2n | 2x_Europe | H1001 | H4002 | | | SRR5124990 | SAMN06141189 |
| SRR5124991 | 2 | North_America_2n | | H1001 | H4012 | | | SRR5124991 | SAMN06141193 |
| SRR5124992 | 2 | North_America_2n | | H4018 | H4035 | | | SRR5124992 | SAMN06141198 |
| SRR5124993 | 2 | Karelia_2n | | H1001 | H3002 | | | SRR5124993 | SAMN06159140 |
| SRR5124994 | 2 | Scandinavia_UK_2n | | H1001 | H3008 | | | SRR5124994 | SAMN06141195 |
| SRR5124995 | 2 | Scandinavia_UK_2n | | H1001 | H4001 | | | SRR5124995 | SAMN06141179 |
| SRR5124996 | 2 | Scandinavia_UK_2n | | H1001 | H2007 | | | SRR5124996 | SAMN06141183 |
| SRR5124997 | 2 | Scandinavia_UK_2n | | H1001 | H1001 | | | SRR5124997 | SAMN06141174 |
| SRR5124998 | 2 | Scandinavia_UK_2n | | H1001 | H4009 | | | SRR5124998 | SAMN06141177 |
| SRR5124999 | 2 | Scandinavia_UK_2n | | H1001 | H1001 | | | SRR5124999 | SAMN06141173 |
| SRR5125000 | 2 | North_America_2n | | H4008 | H4026 | | | SRR5125000 | SAMN06141196 |
| SRR5125001 | 2 | Karelia_2n | | H1001 | H4023 | | | SRR5125001 | SAMN06159141 |
| SRR5125002 | 2 | Scandinavia_UK_2n | | H2004 | H2007 | | | SRR5125002 | SAMN06141184 |
| SRR5125003 | 2 | Central_Europe_2n | 2x_Europe | H1001 | H1001 | | | SRR5125003 | SAMN06141192 |
| SRR7119523 | 2 | Scandinavia_UK_2n | | H4001 | H4004 | | | SRR7119523 | SAMN09069407 |
| SRR7119524 | 2 | Scandinavia_UK_2n | | H2004 | H4008 | | | SRR7119524 | SAMN09069408 |
| SRR7119525 | 2 | Scandinavia_UK_2n | | H1001 | H4029 | | | SRR7119525 | SAMN09069409 |
| SRR7119526 | 2 | Scandinavia_UK_2n | | H1001 | H4027 | | | SRR7119526 | SAMN09069410 |
| SRR7119527 | 2 | Scandinavia_UK_2n | | H1001 | H1001 | | | SRR7119527 | SAMN09069411 |
| SRR7119528 | 2 | Scandinavia_UK_2n | | H4022 | H4030 | | | SRR7119528 | SAMN09069412 |
| SRR7119529 | 2 | Scandinavia_UK_2n | | H1001 | H2004 | | | SRR7119529 | SAMN09069413 |
| SRR7119530 | 2 | Scandinavia_UK_2n | | H2008 | H4016 | | | SRR7119530 | SAMN09069414 |
| SRR7119531 | 2 | Scandinavia_UK_2n | | H1001 | H2008 | | | SRR7119531 | SAMN09069415 |
| SRR7119532 | 2 | Scandinavia_UK_2n | | H1001 | H2007 | | | SRR7119532 | SAMN09069416 |
| SRR7119533 | 2 | Scandinavia_UK_2n | | H1001 | H1001 | | | SRR7119533 | SAMN09069401 |
| SRR7119534 | 2 | Scandinavia_UK_2n | | H1001 | H4016 | | | SRR7119534 | SAMN09069402 |
| SRR7119535 | 2 | Scandinavia_UK_2n | | H3015 | H4004 | | | SRR7119535 | SAMN09069403 |
| SRR7119536 | 2 | Scandinavia_UK_2n | | H1001 | H4001 | | | SRR7119536 | SAMN09069404 |
| SRR7119537 | 2 | Scandinavia_UK_2n | | H1001 | H4001 | | | SRR7119537 | SAMN09069397 |
| SRR7119539 | 2 | Scandinavia_UK_2n | | H3011 | H4001 | | | SRR7119539 | SAMN09069399 |
| SRR7119540 | 2 | Scandinavia_UK_2n | | H1001 | H1001 | | | SRR7119540 | SAMN09069400 |
| SRR7119541 | 2 | Scandinavia_UK_2n | | H1001 | H1001 | | | SRR7119541 | SAMN09069405 |
| SRR7119542 | 2 | Scandinavia_UK_2n | | H1001 | H4027 | | | SRR7119542 | SAMN09069406 |
| SRR7119543 | 2 | Scandinavia_UK_2n | | H3010 | H3015 | | | SRR7119543 | SAMN09069422 |
| SRR7119544 | 2 | Scandinavia_UK_2n | | H1001 | H4010 | | | SRR7119544 | SAMN09069421 |
| SRR7119545 | 2 | Scandinavia_UK_2n | | H1001 | H4009 | | | SRR7119545 | SAMN09069418 |
| SRR7119546 | 2 | Scandinavia_UK_2n | | H3004 | H4034 | | | SRR7119546 | SAMN09069417 |
| SRR7119547 | 2 | Scandinavia_UK_2n | | H3004 | H4004 | | | SRR7119547 | SAMN09069420 |
| SRR7119548 | 2 | Scandinavia_UK_2n | | H1001 | H4001 | | | SRR7119548 | SAMN09069419 |
| SRR9319641 | 2 | Scandinavia_UK_2n | | H3004 | H4016 | | | SRR9319641 | SAMN12086031 |
| SRR9319642 | 2 | Scandinavia_UK_2n | | H1001 | H2008 | | | SRR9319642 | SAMN12086032 |
| SRR9319643 | 2 | Scandinavia_UK_2n | | H1001 | H2004 | | | SRR9319643 | SAMN12086033 |
| SRR9319644 | 2 | Scandinavia_UK_2n | | H4004 | H4010 | | | SRR9319644 | SAMN12086034 |
| SRR9319645 | 2 | Scandinavia_UK_2n | | H1001 | H3004 | | | SRR9319645 | SAMN12086027 |
| SRR9319646 | 2 | Scandinavia_UK_2n | | H4004 | H4030 | | | SRR9319646 | SAMN12086028 |
| SRR9319647 | 2 | Scandinavia_UK_2n | | H4004 | H4013 | | | SRR9319647 | SAMN12086029 |
| SRR9319648 | 2 | Scandinavia_UK_2n | | H1001 | H4008 | | | SRR9319648 | SAMN12086030 |
| SRR9319649 | 2 | Scandinavia_UK_2n | | H3004 | H4020 | | | SRR9319649 | SAMN12086035 |
| TE_10.1-1 | 2 | West_Siberia_2n | | H3002 | H3002 | | | ERR12742547 | ERS18382981 |
| TE_10.1-2 | 2 | West_Siberia_2n | | H3022 | H4001 | | | ERR12742546 | ERS18382982 |
| TE_10.2-1 | 2 | West_Siberia_2n | | H4016 | H4029 | | | ERR12742545 | ERS18382983 |
| TE_10.2-2 | 2 | West_Siberia_2n | | H4016 | H4041 | | | ERR12742544 | ERS18382984 |
| TE_10.3-1 | 2 | West_Siberia_2n | | H3022 | H4035 | | | ERR12742543 | ERS18382985 |
| TE_10.3-2 | 2 | West_Siberia_2n | | H1001 | H3022 | | | ERR11265614 | ERS14775782 |
| TE_10.4-1 | 2 | West_Siberia_2n | | H3022 | H4016 | | | ERR12742542 | ERS18382986 |
| TE_10.4-2 | 2 | West_Siberia_2n | | H3003 | H3022 | | | ERR12742541 | ERS18382987 |
| TE_10.5-1 | 2 | West_Siberia_2n | | H4001 | H4029 | | | ERR12742540 | ERS18382988 |
| TE_10.5-2 | 2 | West_Siberia_2n | | H4001 | H4041 | | | ERR12742539 | ERS18382989 |
| TE_10.6-1 | 2 | West_Siberia_2n | | H2003 | H4010 | | | ERR12742538 | ERS18382990 |
| TE_10.6-2 | 2 | West_Siberia_2n | | H1001 | H2003 | | | ERR12742537 | ERS18382991 |
| TE_10.7-1 | 2 | West_Siberia_2n | | H4001 | H4029 | | | ERR12742536 | ERS18382992 |
| TE_10.7-2 | 2 | West_Siberia_2n | | H2007 | H3003 | | | ERR12742535 | ERS18382993 |
| TE_10.8-1 | 2 | West_Siberia_2n | | H2003 | H4029 | | | ERR12742534 | ERS18382994 |
| TE_10.8-2 | 2 | West_Siberia_2n | | H2003 | H4017 | | | ERR12742533 | ERS18382995 |
| TE_10.9-1 | 2 | West_Siberia_2n | | H1001 | H4010 | | | ERR12742532 | ERS18382996 |

| | | | | | | | | | |
|---|---|---|---|---|---|---|---|---|---|
| TE_10.9-2 | 2 | West_Siberia_2n | | H1001 | H4010 | | | ERR12742531 | ERS18382997 |
| TE_11.1-1 | 2 | West_Siberia_2n | | H1001 | H3022 | | | ERR12742530 | ERS18382998 |
| TE_11.1-2 | 2 | West_Siberia_2n | | H1001 | H4034 | | | ERR11265613 | ERS14775783 |
| TE_11.10-1 | 2 | West_Siberia_2n | | H1001 | H1001 | | | ERR12742529 | ERS18382999 |
| TE_11.10-2 | 2 | West_Siberia_2n | | H1001 | H4034 | | | ERR12742528 | ERS18383000 |
| TE_11.2-1 | 2 | West_Siberia_2n | | H4002 | H4003 | | | ERR12742527 | ERS18383001 |
| TE_11.2-2 | 2 | West_Siberia_2n | | H4002 | H4034 | | | ERR12742526 | ERS18383002 |
| TE_11.3-1 | 2 | West_Siberia_2n | | H1001 | H1001 | | | ERR12742525 | ERS18383003 |
| TE_11.3-2 | 2 | West_Siberia_2n | | H3022 | H4034 | | | ERR12742524 | ERS18383004 |
| TE_11.4-1 | 2 | West_Siberia_2n | | H1001 | H4034 | | | ERR12742523 | ERS18383005 |
| TE_11.4-2 | 2 | West_Siberia_2n | | H4003 | H4021 | | | ERR12742522 | ERS18383006 |
| TE_11.5-1 | 2 | West_Siberia_2n | | H1001 | H3022 | | | ERR12742521 | ERS18383007 |
| TE_11.5-2 | 2 | West_Siberia_2n | | H1001 | H1001 | | | ERR12742520 | ERS18383008 |
| TE_11.6-1 | 2 | West_Siberia_2n | | H1001 | H4034 | | | ERR12742519 | ERS18383009 |
| TE_11.6-2 | 2 | West_Siberia_2n | | H1001 | H1001 | | | ERR12742518 | ERS18383010 |
| TE_11.7-1 | 2 | West_Siberia_2n | | H1001 | H3003 | | | ERR12742517 | ERS18383011 |
| TE_11.7-2 | 2 | West_Siberia_2n | | H1001 | H4034 | | | ERR12742516 | ERS18383012 |
| TE_11.8-1 | 2 | West_Siberia_2n | | H4002 | H4003 | | | ERR12742515 | ERS18383013 |
| TE_11.8-2 | 2 | West_Siberia_2n | | H4003 | H4013 | | | ERR12742514 | ERS18383014 |
| TE_11.9-1 | 2 | West_Siberia_2n | | H4013 | H4034 | | | ERR12742513 | ERS18383015 |
| TE_11.9-2 | 2 | West_Siberia_2n | | H4013 | H4034 | | | ERR12742512 | ERS18383016 |
| TE_3.1-1 | 2 | West_Siberia_2n | | H1001 | H3022 | | | ERR12742511 | ERS18383017 |
| TE_3.1-2 | 2 | West_Siberia_2n | | H3002 | H3022 | | | ERR12742510 | ERS18383018 |
| TE_3.2-1 | 2 | West_Siberia_2n | | H2003 | H4017 | | | ERR12742509 | ERS18383019 |
| TE_3.2-2 | 2 | West_Siberia_2n | | H3003 | H4037 | | | ERR12742508 | ERS18383020 |
| TE_3.3-1 | 2 | West_Siberia_2n | | H1001 | H3003 | | | ERR12742507 | ERS18383021 |
| TE_3.3-2 | 2 | West_Siberia_2n | | H1001 | H3003 | | | ERR12742506 | ERS18383022 |
| TE_3.4-1 | 2 | West_Siberia_2n | | H1001 | H4034 | | | ERR12742505 | ERS18383023 |
| TE_3.4-2 | 2 | West_Siberia_2n | | H4017 | H4010 | | | ERR12742504 | ERS18383024 |
| TE_3.5-1 | 2 | West_Siberia_2n | | H1001 | H3022 | | | ERR12742503 | ERS18383025 |
| TE_3.5-2 | 2 | West_Siberia_2n | | H1001 | H4034 | | | ERR12742502 | ERS18383026 |
| TE_3.6-1 | 2 | West_Siberia_2n | | H4002 | H3003 | | | ERR12742501 | ERS18383027 |
| TE_3.6-2 | 2 | West_Siberia_2n | | H4034 | H4041 | | | ERR12742500 | ERS18383028 |
| TE_3.7-1 | 2 | West_Siberia_2n | | H4016 | H4010 | | | ERR12742499 | ERS18383029 |
| TE_3.7-2 | 2 | West_Siberia_2n | | H3022 | H4010 | | | ERR12742498 | ERS18383030 |
| TE_3.8-1 | 2 | West_Siberia_2n | | H3003 | H4037 | | | ERR12742497 | ERS18383031 |
| TE_3.8-2 | 2 | West_Siberia_2n | | H1001 | H3003 | | | ERR12742496 | ERS18383032 |
| TE_3.9-1 | 2 | West_Siberia_2n | | H3003 | H3022 | | | ERR12742495 | ERS18383033 |
| TE_3.9-2 | 2 | West_Siberia_2n | | H3022 | H3022 | | | ERR12742494 | ERS18383034 |
| TE_4.10-1 | 2 | West_Siberia_2n | | H2003 | H2003 | | | ERR12742493 | ERS18383035 |
| TE_4.10-2 | 2 | West_Siberia_2n | | H2003 | H2003 | | | ERR12742492 | ERS18383036 |
| TE_4.11-1 | 2 | West_Siberia_2n | | H4012 | H4041 | | | ERR12742491 | ERS18383037 |
| TE_4.11-2 | 2 | West_Siberia_2n | | H1001 | H4012 | | | ERR12742490 | ERS18383038 |
| TE_4.2-1 | 2 | West_Siberia_2n | | H1001 | H3022 | | | ERR12742489 | ERS18383039 |
| TE_4.2-2 | 2 | West_Siberia_2n | | H3003 | H3022 | | | ERR12742488 | ERS18383040 |
| TE_4.3-1 | 2 | West_Siberia_2n | | H3002 | H4041 | | | ERR12742487 | ERS18383041 |
| TE_4.3-2 | 2 | West_Siberia_2n | | H4010 | H4037 | | | ERR12742486 | ERS18383042 |
| TE_4.4-1 | 2 | West_Siberia_2n | | H4010 | H4041 | | | ERR12742485 | ERS18383043 |
| TE_4.4-2 | 2 | West_Siberia_2n | | H4010 | H4037 | | | ERR12742484 | ERS18383044 |
| TE_4.5-1 | 2 | West_Siberia_2n | | H3002 | H4029 | | | ERR12742483 | ERS18383045 |
| TE_4.5-2 | 2 | West_Siberia_2n | | H3002 | H4010 | | | ERR12742482 | ERS18383046 |
| TE_4.6-1 | 2 | West_Siberia_2n | | H1001 | H2003 | | | ERR12742481 | ERS18383047 |
| TE_4.6-2 | 2 | West_Siberia_2n | | H3003 | H4010 | | | ERR12742480 | ERS18383048 |
| TE_4.7-1 | 2 | West_Siberia_2n | | H1001 | H4041 | | | ERR12742479 | ERS18383049 |
| TE_4.8-1 | 2 | West_Siberia_2n | | H4034 | H4012 | | | ERR12742478 | ERS18383050 |
| TE_4.8-2 | 2 | West_Siberia_2n | | H4012 | H4037 | | | ERR12742477 | ERS18383051 |
| TE_4.9-1 | 2 | West_Siberia_2n | | H3003 | H4029 | | | ERR12742476 | ERS18383052 |
| TE_4.9-2 | 2 | West_Siberia_2n | | H3002 | H3003 | | | ERR12742475 | ERS18383053 |
| TE_5.1-1 | 2 | West_Siberia_2n | | H3002 | H4034 | | | ERR12742474 | ERS18383054 |
| TE_5.1-2 | 2 | West_Siberia_2n | | H3002 | H3003 | | | ERR12742473 | ERS18383055 |
| TE_5.10-1 | 2 | West_Siberia_2n | | H4012 | H4025 | | | ERR12742472 | ERS18383056 |
| TE_5.10-2 | 2 | West_Siberia_2n | | H4002 | H4021 | | | ERR12742471 | ERS18383057 |
| TE_5.2-1 | 2 | West_Siberia_2n | | H1001 | H3002 | | | ERR12742470 | ERS18383058 |
| TE_5.2-2 | 2 | West_Siberia_2n | | H1001 | H3002 | | | ERR12742469 | ERS18383059 |
| TE_5.3-1 | 2 | West_Siberia_2n | | H1001 | H3003 | | | ERR12742468 | ERS18383060 |
| TE_5.3-2 | 2 | West_Siberia_2n | | H1001 | H3002 | | | ERR12742467 | ERS18383061 |
| TE_5.4-1 | 2 | West_Siberia_2n | | H1001 | H3003 | | | ERR12742466 | ERS18383062 |
| TE_5.4-2 | 2 | West_Siberia_2n | | H4010 | H4034 | | | ERR12742465 | ERS18383063 |
| TE_5.5-1 | 2 | West_Siberia_2n | | H2003 | H3002 | | | ERR12742464 | ERS18383064 |
| TE_5.5-2 | 2 | West_Siberia_2n | | H1001 | H3002 | | | ERR12742463 | ERS18383065 |

| | | | | | | | | | |
|---|---|---|---|---|---|---|---|---|---|
| TE_5.6-1 | 2 | West_Siberia_2n | | H4010 | H4012 | | | ERR12742462 | ERS18383066 |
| TE_5.6-2 | 2 | West_Siberia_2n | | H4010 | H4012 | | | ERR12742461 | ERS18383067 |
| TE_5.7-1 | 2 | West_Siberia_2n | | H1001 | H4004 | | | ERR12742460 | ERS18383068 |
| TE_5.7-2 | 2 | West_Siberia_2n | | H1001 | H4034 | | | ERR12742459 | ERS18383069 |
| TE_5.8-1 | 2 | West_Siberia_2n | | H1001 | H4016 | | | ERR12742458 | ERS18383070 |
| TE_5.8-2 | 2 | West_Siberia_2n | | H4012 | H4016 | | | ERR12742457 | ERS18383071 |
| TE_5.9-1 | 2 | West_Siberia_2n | | H4004 | H4035 | | | ERR12742456 | ERS18383072 |
| TE_5.9-2 | 2 | West_Siberia_2n | | H4004 | H4017 | | | ERR12742455 | ERS18383073 |
| TE_6.1-1 | 2 | West_Siberia_2n | | H1001 | H3002 | | | ERR12742454 | ERS18383074 |
| TE_6.1-2 | 2 | West_Siberia_2n | | H2003 | H3002 | | | ERR12742453 | ERS18383075 |
| TE_6.2-1 | 2 | West_Siberia_2n | | H3002 | H4034 | | | ERR12742452 | ERS18383076 |
| TE_6.2-2 | 2 | West_Siberia_2n | | H3002 | H4034 | | | ERR12742451 | ERS18383077 |
| TE_7.1-1 | 2 | West_Siberia_2n | | H1001 | H4003 | | | ERR12742450 | ERS18383078 |
| TE_7.1-2 | 2 | West_Siberia_2n | | H4002 | H4017 | | | ERR12742449 | ERS18383079 |
| TE_7.2-1 | 2 | West_Siberia_2n | | H3002 | H4041 | | | ERR12742448 | ERS18383080 |
| TE_7.2-2 | 2 | West_Siberia_2n | | H4016 | H4041 | | | ERR12742447 | ERS18383081 |
| TE_7.3-1 | 2 | West_Siberia_2n | | H1001 | H4041 | | | ERR12742446 | ERS18383082 |
| TE_7.3-2 | 2 | West_Siberia_2n | | H4034 | H4041 | | | ERR12742445 | ERS18383083 |
| TE_7.4-1 | 2 | West_Siberia_2n | | H4010 | H4041 | | | ERR12742444 | ERS18383084 |
| TE_7.4-2 | 2 | West_Siberia_2n | | H4010 | H4016 | | | ERR12742443 | ERS18383085 |
| TE_7.5-1 | 2 | West_Siberia_2n | | H1001 | H4002 | | | ERR12742442 | ERS18383086 |
| TE_7.5-2 | 2 | West_Siberia_2n | | H1001 | H4002 | | | ERR12742441 | ERS18383087 |
| TE_7.6-1 | 2 | West_Siberia_2n | | H1001 | H4010 | | | ERR12742440 | ERS18383088 |
| TE_7.6-2 | 2 | West_Siberia_2n | | H3004 | H3004 | | | ERR12742439 | ERS18383089 |
| TE_7.7-1 | 2 | West_Siberia_2n | | H3003 | H4041 | | | ERR12742438 | ERS18383090 |
| TE_7.7-2 | 2 | West_Siberia_2n | | H3003 | H4003 | | | ERR12742437 | ERS18383091 |
| TE_7.8-1 | 2 | West_Siberia_2n | | H4003 | H4041 | | | ERR12742436 | ERS18383092 |
| TE_7.8-2 | 2 | West_Siberia_2n | | H3004 | H4002 | | | ERR12742435 | ERS18383093 |
| TE_7.9-1 | 2 | West_Siberia_2n | | H1001 | H4025 | | | ERR12742434 | ERS18383094 |
| TE_7.9-2 | 2 | West_Siberia_2n | | H1001 | H4025 | | | ERR12742433 | ERS18383095 |
| TE_7mix-1 | 2 | West_Siberia_2n | | H3003 | H4041 | | | ERR12742432 | ERS18383096 |
| TE_7mix-2 | 2 | West_Siberia_2n | | H1001 | H4016 | | | ERR12742431 | ERS18383097 |
| TE_8.1-1 | 2 | West_Siberia_2n | | H2009 | H2009 | | | ERR12742430 | ERS18383098 |
| TE_8.1-2 | 2 | West_Siberia_2n | | H2009 | H3003 | | | ERR12742429 | ERS18383099 |
| TE_8.10-1 | 2 | West_Siberia_2n | | H2003 | H3022 | | | ERR12742428 | ERS18383100 |
| TE_8.10-2 | 2 | West_Siberia_2n | | H2003 | H3003 | | | ERR12742427 | ERS18383101 |
| TE_8.2-1 | 2 | West_Siberia_2n | | H2009 | H3003 | | | ERR12742426 | ERS18383102 |
| TE_8.2-2 | 2 | West_Siberia_2n | | H2009 | H4004 | | | ERR12742425 | ERS18383103 |
| TE_8.3-1 | 2 | West_Siberia_2n | | H2009 | H3003 | | | ERR12742424 | ERS18383104 |
| TE_8.3-2 | 2 | West_Siberia_2n | | H2009 | H4016 | | | ERR12742423 | ERS18383105 |
| TE_8.4-1 | 2 | West_Siberia_2n | | H1001 | H2009 | | | ERR12742422 | ERS18383106 |
| TE_8.4-2 | 2 | West_Siberia_2n | | H1001 | H2003 | | | ERR12742421 | ERS18383107 |
| TE_8.5-1 | 2 | West_Siberia_2n | | H1001 | H3003 | | | ERR12742420 | ERS18383108 |
| TE_8.5-2 | 2 | West_Siberia_2n | | H1001 | H4035 | | | ERR12742419 | ERS18383109 |
| TE_8.6-1 | 2 | West_Siberia_2n | | H3003 | H3022 | | | ERR12742418 | ERS18383110 |
| TE_8.6-2 | 2 | West_Siberia_2n | | H3022 | H3022 | | | ERR12742417 | ERS18383111 |
| TE_8.7-1 | 2 | West_Siberia_2n | | H2009 | H2009 | | | ERR12742416 | ERS18383112 |
| TE_8.7-2 | 2 | West_Siberia_2n | | H2009 | H4016 | | | ERR12742415 | ERS18383113 |
| TE_8.8-1 | 2 | West_Siberia_2n | | H2009 | H3022 | | | ERR12742414 | ERS18383114 |
| TE_8.8-2 | 2 | West_Siberia_2n | | H2003 | H3003 | | | ERR12742413 | ERS18383115 |
| TE_8.9-1 | 2 | West_Siberia_2n | | H2003 | H3022 | | | ERR12742412 | ERS18383116 |
| TE_8.9-2 | 2 | West_Siberia_2n | | H2003 | H2009 | | | ERR12742411 | ERS18383117 |
| TE_8mix-1 | 2 | West_Siberia_2n | | H2007 | H2009 | | | ERR12742410 | ERS18383118 |
| TE_8mix-2 | 2 | West_Siberia_2n | | H2009 | H2009 | | | ERR12742409 | ERS18383119 |
| TE_9.1-1 | 2 | West_Siberia_2n | | H3022 | H4027 | | | ERR12742408 | ERS18383120 |
| TE_9.1-2 | 2 | West_Siberia_2n | | H3022 | H3022 | | | ERR12742407 | ERS18383121 |
| WS_1.1-1 | 4 | Northern_Ural_4n | 4x_Siberia | H1001 | H1001 | H1001 | H2008 | ERR12742406 | ERS18383122 |
| WS_1.1-2 | 4 | Northern_Ural_4n | 4x_Siberia | H1001 | H1001 | H3022 | H3022 | ERR12742405 | ERS18383123 |
| WS_1.2-1 | 4 | Northern_Ural_4n | 4x_Siberia | H1001 | H1001 | H1001 | H3011 | ERR12742404 | ERS18383124 |
| WS_1.2-2 | 4 | Northern_Ural_4n | 4x_Siberia | H1001 | H1001 | H1001 | H1001 | ERR12742403 | ERS18383125 |
| WS_1.3-1 | 4 | Northern_Ural_4n | 4x_Siberia | H1001 | H1001 | H1001 | H2007 | ERR12742402 | ERS18383126 |
| WS_1.3-2 | 4 | Northern_Ural_4n | 4x_Siberia | H4016 | H4016 | H1001 | - | ERR12742401 | ERS18383127 |
| WS_1.4-1 | 4 | Northern_Ural_4n | 4x_Siberia | H1001 | H1001 | H2007 | H3011 | ERR12742400 | ERS18383128 |
| WS_1.4-2 | 4 | Northern_Ural_4n | 4x_Siberia | H1001 | H1001 | H2004 | H3011 | ERR12742399 | ERS18383129 |
| WS_1.5-1 | 4 | Northern_Ural_4n | 4x_Siberia | H1001 | H1001 | H1001 | H1001 | ERR12742398 | ERS18383130 |
| WS_1.5-2 | 4 | Northern_Ural_4n | 4x_Siberia | H1001 | H2008 | H3011 | H4010 | ERR12742397 | ERS18383131 |
| WS_1.6-1 | 4 | Northern_Ural_4n | 4x_Siberia | H1001 | H1001 | H1001 | H2003 | ERR12742396 | ERS18383132 |
| WS_1.6-2 | 4 | Northern_Ural_4n | 4x_Siberia | H1001 | H1001 | H1001 | H1001 | ERR12742395 | ERS18383133 |
| WS_1.7-1 | 4 | Northern_Ural_4n | 4x_Siberia | H1001 | H1001 | H1001 | H1001 | ERR12742394 | ERS18383134 |
| WS_1.7-2 | 4 | Northern_Ural_4n | 4x_Siberia | H1001 | H2003 | H3011 | H4041 | ERR12742393 | ERS18383135 |

**Table S3.** List of the 83 *SRK* alleles identified in *A. arenosa* , with correspondence to previously identified alleles in A. arenosa (Genbank accession and original source given), and functional correspondence to *A. halleri* , *A. lyrata* , and *Capsella grandiflora* alleles, based on phylogenetic relationships. The individual used to extract the full S-domain sequence (exon 1) of each *A. arenosa SRK* allele using the NGSgenotyp pipeline is indicated. The Allele ID corresponds to a new identification scheme based on phylogenetic relationships among *SRK* sequences from different species, in the form HXYYY, with X corresponding to the dominance class of the allele (ranging from 1 to 4 according to Goubet et al. 2012), and YYY corresponding to its functional specificity shared among species and genera (ranging from 001 up to a limit of 999).

| Allele ID (functional group) | *A. arenosa* specific allele ID | Genbank (partial sequence) | Original source | Individual used to extract full sequence | *A. halleri* homolog | *A. lyrata* homolog | *Capsella grandiflora* homolog |
|---|---|---|---|---|---|---|---|
| H1001 | AarSRK01 | MH507385 | Mable et al. 2018 | BRD_04ta_Contig_N5 | AhSRK01 | AlSRK01 | CgrSRK03 |
| H2001 | AarSRK03 | JX464612 | Ruiz-Duarte 2012 | CAR_08ta_Contig_N2 | AhSRK03 | AlSRK03 | |
| H2003 | AarSRK14 | JX464617 | Ruiz-Duarte 2012 | ZID_2da_Contig_N4 | AhSRK09 | AlSRK14 | |
| H2004 | AarSRK08 | JX464614 | Ruiz-Duarte 2012 | BAL_03ta_Contig_N7 | AhSRK19 | AlSRK08 | |
| H2005 | AarSRKH2005 | | this study | FOJ_03da_Contig_N2 | AhSRK23 | | CgrSRK08 |
| H2006 | AarSRK28 | | this study | INE_08ta_Contig_N2 | AhSRK28 | AlSRK28 | CgrSRK24 |
| H2007 | AarSRK18 | | this study | BAB_04da_Contig_N4 | AhSRK33 | AlSRK18 | |
| H2008 | AarSRK06 | JX464613 | Ruiz-Duarte 2012 | BOR_01ta_Contig_N5 | | AlSRK06 | CgrSRK38 |
| H2009 | AarSRK29 | MH507379 | AarSRK18 Mable et al. 2018 | BUT_05ta_Contig_N5 | AhSRK45 | AlSRK29 | CgrSRK04 |
| H2010 | AarSRKH2010 | JX464632 | AarSRK69 Ruiz-Duarte 2012 | HMC_03da_Contig_N8 | AhSRK69 | | CgrSRK39 |
| H2011 | AarSRKH2011 | MH507397 | AarSRK79 Mable et al. 2018 | VLA_06ta_Contig_N11 | | | CgrSRK34 |
| H2018 | AarSRK74 | | this study | INE_06ta_Contig_N7 | AhSRK68 | AlSRK74 | CgrSRK45 |
| H2019 | AarSRKH2019 | | this study | HRA_13ta_Contig_N7 | | | CgrSRKH2019 |
| H2021 | AarSRKH2021 | | this study | SUB_05da_Contig_N6 | | | CgrSRK05 |
| H2022 | AarSRKH2022 | MH507394 | AarSRK78 Mable et al. 2018 | SUB_01da_Contig_N9 | | | CgrSRK27 |
| H2024 | AarSRKH2024 | | this study | HRA_04ta_Contig_N4 | | | CgrSRK40 |
| H2025 | AarSRKH2025 | | this study | CAR_08ta_Contig_N6 | | | |
| H2026 | AarSRKH2026 | JX464633 | AarSRK74 Mable et al. 2018 | FUG_07ta_Contig_N7 | | | |
| H2027 | AarSRKH2027 | MH507389 | AarSRK26 Mable et al. 2019 | PAD_07da_Contig_N6 | | | |
| H2028 | AarSRKH2028 | MH507396 | AarSRK76 Mable et al. 2018 | ZIT_4ta_Contig_N5 | | AlSRKH2028 | |
| H2029 | AarSRKH2029 | | this study | ZAP_07ta_Contig_N15 | | | |
| H3001 | AarSRK17 | MH507386 | Mable et al. 2018 | RZA_08da_Contig_N5 | AhSRK02 | AlSRK17 | CgrSRK18 |
| H3002 | AarSRK37 | MH507392 | Mable et al. 2018 | BIH_06da_Contig_N10 | AhSRK04 | AlSRK37 | CgrSRK17 |
| H3003 | AarSRK21 | MH507387 | Mable et al. 2018 | BAB_10da_Contig_N8 | AhSRK06 | AlSRK21 | CgrSRK12 |
| H3004 | AarSRK16 | JX464620 | Ruiz-Duarte 2012 | MIE_11da_Contig_N20 | AhSRK10 | AlSRK16 | |
| H3005 | AarSRKH3005 | | this study | BRD_03ta_Contig_N25 | AhSRK14 | | CgrSRK32 |
| H3006 | AarSRKH3006 | MH507384 | AarSRK84 Mable et al. 2018 | HMC_01da_Contig_N5 | AhSRK17 | AlSRKH3006 | CgrSRKH3006 |
| H3007 | AarSRKH3007 | | this study | HRA_04ta_Contig_N1 | AhSRK25 | | CgrSRKH3007 |
| H3008 | AarSRK13 | | this study | PHD_02da_Contig_N7 | AhSRK29 | AlSRK13 | CgrH3008 |
| H3009 | AarSRKH3009 | | this study | BAB_10da_Contig_N6 | AhSRK30 | AlSRKH3009 | |
| H3010 | AarSRK05 | | this study | RZA_06da_Contig_N3 | | AlSRK05 | CgrSRK46 |
| H3011 | AarSRK25 | | this study | PAD_08da_Contig_N7 | AhSRK57 | AlSRK25 | CgrSRK53 |
| H3012 | AarSRKH3012 | | this study | OPP_07ta_Contig_N8 | AhSRK60 | AlSRKH3012 | CgrSRK11 |
| H3014 | AarSRKH3014 | | this study | BIH_02da_Contig_N8 | AhSRK58 | AlSRKH3014 | CgrSRK19 |
| H3015 | AarSRK70 | | this study | GOR_01da_Contig_N6 | AhSRK70 | AlSRK70 | CgrSRK20 |
| H3021 | AarSRKH3021 | | this study | HRA_02ta_Contig_N2 | AhSRK53 | AlSRKH3021 | |
| H3022 | AarSRK41 | | this study | CAR_01ta_Contig_N10 | AhSRK63 | AlSRK41 | CgrSRKH3022 |
| H3023 | AarSRK48 | | this study | TRD_06da_Contig_N6 | | AlSRK48 | CgrSRKH3023 |
| H3024 | AarSRK60 | | this study | BEL_05da_Contig_N8 | | AlSRK60 | |
| H3025 | AarSRKH3025 | MH507398 | AarSRK80 Mable et al. 2018 | BAL_07ta_Contig_N4 | | | |
| H3028 | AarSRKH3028 | | this study | BAB_06da_Contig_N7 | AhSRK65 | | |
| H4001 | AarSRK34 | MH507390 | Mable et al. 2018 | RZA_05da_Contig_N7 | AhSRK05 | AlSRK34 | CgrSRK41 |
| H4002 | AarSRK72 | MH507388 | AarSRK24 Mable et al. 2018 | BIH_08da_Contig_N5 | AhSRK07 | AlSRK72 | CgrSRK60 |
| H4003 | AarSRK11 | JX464616 | Ruiz-Duarte 2012 | VEL_08da_Contig_N10 | AhSRK11 | AlSRK11 | CgrSRKH4003 |
| H4004 | AarSRK42 | MH507382 | Mable et al. 2018 | BUD_04da_Contig_N8 | AhSRK12 | AlSRK42 | CgrSRK16 |
| H4005 | AarSRK66 | | this study | BDO_01da_Contig_N9 | AhSRK13 | AlSRK66 | CgrSRK55 |
| H4006 | AarSRK71 | | this study | BAB_03da_Contig_N5 | AhSRK15 | AlSRK71 | CgrSRKH4006 |
| H4007 | AarSRK31 | | this study | FOJ_01da_Contig_N10 | AhSRK16 | AlSRK31 | CgrSRK48 |
| H4008 | AarSRK39 | | this study | BIH_05da_Contig_N7 | AhSRK18 | AlSRK39 | CgrSRK15 |
| H4009 | AarSRK69 | | this study | PRE_04da_Contig_N7 | AhSRK20 | AlSRK69 | CgrSRK58 |
| H4010 | AarSRK15 | | this study | KAM_05ta_Contig_N5 | AhSRK21 | AlSRK15 | CgrSRK26 |
| H4011 | AarSRK46 | | this study | CRO_001_be_Contig_N5 | AhSRK22 | AlSRK46 | CgrSRK61 |
| H4012 | AarSRK20 | | this study | BUD_05da_Contig_N8 | AhSRK24 | AlSRK20 | CgrSRK29 |
| H4013 | AarSRK22 | | this study | BEL_08da_Contig_N9 | AhSRK26 | AlSRK22 | CgrSRKH4013 |
| H4014 | AarSRK68 | | this study | TRD_04da_Contig_N4 | AhSRK32 | AlSRK68 | CgrSRKH4014 |

| | | | | | | | |
|---|---|---|---|---|---|---|---|
| H4015 | AarSRK09 | | this study | MAR_01ta_Contig_N3 | AhSRK34 | AlSRK09 | CgrSRK62 |
| H4016 | AarSRK10 | | this study | SUB_04da_Contig_N7 | AhSRK35 | AlSRK10 | |
| H4017 | AarSRK36 | | this study | BEL_07da_Contig_N8 | AhSRK36 | AlSRK36 | CgrSRKH4017 |
| H4018 | AarSRK27 | | this study | BAB_08da_Contig_N3 | AhSRK37 | AlSRK27 | CgrSRKH4018 |
| H4019 | AarSRK65 | JX464627 | Ruiz-Duarte 2012 | MAR_01ta_Contig_N2 | AhSRK38 | AlSRK65 | CgrSRK35 |
| H4020 | AarSRK35 | MH507391 | Mable et al. 2018 | BEL_05da_Contig_N5 | AhSRK39 | AlSRK35 | CgrSRK02 |
| H4021 | AarSRKH4021 | | this study | MIE_03da_Contig_N12 | AhSRK40 | AlSRKH4021 | CgrSRK56 |
| H4022 | AarSRK67 | MH507395 | AarSRK77 Mable et al. 2018 | HNE_01da_Contig_N12 | AhSRK41 | AlSRK67 | |
| H4023 | AarSRK43 | | this study | KZL_05da_Contig_N7 | AhSRK43 | AlSRK43 | CgrSRK14 |
| H4024 | AarSRK63 | | this study | RZA_05da_Contig_N8 | AhSRK47 | AlSRK63 | CgrSRK21 |
| H4025 | AarSRK04 | JX464631 | Ruiz-Duarte 2012 | KAM_05ta_Contig_N10 | AhSRK71 | AlSRK04 | CgrSRKH4025 |
| H4026 | AarSRK23 | | this study | BUD_06da_Contig_N6 | AhSRK42 | AlSRK23 | CgrSRK31 |
| H4027 | AarSRK30 | | this study | OPP_04ta_Contig_N2 | AhSRK62 | AlSRK30 | CgrSRK37 |
| H4028 | AarSRK33 | | this study | GOR_07da_Contig_N6 | AhSRK51 | AlSRK33 | CgrSRKH4028 |
| H4029 | AarSRK38 | | this study | PRE_08da_Contig_N2 | AhSRK64 | AlSRK38 | CgrSRKH4029 |
| H4030 | AarSRK50 | | this study | KRM_06da_Contig_N7 | AhSRK50 | AlSRK50 | CgrSRKH4030 |
| H4031 | AarSRKH4031 | | this study | BAB_12da_Contig_N3 | AhSRK61 | AlSRKH4031 | CgrSRK28 |
| H4034 | AarSRK73 | MH507393 | AarSRK50 Mable et al. 2018 | BEL_04da_Contig_N9 | AhSRK54 | AlSRK73 | CgrSRK59 |
| H4035 | AarSRK19 | | this study | BIH_05da_Contig_N8 | AhSRK31 | AlSRK19 | CgrSRK22 |
| H4036 | AarSRK12 | | this study | CAR_03ta_Contig_N3 | AhSRK55 | AlSRK12 | CgrSRK54 |
| H4037 | AarSRK45 | | this study | KAM_08ta_Contig_N5 | AhSRK66 | AlSRK45 | CgrSRKH4037 |
| H4038 | AarSRK44 | | this study | INE_08ta_Contig_N8 | AhSRK52 | AlSRK44 | CgrSRKH4038 |
| H4039 | AarSRK61 | | this study | VEL_01da_Contig_N10 | AhSRK56 | AlSRK61 | CgrSRK57 |
| H4040 | AarSRK62 | | this study | BDO_04da_Contig_N10 | AhSRK44 | AlSRK62 | CgrSRKH4040 |
| H4041 | AarSRK64 | | this study | BUD_02da_Contig_N6 | AhSRK67 | AlSRK64 | |
| H4042 | AarSRKH4042 | | this study | OPP_03ta_Contig_N6 | AhSRK46 | AlSRKH4042 | CgrSRKH4042 |
| H4043 | AarSRKH4043 | | this study | SUB_04da_Contig_N5 | AhSRK59 | AlSRKH4043 | |
| H4048 | AarSRKH4048 | | this study | BEL_03da_Contig_N9 | | | |

**Table S4**. List of the 66 *SRK* alleles identified in *A. lyrata*, with correspondence to previously identified alleles in *A. lyrata* (Genbank accession and original source given), and functional correspondence to *A. halleri*, *A. arenosa*, and *Capsella grandiflora* alleles, based on phylogenetic relationships. The individual used to extract the full S-domain sequence (exon 1) of the new *A. lyrata SRK* alleles using the NGSgenotyp pipeline is indicated. The Allele ID corresponds to a new identification scheme based on phylogenetic relationships among SRK sequences from different species, in the form HXYYY, with X corresponding to the dominance class of the allele (ranging from I to IV according to Goubet et al. 2012), and YYY corresponding to its functional specificity shared among species and genera (ranging from 001 up to a limit of 999).

| Allele ID (functional group) | *A. lyrata* specific allele ID | Genbank (partial sequence) | Original source | Individual used to extract full sequence | *A. halleri* homolog | *A. arenosa* homolog | *Capsella grandiflora* homolog |
|---|---|---|---|---|---|---|---|
| H1001 | AlSRK01 | KJ772402 | Goubet et al. 2012 | | AhSRK01 | AarSRK01 | CgrSRK03 |
| H2001 | AlSRK03 | AF328992 | Schierup et al. 2001 | | AhSRK03 | AarSRK03 | |
| H2003 | AlSRK14 | KJ772405 | Goubet et al. 2012 | | AhSRK09 | AarSRK14 | |
| H2004 | AlSRK08 | AY186766 | Charlesworth et al. 2003 | | AhSRK19 | AarSRK08 | |
| H2006 | AlSRK28 | JX464655 | Ruiz-Duarte 2012 | | AhSRK28 | AarSRK28 | CgrSRK24 |
| H2007 | AlSRK18 | KJ772412 | Goubet et al. 2012 | | AhSRK33 | AarSRK18 | |
| H2008 | AlSRK06 | GQ351354 | Boggs et al. 2009 | | | AarSRK06 | CgrSRK38 |
| H2009 | AlSRK29 | AY186776 | Charlesworth et al. 2003 | | AhSRK45 | AarSRK29 | CgrSRK04 |
| H2018 | AlSRK74 | JX464649 | Ruiz-Duarte 2012 | ERR3514874_Contig_N22 | AhSRK68 | AarSRKH74 | CgrSRK45 |
| H2028 | AlSRKH2028 | | this study | PU_6.8-2_Contig_N26 | | AarSRKH2028 | |
| H3001 | AlSRK17 | AY186772 | Charlesworth et al. 2003 | | AhSRK02 | AarSRK17 | CgrSRK18 |
| H3002 | AlSRK37 | DQ520289 | Bechsgaard et al. 2006 | | AhSRK04 | AarSRK37 | CgrSRK17 |
| H3003 | AlSRK21 | EU878016 | Castric et al. 2008 | | AhSRK06 | AarSRK21 | CgrSRK12 |
| H3004 | AlSRK16 | HQ379629 | Guo et al. 2011 | | AhSRK10 | AarSRK16 | |
| H3006 | AlSRKH3006 | | this study | MW0079609_4_Contig_N7 | AhSRK17 | AarSRKH3006 | CgrSRKH3006 |
| H3008 | AlSRK13 | XM_002869844 | Hu et al. 2011 | | AhSRK29 | AarSRK13 | CgrH3008 |
| H3009 | AlSRKH3009 | | this study | MW0079568-1_1_Contig_N37 | AhSRK30 | AarSRKH3009 | |
| H3010 | AlSRK05 | AF328990 | Schierup et al. 2001 | | | AarSRK05 | CgrSRK46 |
| H3011 | AlSRK25 | GQ351355 | Boggs et al. 2009 | | AhSRK57 | AarSRK25 | CgrSRK53 |
| H3012 | AlSRKH3012 | | this study | NT17.1-4_Contig_N10 | AhSRK60 | AarSRKH3012 | CgrSRK11 |
| H3014 | AlSRKH3014 | | this study | NT15_3_1_Contig_N15 | AhSRK58 | AarSRKH3014 | CgrSRK19 |
| H3015 | AlSRK70 | SRS3258660 | Takou et al. 2021 | | AhSRK70 | AarSRK70 | CgrSRK20 |
| H3021 | AlSRKH3021 | | this study | PU_4mix-2_Contig_N15 | AhSRK53 | AarSRKH3021 | |
| H3022 | AlSRK41 | MH507372 | Mable et al. 2018 | | AhSRK63 | AarSRK41 | CgrSRKH3022 |
| H3024 | AlSRK60 | JX464644 | Ruiz-Duarte 2012 | | | AarSRK60 | |
| H4001 | AlSRK34 | EU878020 | Castric et al. 2008 | | AhSRK05 | AarSRK34 | CgrSRK41 |
| H4002 | AlSRK72 | ERS4653247 | Takou et al. 2021 | | AhSRK07 | AarSRK72 | CgrSRK60 |
| H4003 | AlSRK11 | AY186768 | Charlesworth et al. 2003 | | AhSRK11 | AarSRK11 | CgrSRKH4003 |
| H4004 | AlSRK42 | EU878023 | Castric et al. 2008 | | AhSRK12 | AarSRK42 | CgrSRK16 |
| H4005 | AlSRK66 | SRS1874397 | Takou et al. 2021 | | AhSRK13 | AarSRK66 | CgrSRK55 |
| H4006 | AlSRK71 | ERS3736358 | Takou et al. 2021 | | AhSRK15 | AarSRK71 | CgrSRKH4006 |
| H4007 | AlSRK31 | DQ520287 | Bechsgaard et al. 2006 | | AhSRK16 | AarSRK31 | CgrSRK48 |
| H4008 | AlSRK39 | KJ772418 | Goubet et al. 2012 | | AhSRK18 | AarSRK39 | CgrSRK15 |
| H4009 | AlSRK69 | SRS3258658 | Takou et al. 2021 | | AhSRK20 | AarSRK69 | CgrSRK58 |
| H4010 | AlSRK15 | KC207413 | Dwyer et al. 2013 | | AhSRK21 | AarSRK15 | CgrSRK26 |
| H4011 | AlSRK46 | MH507373 | Mable et al. 2018 | | AhSRK22 | AarSRK46 | CgrSRK61 |
| H4012 | AlSRK20 | AF328995 | Schierup et al. 2001 | | AhSRK24 | AarSRK20 | CgrSRK29 |
| H4013 | AlSRK22 | DQ520285 | Bechsgaard et al. 2006 | | AhSRK26 | AarSRK22 | CgrSRKH4013 |
| H4014 | AlSRK68 | JX464654 | Ruiz-Duarte 2012 | | AhSRK32 | AarSRK68 | CgrSRKH4014 |
| H4015 | AlSRK09 | DQ520282 | Bechsgaard et al. 2006 | | AhSRK34 | AarSRK09 | CgrSRK62 |
| H4016 | AlSRK10 | AY186767 | Charlesworth et al. 2003 | | AhSRK35 | AarSRK10 | |
| H4017 | AlSRK36 | KC207416 | Dwyer et al. 2013 | | AhSRK36 | AarSRK36 | CgrSRKH4017 |
| H4018 | AlSRK27 | EU878017 | Castric et al. 2008 | | AhSRK37 | AarSRK27 | CgrSRKH4018 |
| H4019 | AlSRK65 | ERS4653252 | Takou et al. 2021 | | AhSRK38 | AarSRK65 | CgrSRK35 |
| H4020 | AlSRK35 | EU878021 | Castric et al. 2008 | | AhSRK39 | AarSRK35 | CgrSRK02 |
| H4021 | AlSRKH4021 | | this study | ERR3514896_Contig_N14 | AhSRK40 | AarSRKH4021 | CgrSRK56 |
| H4022 | AlSRK67 | ERS3541323 | Takou et al. 2021 | | AhSRK41 | AarSRK67 | |
| H4023 | AlSRK43 | EU878024 | Castric et al. 2008 | | AhSRK43 | AarSRK43 | CgrSRK14 |
| H4024 | AlSRK63 | ERS4653250 | Takou et al. 2021 | | AhSRK47 | AarSRK63 | CgrSRK21 |
| H4025 | AlSRK04 | AF328994 | Schierup et al. 2001 | | AhSRK71 | AarSRK04 | CgrSRKH4025 |
| H4026 | AlSRK23 | AF328997 | Schierup et al. 2001 | | AhSRK42 | AarSRK23 | CgrSRK31 |
| H4027 | AlSRK30 | FJ867321.1 | Guo et al. 2009 | | AhSRK62 | AarSRK30 | CgrSRK37 |
| H4028 | AlSRK33 | EU878019 | Castric et al. 2008 | | AhSRK51 | AarSRK33 | CgrSRKH4028 |
| H4029 | AlSRK38 | HQ379630 | Guo et al. 2011 | | AhSRK64 | AarSRK38 | CgrSRKH4029 |
| H4030 | AlSRK50 | HQ379631 | Guo et al. 2011 | | AhSRK50 | AarSRK50 | CgrSRKH4030 |

| | | | | | | | |
|---|---|---|---|---|---|---|---|
| H4031 | AlSRKH4031 | | this study | PU_6.5-2_Contig_N1 | AhSRK61 | AarSRKH4031 | CgrSRK28 |
| H4034 | AlSRK73 | ERS3541316 | Takou et al. 2021 | | AhSRK54 | AarSRK73 | CgrSRK59 |
| H4035 | AlSRK19 | AF328998 | Schierup et al. 2001 | | AhSRK31 | AarSRK19 | CgrSRK22 |
| H4036 | AlSRK12 | AY186769 | Charlesworth et al. 2003 | | AhSRK55 | AarSRK12 | CgrSRK54 |
| H4037 | AlSRK45 | EU878026 | Castric et al. 2008 | | AhSRK66 | AarSRK45 | CgrSRKH4037 |
| H4038 | AlSRK44 | EU878025 | Castric et al. 2008 | | AhSRK52 | AarSRK44 | CgrSRKH4038 |
| H4039 | AlSRK61 | ERS4653249 | Takou et al. 2021 | | AhSRK56 | AarSRK61 | CgrSRK57 |
| H4040 | AlSRK62 | ERS4653250 | Takou et al. 2021 | | AhSRK44 | AarSRK62 | CgrSRKH4040 |
| H4041 | AlSRK64 | ERS4653251 | Takou et al. 2021 | | AhSRK67 | AarSRK64 | |
| H4042 | AlSRKH4042 | | this study | MW0079553_5_Contig_N8 | AhSRK46 | AarSRKH4042 | CgrSRKH4042 |
| H4043 | AlSRKH4043 | | this study | NT20.1-1_Contig_N9 | AhSRK59 | AarSRKH4043 | |

**Table S5.** Estimation of the number of alleles, gene diversity and observed heterozygosity at the S-locus in *Arabidopsis arenosa* in regional samples from the diploid and tetraploid gene pools.

| Regional sample* | Number of individuals | Number of S-alleles | Allelic richness (among 10 gene copies) | Gene diversity ($H_e$) | Observed heterozygosity ($H_o$) |
|---|---|---|---|---|---|
| 2x_WCa | 80 | 65 | 8.91 | 0.970 | 0.962 |
| 2x_ECa | 25 | 31 | 7.48 | 0.887 | 0.840 |
| 2x_SCa | 15 | 23 | 9.13 | 0.979 | 1.000 |
| 2x_Pan | 24 | 35 | 8.90 | 0.968 | 0.917 |
| 2x_Din | 30 | 35 | 8.94 | 0.972 | 1.000 |
| 2x_Bal | 32 | 31 | 8.45 | 0.959 | 1.000 |
| ***Mean 2X ±S.D.*** | ***34.3 ±23.1*** | ***36.7 ±14.6*** | ***8.64 ±0.61*** | ***0.956 ±0.035*** | ***0.953 ±0.064*** |
| 4x_WCa | 70 | 56 | 5.62 | 0.728 | 0.731 |
| 4x_ECa | 22 | 34 | 6.51 | 0.820 | 0.780 |
| 4x_SCa | 49 | 51 | 6.02 | 0.771 | 0.748 |
| 4x_Her | 42 | 48 | 5.72 | 0.740 | 0.750 |
| 4x_SWA | 25 | 22 | 4.29 | 0.567 | 0.547 |
| 4x_ALP | 88 | 59 | 5.14 | 0.672 | 0.652 |
| 4x_RUD | 45 | 27 | 4.49 | 0.600 | 0.570 |
| ***Mean 4X ±S.D.*** | ***48.7 ±23.6*** | ***42.4 ±14.7*** | ***5.40 ±0.81*** | ***0.700 ±0.091*** | ***0.683 ±0.094*** |

* WCa: Western Carpathian; ECa: East Carpathian; SCa: South Carpathian; Pan: Pannonian; Din: Dinaric; Bal: Baltic; Her: Hercynian; SWA: Swabian; ALP: Alpine; RUD: Ruderal. From Monnahan et al. (2019).

**Table S6.** Estimation of the number of alleles, gene diversity and observed heterozygosity at the S-locus in *Arabidopsis lyrata* in regional samples from the diploid and tetraploid gene pools.

| Regional sample | Number of individuals | Number of S-alleles | Allelic richness (among 10 gene copies) | Gene diversity ($H_e$) | Observed heterozygosity ($H_o$) |
|---|---|---|---|---|---|
| 2x_Siberia | 132 | 55 | 8.48 | 0.955 | 0.871 |
| 2x_Europe | 32 | 29 | 7.41 | 0.901 | 0.812 |
| ***Mean 2X ±S.D.*** | | ***42.0 ±18.4*** | ***7.95 ±0.76*** | ***0.928 ±0.038*** | ***0.842 ±0.042*** |
| 4x_Siberia | 103 | 59 | 6.13 | 0.786 | 0.761 |
| 4x_Europe | 30 | 29 | 4.78 | 0.635 | 0.589 |
| ***Mean 4X ±S.D.*** | | ***44.0 ±21.2*** | ***5.46 ±0.95*** | ***0.710 ±0.107*** | ***0.675 ±0.122*** |

**Table S7.** Comparison of within S-allele nucleotide diversity between diploid and tetraploid populations of *A. arenosa* and *A. lyrata*. Values of nucleotide diversity (**π**) and net nucleotide divergence between cytotypes are reported for each selected S-allele. The number of gene copies of each S-allele in each species and cytotype is given. S-alleles are grouped by dominance class.

| | *A. arenosa* | | | | | *A. lyrata* | | | | |
|---|---|---|---|---|---|---|---|---|---|---|
| | diploids | | tetraploids | | dipl-tetrapl | diploids | | tetraploids | | dipl-tetrapl |
| | number of gene copies | π | number of gene copies | π | net divergence | number of gene copies | π | number of gene copies | π | net divergence |
| Class I | | | | | | | | | | |
| H1001 | 15 | 0.008660 | 15 | 0.00786 | -0.000060 | 121 | 0.00196 | 54 | 0.00167 | 0.00015 |
| Class II | | | | | | | | | | |
| H2001 | 7 | 0.002563 | 10 | 0.00309 | -0.000027 | 2 | 0.00000 | 6 | 0.01987 | 0.00133 |
| H2003 | 5 | 0.002583 | 8 | 0.00195 | 0.000127 | 27 | 0.00044 | 2 | 0.00095 | 0.00000 |
| H2004 | 9 | 0.002467 | 9 | 0.00267 | 0.000468 | 21 | 0.00085 | 7 | 0.00080 | 0.00003 |
| H2006 | 7 | 0.001134 | 16 | 0.00274 | 0.000093 | 6 | 0.00051 | 3 | 0.01107 | -0.00184 |
| H2007 | 14 | 0.001901 | 17 | 0.00270 | 0.000863 | 13 | 0.00185 | 5 | 0.00515 | 0.00584 |
| H2008 | 5 | 0.002799 | 7 | 0.01060 | -0.000652 | 3 | 0.00104 | 9 | 0.00400 | 0.00151 |
| H2009 | 5 | 0.002131 | 12 | 0.00309 | -0.000026 | 15 | 0.00103 | 8 | 0.00161 | 0.00001 |
| Class III | | | | | | | | | | |
| H3001 | 5 | 0.006971 | 7 | 0.00385 | 0.002062 | 9 | 0.00031 | 2 | 0.00990 | -0.00006 |
| H3002 | 5 | 0.001091 | 6 | 0.00106 | 0.001489 | 27 | 0.00011 | 0 | / | / |
| H3003 | 7 | 0.000669 | 4 | 0.00182 | 0.002316 | 28 | -0.00004 | 5 | 0.00070 | 0.00065 |
| H3004 | 7 | 0.006972 | 5 | 0.00184 | 0.002374 | 11 | -0.00025 | 2 | 0.01483 | 0.00064 |
| H3022 | 4 | 0.001541 | 5 | 0.00000 | 0.000910 | 25 | 0.00070 | 2 | 0.00079 | 0.00060 |
| Class IV | | | | | | | | | | |
| H4003 | 6 | 0.002217 | 8 | 0.00049 | 0.000379 | 8 | 0.00242 | 0 | / | / |
| H4004 | 9 | 0.001504 | 6 | 0.00406 | 0.001133 | 19 | 0.00309 | 4 | 0.00403 | 0.00141 |
| H4010 | 6 | 0.001857 | 7 | 0.00225 | 0.000725 | 26 | 0.00612 | 8 | 0.01089 | -0.00013 |
| H4015 | 4 | 0.000254 | 9 | 0.00396 | 0.001223 | 4 | 0.00000 | 5 | 0.00279 | -0.00034 |
| H4023 | 7 | 0.003577 | 5 | 0.00063 | 0.001425 | 3 | 0.00615 | 2 | 0.00000 | 0.00747 |
| H4026 | 7 | 0.002381 | 4 | 0.00095 | 0.005912 | 3 | 0.00154 | 4 | 0.00146 | 0.00000 |
| H4027 | 6 | 0.005659 | 7 | 0.00107 | 0.000062 | 10 | 0.00221 | 0 | / | / |
| H4030 | 5 | 0.002077 | 6 | 0.00321 | -0.000221 | 5 | 0.00234 | 2 | 0.00369 | 0.00001 |
| H4034 | 5 | 0.002816 | 8 | 0.00217 | 0.000399 | 27 | 0.00321 | 5 | 0.00508 | -0.00028 |
| H4035 | 5 | 0.005054 | 5 | 0.00291 | 0.000826 | 11 | 0.00217 | 5 | 0.00128 | -0.00009 |
